# Age-dependent brain proteome remodeling links *Abca7* deficiency to insulin signaling and neuroinflammation in Alzheimer’s disease mice

**DOI:** 10.64898/2026.07.28.741021

**Authors:** Daniel Mittli, Jens Pahnke

## Abstract

The human *ABCA7* gene, which encodes ATP-binding cassette transporter A7 (ABCA7), is one of the strongest genetic risk factors for late-onset Alzheimer’s disease (AD), yet the molecular mechanisms linking ABCA7 deficiency to AD remain incompletely understood. Because impaired insulin signaling and neuroinflammation are increasingly recognized as key contributors to AD pathogenesis, and ABCA7 has been implicated in both metabolic and immune processes, we investigated whether ABCA7 deficiency is associated with alterations in these pathways using comparative brain proteomics. Whole-brain proteomes from wild-type, *Abca7* knockout, AD model (APP), and APP- *Abca7* knockout mice were analyzed at 50, 100, 150, and 200 days of age using label-free quantitative proteomics, followed by differential abundance analysis, gene ontology enrichment, subcellular localization analysis, protein-protein interaction network analysis, and comparison with human AD brain proteomic datasets. Comparative analyses identified age-dependent proteomic alterations associated with *Abca7* deficiency, amyloid-beta pathology, and their interaction. Across all genotype comparisons, recurrently altered proteins and functional interaction networks consistently converged on insulin receptor and PI3K/AKT signaling, MAPK signaling, immune and complement pathways, vesicle trafficking and acidification, and the ubiquitin-proteasome system. Subcellular enrichment analysis further indicated preferential involvement of membrane-associated proteins, consistent with the established role of ABCA7 in membrane lipid transport and trafficking. Comparison with human AD proteomic data demonstrated substantial overlap while also identifying potentially novel proteins associated with *Abca7*deficiency in our mouse models. Together, these findings provide a systems-level characterization of the brain proteomic consequences of *Abca7* deficiency and suggest that dysregulation of interconnected metabolic, immune, and vesicle-associated pathways may contribute to AD-related brain pathology.

## 1. Introduction

Dementia, or major neurocognitive disorder, is a progressive decline in cognitive function across one or more domains (e.g., memory, language, problem-solving, and general thinking abilities) that severely interferes with daily life and impairs independence (Jones et al., 2025). The global prevalence of dementia was estimated to be around 57 million individuals in 2021, and nearly 10 million new cases are expected to emerge every year (World Health Organization, 2025). Dementia can develop due to a range of neurodegenerative conditions, among which Alzheimer’s disease (AD) is the most common, accounting for 60-70% of all cases (Yoon et al., 2023). The currently dominant hypothesis of AD pathogenesis emphasizes the cerebral aggregation of extracellular amyloid-beta peptides (A*β*) into senile plaques and the intraneuronal accumulation of hyperphosphorylated tau protein forming neurofibrillary tangles. These pathological features are thought to drive synaptic and neuronal dysfunction via various molecular and cellular processes, ultimately leading to irreversible synapse loss and neuronal cell death responsible for brain atrophy and cognitive decline (Hampel et al., 2021; Mehta and Schneider, 2022). Despite intense research focusing on these A*β*and tau-driven neurobiological changes, the exact etiology and pathogenesis of AD remain incompletely understood and, consequently, there is still no effective causal therapy for the disease.

In addition to the A*β*and tau-related mechanisms, the pathophysiology of AD encompasses several synergistically interacting molecular and cellular processes that contribute to the impairment of brain tissue homeostasis and exacerbate neurodegenerative changes. The most important of these processes include brain vascular dysfunction, impaired ABC transporterand glymphaticmediated brain clearance, mitochondrial and metabolic disturbances, oxidative stress, neuroinflammation, and neurotransmitter imbalance (Butterfield and Halliwell, 2019; Dib et al., 2021; Górska et al., 2026; Mehta and Mehta, 2023; Santos-Garcia et al., 2025; Steffen et al., 2018). Among the metabolic components of AD, disturbances in cellular lipid and glucose metabolism appear to play a key role (Balakrishnan et al., 2026; Butterfield and Halliwell, 2019). This is supported by the lipid transport-related functions associated with two of the strongest genetic risk determinants for sporadic, late-onset AD, the *APOE ε*4 allele and loss-of-function variants in *ABCA7* (El Gaamouch et al., 2016; Steinberg et al., 2015; Strittmatter and Roses, 1995), as well as by the robust epidemiological and mechanistic association between AD and type 2 diabetes mellitus (T2DM) (Wei et al., 2021). Specifically, both the apolipoprotein E *ε*4 isoform (APOE *ε*4, encoded by *APOE*) and ATP-binding cassette transporter A7 (ABCA7, encoded by *ABCA7*) are involved in the transport, clearance, secretion, and overall homeostasis of brain lipids, particularly cholesterol and phospholipids (Blumenfeld et al., 2024; Dib et al., 2021; Lamartiniere et al., 2018). Additionally, APOE has been shown to interact with ABCA7 and to regulate its lipid transport activity (Fang et al., 2025), further highlighting the link between dysregulated brain lipid metabolism and AD pathogenesis. Genome-wide association studies indicate that loss-of-function variants in *ABCA7* show a strong association with increased AD risk (odds ratio *≈* 2); these disease-associated variants likely result in reduced ABCA7 protein levels, insufficient to carry out its physiological function (Duchateau et al., 2024; Hollingworth et al., 2011; Steinberg et al., 2015). In addition to these mechanistic insights, lipidomic analyses further support the role of brain lipid dysregulation in AD pathology (Jayaprakash et al., 2026).

Another major metabolic component of AD is associated with impairments in cerebral insulin signaling and glucose metabolism. Insulin resistance (IR) is a condition in which the ability of cells to respond to insulin is reduced due to disruption of intracellular insulin signaling. IR is a central mechanism in the pathogenesis of T2DM; however, its involvement in AD-related brain pathology has also been established (Vinuesa et al., 2021; Wei et al., 2021; Yoon et al., 2023). In addition to this shared molecular mechanism, epidemiological data further support a partial overlap in the pathobiology of AD and T2DM, which has led to the concept of “AD as type 3 diabetes”. For example, a high proportion of AD patients exhibit impaired glucose metabolism or T2DM, and T2DM is known to roughly double the risk of developing AD (Wei et al., 2021). Brain IR has been shown to exacerbate AD pathology via multiple neurobiological alterations. Notably, impaired neuronal glucose utilization and loss of the neurotrophic effects of insulin may lead to deficits in neuronal differentiation, dendritic growth, synaptic plasticity, and overall neuronal survival, thereby enhancing the neurodegenerative effects of A*β* and tau pathologies (Vinuesa et al., 2021). In addition, disrupted cerebral insulin signaling promotes oxidative stress and neuroinflammation, resulting in increased A*β* production, impaired A*β* clearance, and enhanced tau hyperphosphorylation (de la Monte, 2012; Wei et al., 2021). Brain IR and neuroinflammation appear to form a vicious cycle central to AD pathology, as a major mechanism of brain IR is the serine phosphorylation of insulin receptor substrate 1 (IRS-1) by stress kinases (e.g., JNK, IKK, PKR), which are preferentially activated by proinflammatory mediators, such as tumor necrosis factor-*α* (TNF-*α*). This inflammation-related serine phosphorylation of IRS-1 interferes with the physiological, tyrosine phosphorylation-dependent insulin signaling, a mechanism also described in T2DM (Ferreira et al., 2014; Vinuesa et al., 2021). Independent of metabolic disturbances, neuroinflammation represents a central pathological feature of AD, largely driven by A*β*and tau-induced glial reactivity (particularly in microglia and astrocytes), which promotes a proinflammatory tissue environment through the activation of inflammatory signaling pathways and the release of soluble mediators (e.g., cytokines, chemokines, and lipid mediators) (Ardura-Fabregat et al., 2017; Novoa et al., 2022). However, in addition to proinflammatory signaling, dysregulated de novo lipogenesis and associated lipid accumulation can also promote IR and further potentiate inflammatory processes in AD (Khan et al., 2023). As mentioned earlier, the primary function of ABCA7 is to mediate lipid efflux and trafficking, although its involvement in systemic immune processes (e.g., macrophage phagocytosis and natural killer T cell maturation), as well as in the glial regulation of AD-related neuroinflammation is also well established (Dib et al., 2021; Nowyhed et al., 2017; Santos-Garcia et al., 2025; Villa et al., 2024). These observations provide a strong mechanistic link between dysregulated lipid metabolism, brain IR, and neuroinflammation in AD.

Taken together, recent data suggest that ABCA7 is mechanistically linked to key metabolic and inflammatory processes underlying AD. Notably, the human *ABCA7* gene is one of the strongest genetic susceptibility loci for AD (Duchateau et al., 2024; Gialama et al., 2024); however, its precise contribution to disease pathogenesis remains incompletely understood. This raises the question of whether ABCA7 hypofunction may increase AD risk by enhancing cerebral IR and neuroinflammation, in addition to its involvement in lipid metabolism. Establishing ABCA7 as a modulator of both insulin signaling and immune homeostasis in the brain would position it as a promising target for therapeutic intervention and advance our understanding of AD biology. Accordingly, pharmacological modulation and structure-guided ligand discovery have been proposed as strategies for investigating ABCA transporters, including ABCA7, as potential diagnostic and therapeutic targets (Namasivayam et al., 2021; Pahnke et al., 2021). To address this, the primary aim of the present study was to identify insulin signaling- and neuroinflammation-related proteomic alterations induced by *Abca7* deficiency in the presence or absence of A*β* pathology. To this end, we performed proteomic analyses of whole-brain tissue from wild-type control mice (C57BL/6J, referred to as B6), *Abca7* knockout mice (hA7ko), AD model mice (APP), and AD model mice with *Abca7* deficiency (APP-hA7ko) across four age groups (50, 100, 150, and 200 days). To systematically dissect the individual and combined effects of *Abca7* deficiency and A*β* pathology, we adopted a comparative brain proteomics strategy based on four predefined genotype comparisons. First, B6 versus hA7ko defined the baseline proteomic consequences of *Abca7* deficiency in a non-amyloidogenic context, highlighting proteomic changes that may reflect increased vulnerability prior to overt AD pathology. Second, hA7ko versus APP-hA7ko assessed how A*β* pathology manifests in the presence of *Abca7* deficiency, thereby revealing how pre-existing *Abca7* loss modulates proteomic responses to A*β* burden, particularly in insulin signaling and neuroinflammatory pathways. Third, APP versus APP-hA7ko isolated the contribution of *Abca7* deficiency to A*β*-driven pathology, addressing whether loss of *Abca7* function modifies disease-related processes in an established AD model. In addition to the above comparisons, B6 versus APP-hA7ko provided an integrated view of the combined effects of A*β* pathology and *Abca7* deficiency, capturing their joint impact on brain proteomic alterations, although their individual contributions cannot be distinguished in this comparison.

The present bioinformatic analysis demonstrates that *Abca7* deficiency is associated with widespread remodeling of brain proteomic networks linked to insulin signaling and neuroinflammatory processes, both in the absence and presence of A*β* pathology. Across the four genotype comparisons, recurrent alterations consistently converged on a limited number of interconnected molecular systems, including insulin receptor and PI3K/AKT signaling, MAPK and NF-*κ*B signaling, vesicle trafficking and acidification, and the ubiquitin-proteasome system. The observed enrichment of membrane-associated proteins further supports a central role of ABCA7 in plasma membrane-associated signaling and trafficking processes, whereas comparison with human AD brain proteomic datasets revealed substantial overlap while also identifying novel candidate proteins associated with *Abca7* deficiency. Together, these findings provide a systems-level framework linking *Abca7* hypofunction to metabolic and inflammatory mechanisms relevant to AD pathogenesis and identify molecular pathways that may represent promising targets for future mechanistic and therapeutic studies.

## 2. Materials and methods

In the present study, we performed a bioinformatic analysis of our previously generated proteomic dataset, which was deposited in PRIDE – Proteomics Identification Database (project identifier: PXD053250, project title: “ABCA7 transporter functional role on Alzheimer’s disease onset and progression”) (https://www.ebi.ac.uk/pride/archive/projects/PXD053250). Therefore, we briefly introduce the investigated mouse lines, and summarize the key aspects of tissue sample preparation, the liquid chromatography-tandem mass spectrometry (LC-MS/MS) analysis of the samples (data acquisition), and the processing of raw LC-MS/MS data (including protein identification, inference, and quantification), referring the reader to our previous publications (Górska et al., 2026; Santos-Garcia et al., 2025). We then provide a more detailed description of the downstream bioinformatic investigation of the proteomic dataset, which underlies the main analyses and results presented in this paper.

Gene and protein nomenclature follows HGNC and Mouse Genome Informatics conventions. Human gene symbols are italicized and uppercase (e.g., *ABCA7*); mouse gene and allele symbols are italicized with an initial uppercase letter followed by lowercase letters (e.g., *Abca7*); and protein symbols are uppercase and non-italicized (e.g., ABCA7 and NFKBIB).

### 2.1. Animals and sample preparation

To investigate the proteome-level role of *Abca7* in AD-related alterations in brain insulin signaling and inflammatory processes, we analyzed whole-brain proteomes from the following four mouse (Mus musculus) lines: (i) wild-type C57BL/6J mice (hereafter referred to as B6); (ii) mice with constitutive *Abca7* knockout, in which the murine locus was replaced by the human *ABCA7* sequence and subsequently inactivated (allele: *Abca7* ^tm1.1(ABCA7)Pahnk^; hereafter referred to as hA7ko); (iii) APPPS1-21 mice (B6.Cg-Tg(Thy1-APPSw,Thy1-PSEN1L166P)21JkcrPahnk) used for AD modelling (hereafter referred to as APP); and (iv) APP mice crossed with hA7ko mice, showing both progressive A*β* pathology and *Abca7* deficiency (hereafter referred to as APP-hA7ko). The *Abca7* knockout allele was homozygous (*Abca7* ^-/-^), whereas the APPPS1-21 transgene was hemizygous (+/Tg). APPPS1-21 hemizygous, *Abca7* -heterozygous males were crossed with *Abca7* -heterozygous females to generate experimental animals, which were allocated to the experimental groups according to genotype. The detailed description of the mouse models and breeding scheme can be found in our previous publication (Santos-Garcia et al., 2025). The animals were housed at the Department of Comparative Medicine, Oslo University Hospital (Oslo, Norway) under standard laboratory conditions (22 *±* 1 *^◦^*C, 62 *±* 5% relative humidity, 15 air changes per hour, 12-hour light/12-hour dark cycle) in Eurostandard type III cages filled with standard bedding material and provided with additional environmental enrichment. The animals were grouped into cohorts of up to 8 individuals per cage. Mice had ad libitum access to food and water. Health monitoring was performed three times per year in accordance with FELASA guidelines. Housing and procedures were conducted following the guidelines for animal experimentation of the European Union and Norwegian national legislation. The animal study protocol and breeding were approved by the KPM Forsøksdyrutvalget and the Norwegian Food Safety Authority (Mattilsynet; IV2-2022, FOTS 20540). In the present study, we analyzed whole-brain proteomic data from animals of four different ages (50, 100, 150, and 200 days) and both sexes, which resulted in 16 different experimental groups. The total number of animals included in the analyses was n = 83, and the composition of the final experimental groups was as follows: B6 50 days, n = 5 (2 females, 3 males); B6 100 days, n = 6 (4 females, 2 males); B6 150 days, n = 5 (3 females, 2 males); B6 200 days, n = 6 (4 females, 2 males); hA7ko 50 days, n = 5 (2 females, 3 males); hA7ko 100 days, n = 5 (2 females, 3 males); hA7ko 150 days, n = 5 (2 females, 3 males); hA7ko 200 days, n = 6 (2 females, 4 males); APP 50 days, n = 5 (2 females, 3 males); APP 100 days, n = 5 (2 females, 3 males); APP 150 days, n = 5 (2 females, 3 males); APP 200 days, n = 5 (2 females, 3 males); APP-hA7ko 50 days, n = 5 (2 females, 3 males); APP-hA7ko 100 days, n = 5 (2 females, 3 males); APP-hA7ko 150 days, n = 5 (3 females, 2 males); APP-hA7ko 200 days, n = 5 (2 females, 3 males).

Whole-brain tissue samples were prepared as described earlier (Górska et al., 2026; Santos-Garcia et al., 2025). Briefly, mice were euthanized by cervical dislocation, transcardially perfused with icecold PBS, and brains were removed and divided into two hemispheres. One of the hemispheres was snap-frozen in liquid nitrogen and stored at -80 *^◦^*C until protein extraction. Thus, brain tissue samples from individual animals served as biological replicates for downstream proteomic analyses. Whole hemispheres were homogenized, and equal amounts of brain tissue homogenate (approximately 20 *µ*g total protein) were precipitated on amine beads according to a published protocol (Batth et al., 2019). The precipitated proteins were dissolved in 50 mM ammonium bicarbonate, reduced, alkylated and digested with trypsin (1:50 enzyme to protein ratio, Promega, Madison, WI, USA) at 37 *^◦^*C overnight. Digested peptides were then acidified and desalted on Evotips according to the manufacturer’s protocol (Evosep Biosystems, Odense, Denmark).

### 2.2. LC-MS/MS-based quantitative proteomics

To perform the quantitative proteomic analysis of brain tissue samples, a standard, mass spectrometry (MS)-based bottom-up proteomics workflow was applied as described earlier (Górska et al., 2026). Briefly, liquid chromatography-tandem mass spectrometry (LC-MS/MS) analysis was performed using an Evosep one LC system (Evosep Biosystems) coupled to a timsTOF fleX mass spectrometer, using a CaptiveSpray nano electrospray ion source (Bruker Corporation, Billerica, MA, USA). 200 ng of digested peptides were loaded onto a capillary C18 column (150 mm *×* 75 *µ*m, 1.7 *µ*m particle size, 120 Å pore size; IonOpticks, Fitzroy, VIC, Australia). Peptides were separated at 50 *^◦^*C using a standard 40 sample/day method from Evosep. The mass spectrometer was operated in dia-PASEF mode (Skowronek and Meier, 2022). Mass spectra for MS were recorded between m/z 100-1700, and ion mobility resolution was set to 0.85-1.30 V*·*s*·*cm-1 over a ramp time of 100 ms. The MS/MS mass range was set to m/z 475-1000, and ion-mobility resolution to 0.85-1.27 V*·*s*·*cm-1 to exclude singly charged ions. The estimated cycle time was 0.95 s with eight cycles using data-independent acquisition (DIA) windows of 25 Da. Collisional energy was ramped from 20 eV at 0.60 V*·*s*·*cm-1 to 59 eV at 1.60 V*·*s*·*cm-1. All samples were analyzed within a single MS run to avoid inter-run variability. Raw data files from LC-MS/MS analyses were submitted to DIA data processing (DIA-NN v1.8.1) for protein identification and label-free quantification (LFQ) using the library-free function and the mobility module for dia-PASEF analysis (Demichev et al., 2020, 2022). In silico digestion of the UniProtKB mouse (Mus musculus) database (The UniProt Consortium., 2024) was performed to generate a spectral library. Carbamidomethyl was set as a fixed modification. Trypsin digestion without proline restriction enzyme option was used, with one allowed miscleavage, and peptide length range was set to 7-30 amino acids. The mass accuracy was set to 15 ppm, and allowed precursor false discovery rate (FDR) was 0.01 (1%).

LFQ intensities (relative protein abundances) obtained from DIA data processing were log2transformed prior to statistical group comparisons to stabilize variance and approximate a normal distribution (Aguilan et al., 2020; Zhao et al., 2024). Missing values (non-detection) were substituted with a constant integer number (8.00) corresponding to the lower intensity range of the dataset, estimating signals below the detection limit (left-censored data imputation) (Webb-Robertson et al., 2015). For each age group (50, 100, 150, and 200 days), predefined pairwise comparisons between genotypes (B6 vs. hA7ko, hA7ko vs. APP-hA7ko, APP vs. APP-hA7ko, and B6 vs. APP-hA7ko) were analyzed using a two-tailed Welch’s t-test. For each group comparison, the fold change (FC) in protein abundance between the two conditions was calculated using the group means of log2transformed LFQ intensities (log2(FC) = mean LFQ intensity 1 – mean LFQ intensity 2). Data preprocessing and group comparisons were performed using Perseus v2.0.11 (Max Planck Institute of Biochemistry, Martinsried, Germany) (Tyanova et al., 2016) and Microsoft Excel 365 (Microsoft Corporation, Redmond, WA, USA).

### 2.3. Bioinformatic and statistical analyses of brain proteomic data

#### 2.3.1. Quality control of quantitative proteomic data

Quality control analyses of log2-transformed LFQ intensities were performed to assess reproducibility, technical noise, and overall data quality based on established practices in quantitative omics approaches (Albrecht et al., 2010; De Hertogh et al., 2010; Schessner et al., 2022; Tsantilas et al., 2024). First, we assessed the global missingness across the entire dataset (all samples *×* all proteins), which was defined as the percentage of missing values relative to the total number of values in the dataset. Per-sample missingness was calculated analogously, considering only values within each individual sample. Biological and technical reproducibility were assessed using pairwise correlation plots together with Pearson correlation calculations, protein-wise standard deviation (SD) histograms, mean-SD plots, and MA plots. To evaluate the reproducibility of protein quantification among reliably detected proteins, these analyses were restricted to proteins quantified in all samples (i.e., after applying a 100% completeness filter). Data from two randomly selected animals per experimental group (replicate pairs) were used to generate pairwise correlation plots and calculate Pearson correlations to assess similarity between replicate samples. To visualize the spread of variability (SD), histograms of protein-wise SDs were generated for each experimental group, and the median and interquartile range (IQR) were calculated. To further assess data distribution quality and consistency between replicates, mean-SD plots were prepared for each experimental group, showing the empirical relationship between protein intensity (average abundance) and variability (SD). Similarly, MA plots were generated for each experimental group using randomly selected replicate pairs to identify intensity-dependent bias and potential normalization issues. In these plots, M represents the difference between the two samples (M = LFQ intensity 1 – LFQ intensity 2), while A represents the average abundance of the two samples (A = (LFQ intensity 1 + LFQ intensity 2) / 2). Quality control calculations were performed using Microsoft Excel 365 (Microsoft Corporation, Redmond, WA, USA) and Origin 2026 (OriginLab Corporation, Northampton, MA, USA), and plots were prepared in Origin 2026.

#### 2.3.2. Compilation of reference protein lists

The aim of the present study was to identify *Abca7* deficiency-induced brain proteomic changes related to insulin signaling and neuroinflammation, with or without A*β* pathology; thus, we focused on proteins taking part in inflammatory processes and insulin signaling. To obtain a comprehensive reference list of these proteins, we performed a database search using the expert-curated Reactome pathway database (https://reactome.org/) (Ragueneau et al., 2026) and the community-curated WikiPathways pathway resource (https://www.wikipathways.org/) (Agrawal et al., 2023), and downloaded the protein lists of those mouse (Mus musculus) pathways that can be associated with insulin signaling and neuroinflammatory processes. The following molecular pathways were downloaded from the two databases (pathway name and pathway identifier): Reactome – Signaling by Insulin receptor (R-MMU-74752), Chemokine receptors bind chemokines (R-MMU-380108), Eicosanoid ligand-binding receptors (R-MMU-391903), Interferon Signaling (R-MMU-913531), Nucleotide-binding domain, leucine rich repeat containing receptor (NLR) signaling pathways (R-MMU-168643), ROS and RNS production in phagocytes (R-MMU-1222556), Signaling by CSF3 (G-CSF) (R-MMU-9674555), Signaling by Interleukins (R-MMU-449147.1), Signaling by the B Cell Receptor (BCR) (R-MMU-983705), TCR signaling (R-MMU-202403), TNFR2 non-canonical NF-kB pathway (R-MMU-5668541), Toll-like Receptor Cascades (R-MMU-168898); WikiPathways – Insulin signaling (WP65), B cell receptor signaling pathway (WP274), Chemokine signaling pathway (WP2292), Cytokines and inflammatory response (WP222), Eicosanoid synthesis (WP318), IL-1 signaling pathway (WP37), IL-2 signaling pathway (WP450), IL-3 signaling pathway (WP373), IL-4 signaling pathway (WP93), IL-5 signaling pathway (WP151), IL-6 signaling pathway (WP387), IL-7 signaling pathway (WP297), IL-9 signaling pathway (WP10), Inflammatory response pathway (WP458), Macrophage markers (WP2271), Microglia pathogen phagocytosis pathway (WP3626), Nod-like receptor (NLR) signaling pathway (WP1256), Prostaglandin synthesis and regulation (WP374), T cell receptor signaling pathway (WP480), TNF-alpha NF-kB signaling pathway (WP246), Toll-like receptor signaling (WP88). Because the protein lists for these signaling pathways overlapped substantially, we merged them and removed duplicates to obtain unified, non-overlapping protein sets for insulin signaling (217 proteins) and neuroinflammatory processes (1230 proteins) (Supplementary Material, Table S1). After quality control of the complete proteomic dataset (see previous section), we focused the downstream bioinformatic analyses on these selected proteins.

#### 2.3.3. Identification and exploratory analyses of differentially abundant proteins

To identify differentially abundant proteins (DAPs), volcano plots were prepared for each comparison using t-test p-values (-log10(p-value), statistical significance) and FC data (log2(FC), magnitude of change). DAPs were defined by a square cutoff method, where p-value threshold was 0.05 (-log10(p-value) *≥* 1.30) and FC threshold was *±*1.5 (|log2(FC)| *≥* 0.585). Volcano plots were generated using the VolcaNoseR web app (https://huygens.science.uva.nl/VolcaNoseR/) (Goedhart and Luijsterburg, 2020). Unsupervised exploratory analysis of proteins identified as differentially abundant in at least one comparison was performed using principal component analysis (PCA) plots and heatmaps with marginal dendrograms (showing hierarchical clustering), which are commonly applied in the evaluation of multi-condition proteomic experiments (Schessner et al., 2022). PCA plots and heatmaps (combined with hierarchical clustering) were created using the ClustVis web tool (https://biit.cs.ut.ee/clustvis/) (Metsalu and Vilo, 2015). For PCA, row-wise unit variance scaling was applied prior to analysis, and singular value decomposition (SVD) was used to compute principal components. Scree plots showing the variance explained by each principal component were also prepared. For heatmap visualization, rows were mean-centered and scaled to unit variance. Hierarchical clustering of samples (i.e., animals) was performed to assess whether proteomic data-based clustering reflected the genotype-defined experimental groups. Pairwise distances were calculated using Pearson correlation, and average linkage (average distance of all possible pairs) was used as the linkage method.

#### 2.3.4. Functional annotation, gene ontology enrichment, and subcellular localization analyses

We started the functional annotation of DAPs by manually reviewing the top 20 hits (or fewer, if < 20 were identified) from each comparison using their corresponding reviewed (Swiss-Prot) UniProtKB entries (https://www.uniprot.org/) (The UniProt Consortium., 2024). These top hits were subsequently assigned to functional categories based on their primary biological function and involvement in molecular pathways. Gene ontology (GO) enrichment analysis (pathway database: GO biological process) of the complete lists of DAPs from each comparison was performed using ShinyGO v0.85.1 (https://bioinformatics.sdstate.edu/go/) (Ge et al., 2019), provided that at least 10 DAPs were identified in a given comparison. For each enrichment analysis, the background set consisted of all quantified proteins in our dataset (7636 proteins), rather than the entire mouse genome, and default settings of ShinyGO were applied. P-values were calculated using the hypergeometric test, and FDR was controlled using the Benjamini-Hochberg procedure to correct for multiple testing. Fold enrichment (indicating effect size) was defined as the percentage of proteins in the query list associated with a given pathway divided by the corresponding percentage in the background protein set. To assess which subcellular compartments the detected DAPs are associated with, we performed subcellular compartment enrichment analysis for each comparison using SubcellulaRVis (https://shiny.its.manchester.ac.uk/subcellularvis/) (Watson et al., 2022). As for the GO enrichment analysis, the complete list of quantified proteins was used as the background set.

#### 2.3.5. Protein-protein interaction network analyses and comparison with human AD proteomic datasets

To perform the in silico investigation of functional protein-protein interactions (PPIs) among the DAPs, PPIs were mapped using STRING v12.0 (https://string-db.org/) (Szklarczyk et al., 2023), and functional PPI networks were visualized using Cytoscape v3.10.4 (https://cytoscape.org/) (Shannon et al., 2003). PPI mapping was performed for each comparison, provided that at least 10 DAPs were quantified. To identify potential functional interactions, the full mouse (Mus musculus) STRING network was queried, and undirected edges were introduced between proteins based on the following interaction sources: experiments, databases, neighborhood, gene fusion, co-occurrence, co-expression (textmining excluded). For PPI network construction, only query proteins and interactions with medium confidence (minimum required interaction score: 0.400) were considered. PPI data were then exported and visualized in Cytoscape, where node size represented statistical significance (-log10(p-value)) and node fill color represented the magnitude of change (log2(FC)) for each DAP within a given comparison. Clustering of the resulting PPI networks was performed using the clusterMaker app in Cytoscape with the community analysis-based GLay clustering algorithm (Su et al., 2010). To assess the primary biological interpretation of each cluster, GO enrichment analysis was performed, and the PPI network figures were annotated with the top three enriched GO terms for each cluster. In an additional set of PPI networks, we specifically highlighted comparison-specific DAPs, defined as DAPs identified in a given comparison only and not overlapping with DAPs from any other comparison at the same age. The functional interpretation of these comparison-specific DAPs was likewise assessed using GO biological process enrichment analysis. Finally, to evaluate the translational relevance of our proteomic findings, we compared the DAP lists obtained from all 16 genotype-age comparisons with NeuroPro, a searchable database integrating more than 8000 AD-related human proteins compiled from 55 published studies investigating proteomic changes in the human brain (https://neuropro.biomedical.hosting/) (Askenazi et al., 2023). For each comparison, we determined the proportions of DAPs represented in the NeuroPro database and those not represented (novel DAPs).

## 3. Results

The following sections present the results according to the analytical workflow applied to the quantitative proteomic dataset. First, the quality of the quantitative proteomic data and an overview of DAPs across the predefined genotype comparisons are presented. The identified proteomic alterations are then systematically characterized using exploratory, functional, and network-based bioinformatic analyses, followed by the description of representative protein abundance changes and comparison with human AD proteomic datasets to assess the translational relevance of the findings.

### 3.1. Quantitative proteomics data show low missingness, high reproducibility, and no major batch effects

In total, 7636 proteins were quantified in whole-brain tissue samples from 83 mice across four genotypes (B6, hA7ko, APP, APP-hA7ko) and four ages (50, 100, 150, 200 days), yielding 16 experimental groups. Quality control analyses were performed within groups to assess data completeness, variance structure, and reproducibility. Global and per-sample missingness, average LFQ intensities, pairwise correlation plots, protein-wise SD histograms, mean-SD plots, and MA plots were generated based on protein LFQ values (Supplementary Material, Figs. S1–S6). The complete dataset used for missingness assessment also included additional samples not relevant for downstream analyses in this study (total n = 128). In this 128 *×* 7636 matrix, global missingness was 8.70%, and per-sample missingness ranged from 4.54% to 21.32% (Supplementary Material, Fig. S1). Average log2 LFQ intensity levels and their spread (mean *±* SD) were very similar across the investigated genotype-age groups (lowest in B6 200 days: 13.23 *±* 2.67; highest in APP 200 days: 13.46 *±* 2.51), indicating the absence of major batch effects or normalization artifacts (Supplementary Material, Fig. S2). A total of 4774 proteins (62.52%) were detected in all samples (i.e., these proteins showed 100% completeness). To avoid bias from low-abundance or stochastically detected proteins, subsequent quality control analyses were restricted to these 4774 proteins. Pairwise Pearson correlations of protein LFQ intensities between randomly selected biological replicates were consistently high (lowest in hA7ko 50 days: r = 0.9260; highest in APP-hA7ko 50 days: r = 0.9953), supporting good technical and biological reproducibility (Supplementary Material, Fig. S3). Protein-wise SD histograms of LFQ data showed low within-group variability with narrow distributions (lowest in APP-hA7ko 50 days: median SD = 0.12, IQR = 0.09–0.17; highest in hA7ko 100 days: median SD = 0.32, IQR = 0.22–0.45), further indicating high data quality (Supplementary Material, Fig. S4). Consistent patterns in mean-SD plots and MA plots across genotype-age groups, without evident intensity-dependent bias, also support the robustness of the quantitative proteomic data (Supplementary Material, Figs. S5 and S6).

### 3.2. Differentially abundant proteins and unsupervised exploratory analysis demonstrate early Abca7 deficiency effects and progressive genotype separation

The downstream bioinformatic analysis of the quantitative proteomic data focused on reference sets of 217 insulin signalingand 1230 neuroinflammation-related proteins. Of these, 136 (62.67%) and 568 (46.18%) proteins, respectively, were identified in our dataset. Seventy-five proteins were present in both sets; thus, the final non-overlapping list of identified insulin signalingand neuroinflammation-related proteins comprised 629 proteins (Supplementary Material, Tables S1 and S2).

Fig. 1 summarizes the overall study design and the distribution of DAPs across genotypes and ages, showing the experimental groups and predefined pairwise comparisons (Fig. 1A), the total number of DAPs per comparison and age (Fig. 1B), and the number of comparison-specific and shared DAPs for each age and comparison (Fig. 1C). In B6 vs. hA7ko, a relatively high number of DAPs were detected at 50 and 100 days (138 and 158 DAPs, respectively), which decreased at 150 days (106 DAPs) and increased again at 200 days (130 DAPs). The hA7ko vs. APPhA7ko comparison showed a similar pattern, with many DAPs at 50 and 100 days (156 and 174, respectively), followed by a marked decrease at 150 days (28 DAPs) and a subsequent increase at 200 days (75 DAPs). In contrast, APP vs. APP-hA7ko and B6 vs. APP-hA7ko displayed a clear age-dependent increase in DAP numbers. In APP vs. APP-hA7ko, we identified 14, 10, 60, and 129 DAPs at 50, 100, 150, and 200 days, respectively. In B6 vs. APP-hA7ko, the corresponding numbers were 18, 61, 92, and 178 DAPs. Except for a few cases, the numbers of upand down-regulated DAPs were broadly similar within each comparison and age (Fig. 1B).

**Figure 1:**
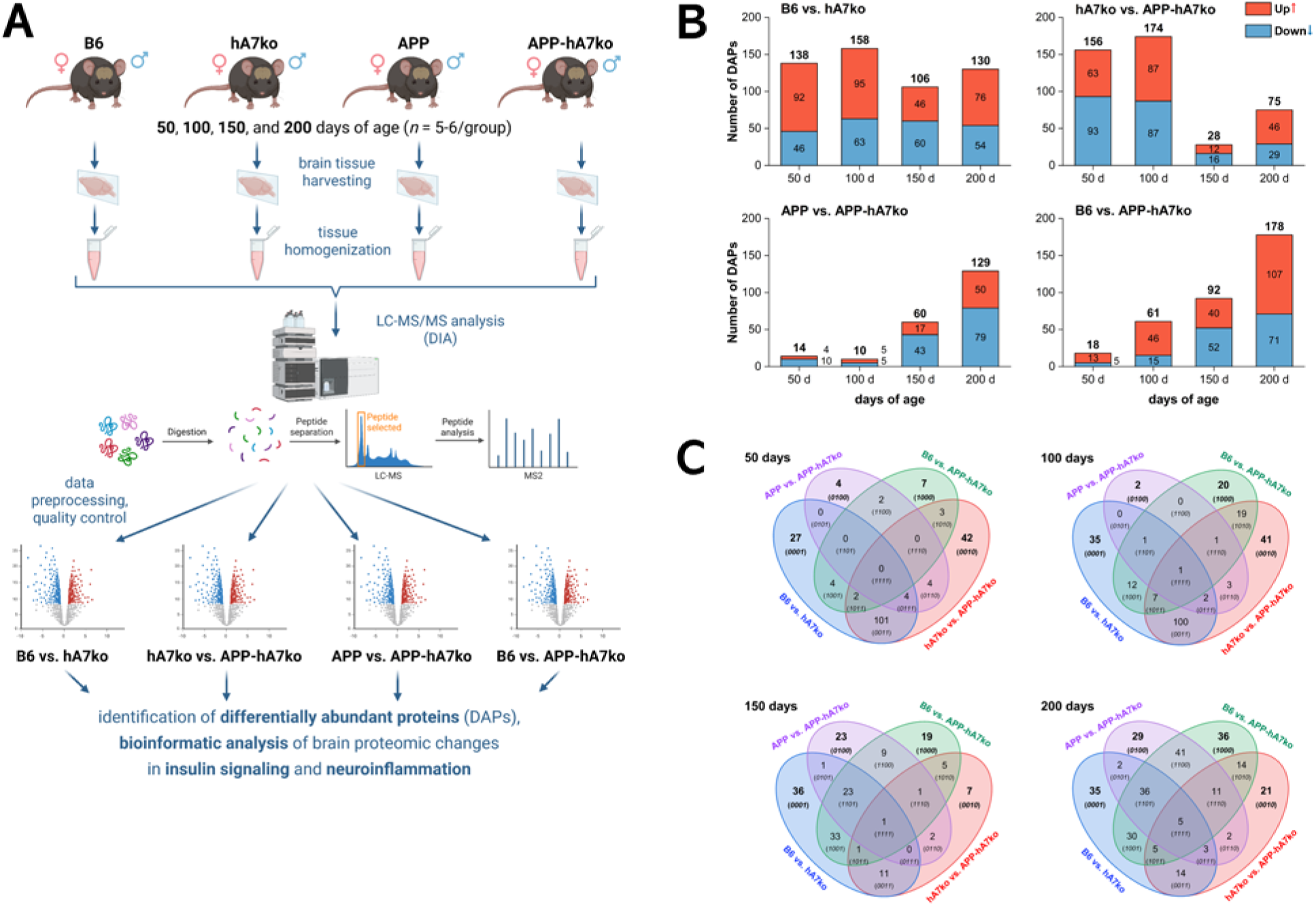
Study design and overview of differentially abundant proteins across comparisons and ages. (A) Schematic of the experimental groups (B6, hA7ko, APP, APP-hA7ko), age points (50, 100, 150, 200 days), and predefined pairwise comparisons performed on quantitative proteomic data. (B) Total numbers of DAPs per comparison and age, with stacked bars indicating up(red) and down-regulated (blue) proteins (DAPs were defined as p-value *≤* 0.05 and |FC| *≥* 1.5). (C) Venn diagrams for each age showing the number of comparison-specific and shared DAPs among the four comparisons (B6 vs. hA7ko: blue; hA7ko vs. APP-hA7ko: red; APP vs. APP-hA7ko: purple; B6 vs. APP-hA7ko: green). The numbers of DAPs indicate that *Abca7* deficiency and A*β* pathology in the context of *Abca7* deficiency induce early proteomic alterations in brain insulin signaling and neuroinflammatory processes, while the effect of *Abca7* loss on A*β*-driven pathology and the combined APP-hA7ko phenotype becomes pronounced at later ages.

The overlap analysis (Fig. 1C) revealed that at 50 days no DAPs were common to all four comparisons, whereas 27, 42, 4, and 7 DAPs were specific to B6 vs. hA7ko, hA7ko vs. APP-hA7ko, APP vs. APP-hA7ko, and B6 vs. APP-hA7ko, respectively. B6 vs. hA7ko and hA7ko vs. APPhA7ko shared many DAPs (101) at this age. At 100 days, one DAP (UBE2D3) was shared by all four comparisons, and 35, 41, 2, and 20 DAPs were comparison-specific in B6 vs. hA7ko, hA7ko vs. APP-hA7ko, APP vs. APP-hA7ko, and B6 vs. APP-hA7ko, respectively; again, B6 vs. hA7ko and hA7ko vs. APP-hA7ko shared many DAPs (100). At 150 days, one DAP (PEG3) was common to all four comparisons, whereas 36, 7, 23, and 19 DAPs were specific to B6 vs. hA7ko, hA7ko vs. APP-hA7ko, APP vs. APP-hA7ko, and B6 vs. APP-hA7ko, respectively. At 200 days, five DAPs (FCER1G, PPP1R13L, YES1, ELOC, UBE2V1) were shared by all four comparisons, and 35, 21, 29, and 36 DAPs were comparison-specific in B6 vs. hA7ko, hA7ko vs. APP-hA7ko, APP vs. APP-hA7ko, and B6 vs. APP-hA7ko, respectively.

Overall, these patterns indicate that *Abca7* deficiency (B6 vs. hA7ko) and A*β* pathology in the context of *Abca7* deficiency (hA7ko vs. APP-hA7ko) are associated with substantial early (50–100 days) proteomic alterations in brain insulin signaling and neuroinflammatory processes, whereas the impact of *Abca7* loss on A*β*-driven pathology (APP vs. APP-hA7ko) and the combined APPhA7ko phenotype (B6 vs. APP-hA7ko) becomes more prominent at later ages (150–200 days). The complete lists of DAPs and comparison-specific DAPs for all comparisons and ages are provided in the Supplementary Material (Tables S3 and S4).

The unsupervised exploratory analysis of quantitative proteomic data was performed by PCA and hierarchical clustering of LFQ values of DAPs identified at each age (Fig. 2). Scree plots of the PCA (Supplementary Material, Fig. S7) showed that the first few principal components captured a substantial proportion of the variance at all ages, justifying the use of the first two components for visualization. For each PCA, 95% prediction ellipses were plotted around genotype groups; thus, non-overlapping ellipses indicate distinct clustering in PCA space.

**Figure 2:**
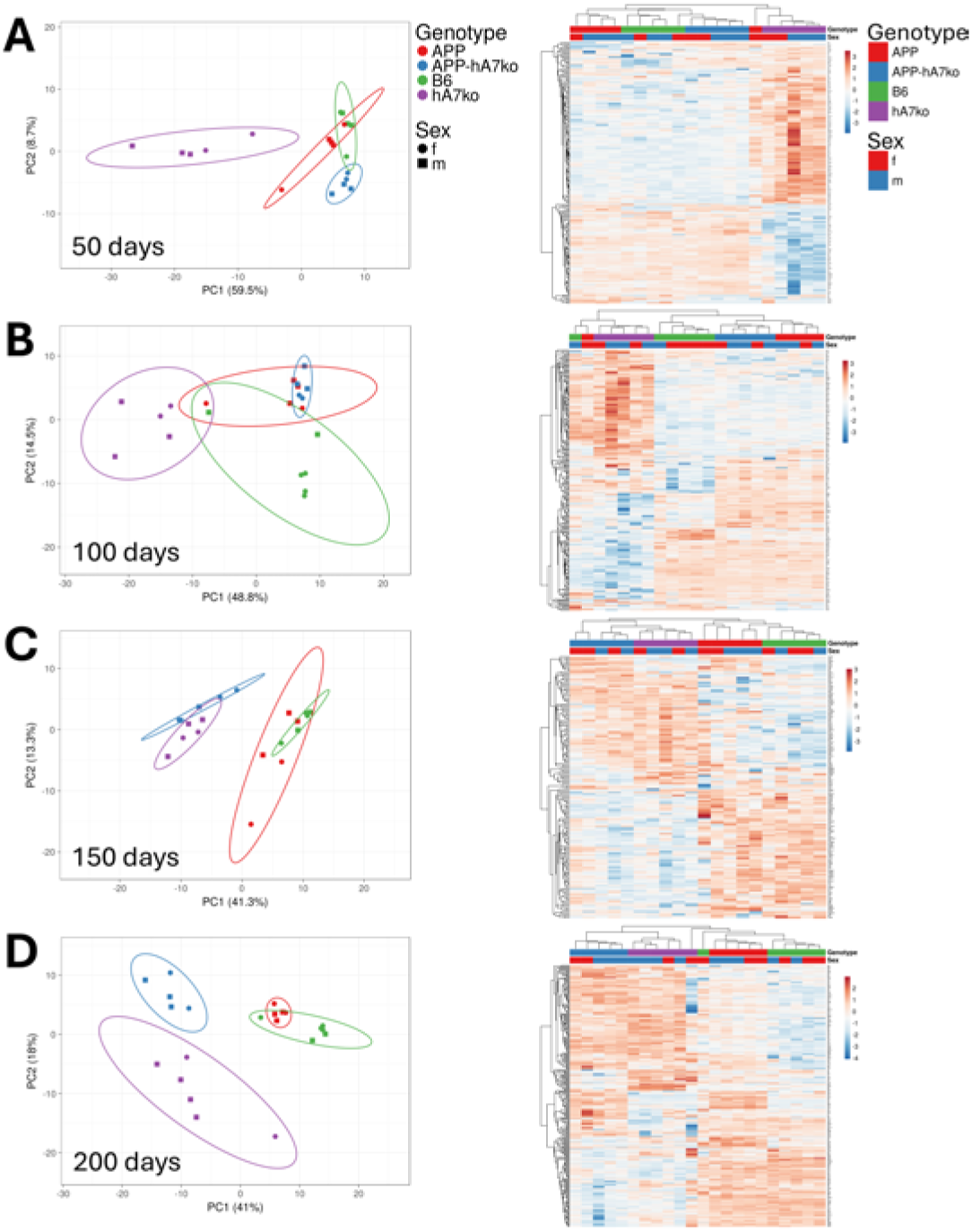
Principal component analysis and hierarchical clustering of differentially abundant proteins by genotype at each age. For each age (A: 50 days; B: 100 days; C: 150 days; D: 200 days), PCA was performed on LFQ values of DAPs, with samples colored by genotype (B6: green; hA7ko: purple; APP: red; APP-hA7ko: blue) and shaped by sex (females: circles; males: squares). 95% prediction ellipses are shown to indicate group dispersion. Corresponding heatmaps show hierarchical clustering of the same DAPs, with samples annotated by genotype (same color code as for PCA) and sex (females: red; males: blue). PCA and clustering suggest that genotype-based separation becomes more pronounced with age. hA7ko animals are already clearly separated at 50 days, and *Abca7* -deficient (hA7ko, APP-hA7ko) and *Abca7* -expressing (B6, APP) animals form distinct clusters at later ages, most evidently at 150 days. APP-hA7ko samples appear increasingly distinct from both APP and hA7ko, particularly at 200 days.

At 50 days (Fig. 2A), PCA clearly separated hA7ko animals from the other genotypes, which clustered more closely together. Hierarchical clustering segregated animals largely by genotype, with only one B6 and one APP animal assigned to clusters not matching their genotype. At 100 days (Fig. 2B), PCA did not yet show a complete separation of genotypes, although a trend toward genotype-dependent separation was evident. Hierarchical clustering grouped hA7ko and APP-hA7ko animals fully according to genotype, while again one B6 and one APP animal were misclustered. At 150 days (Fig. 2C), PCA clearly separated *Abca7* -deficient animals (hA7ko and APP-hA7ko) from *Abca7* -expressing animals (B6 and APP). This pattern was recapitulated by hierarchical clustering, which first split the samples into two major clusters according to *Abca7* status. Within these, both B6 and hA7ko animals formed well-defined subclusters, whereas two APP and two APP-hA7ko animals were not perfectly grouped by genotype. At 200 days (Fig. 2D), three groups occupied clearly distinct regions of PCA space: B6 and APP together (although they showed an apparent tendency for separation), hA7ko, and APP-hA7ko. In the corresponding hierarchical clustering, APP and APP-hA7ko animals each formed distinct clusters, while one B6 and one hA7ko animal were assigned outside their respective genotype clusters.

Overall, the unsupervised exploratory analysis of differentially abundant insulin signalingand neuroinflammation-related proteins suggests that genotype-based separation became more pronounced with age. APP-hA7ko samples appeared increasingly distinct from both APP and hA7ko, particularly at 200 days, whereas hA7ko samples already showed clear separation at 50 days. It is also noteworthy that APP animals were clearly separated from both hA7ko and APP-hA7ko animals in most cases. No obvious clustering by sex was observed at any age, which may reflect the relatively small number of animals per sex group.

### 3.3. Abca7 deficiency is associated with age-dependent alterations in membrane-associated immune and insulin signaling protein networks

*Abca7* deficiency (assessed by the B6 vs. hA7ko comparison) induced widespread proteomic changes in brain insulin signaling- and neuroinflammation-related processes already at a young age. At 50 days, among 138 DAPs, 92 showed increased and 46 showed decreased LFQ abundances in hA7ko compared with B6 (Fig. 3A). The most strongly upregulated proteins included NFKBIB (an inhibitor of NF-*κ*B signaling), RPS6KB1 (a serine/threonine kinase that regulates translation and cell growth), RELA (the p65 subunit of the NF-*κ*B transcription factor complex), SMARCC2 (a core subunit of the neuron-specific SWI/SNF chromatin-remodeling complex), and GYG1 (a priming enzyme in glycogen biosynthesis). Among the most strongly downregulated proteins were ADCY5 (a membrane-bound adenylate cyclase generating cAMP), GNG2 (a gamma subunit of heterotrimeric G proteins), ATP2B4 (a plasma membrane calcium ATPase), GLG1 (a Golgi-resident glycoprotein involved in protein trafficking), and GNAI3 (an inhibitory alpha subunit of heterotrimeric G proteins). The complete list of DAPs, including their corresponding FC and p-values, is provided in the Supplementary Material (Table S3). GO enrichment analysis of DAPs at 50 days indicated associations with defense and immune responses, intracellular signal transduction, and regulation of cell communication (Fig. 3A).

**Figure 3:**
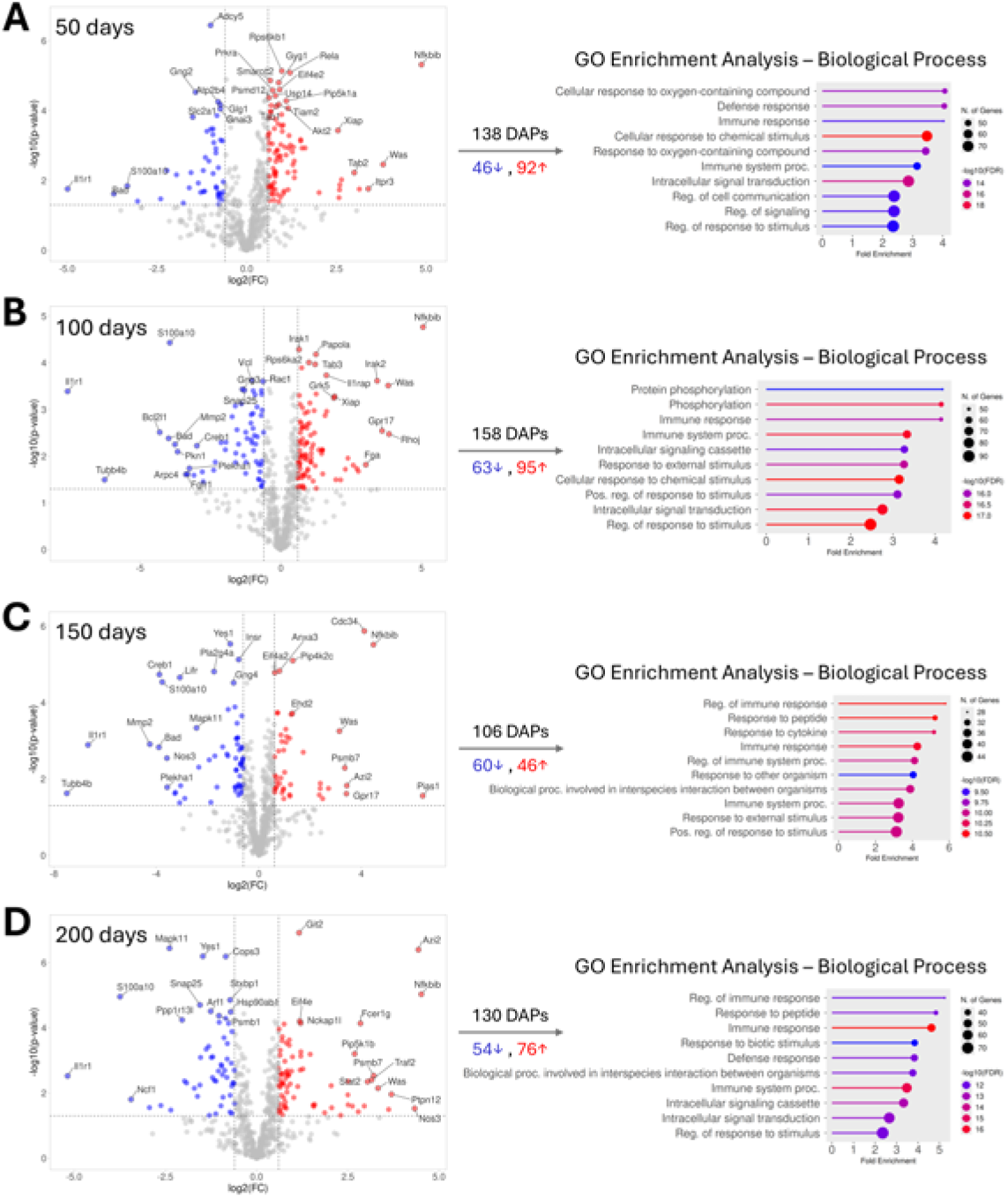
Differentially abundant proteins between B6 and hA7ko mice across ages. Volcano plots show DAPs at 50 (A), 100 (B), 150 (C), and 200 days (D), with proteins significantly increased (red dots) or decreased (blue dots) in hA7ko brains relative to B6 (DAPs were defined as p-value *≤* 0.05 and |FC| *≥* 1.5, corresponding to -log10(pvalue) *≥* 1.30 and |log2(FC)| *≥* 0.585). Top 25 DAPs with the largest and most significant abundance changes are labeled, and the number of DAPs (total, increased, decreased) is shown as well. For each age, lollipop plots on the right summarize the top GO biological process terms (up to 10) enriched among all DAPs (upand downregulated together), suggesting predominant alterations in immune and intracellular signaling processes and cellular responses to peptide stimuli.

At 100 days, 158 DAPs were identified, of which 95 were increased and 63 decreased in hA7ko relative to B6 (Fig. 3B). The most prominently upregulated proteins included NFKBIB (NF-*κ*B signaling inhibitor), IRAK1 (an IL-1 receptor-associated serine/threonine kinase), PAPOLA (a 3’poly(A) polymerase involved in mRNA processing), RPS6KA2 (a serine/threonine kinase involved in MAPK signaling), and TAB3 (an adaptor protein in the JNK and NF-*κ*B pathways). Among the most strongly downregulated DAPs were S100A10 (an S100 family protein associated with membrane repair and plasminogen activation), VCL (an actin filament-binding protein), RAC1 (a Rho family small GTPase regulating actin dynamics and cell signaling), GNG3 (a gamma subunit of heterotrimeric G proteins), and SNAP25 (a SNARE protein involved in vesicle docking and membrane fusion). GO enrichment analysis at 100 days suggested alterations in immune response, intracellular signal transduction, and regulation of cellular responses by protein phosphorylation (Fig. 3B).

At 150 days, the number of DAPs decreased to 106, with fewer upregulated (46) than downregulated (60) proteins (Fig. 3C). The most strongly upregulated proteins included CDC34 (a ubiquitin-conjugating E2 enzyme), NFKBIB (NF-*κ*B signaling inhibitor), PIP4K2C (a phosphatidylinositol-5-phosphate 4-kinase that regulates phosphoinositide signaling at cellular membranes), ANXA3 (a calcium-dependent phospholipid-binding protein), and EIF4A2 (an RNA helicase involved in translation initiation). Among the most strongly downregulated DAPs were YES1 (a non-receptor tyrosine kinase regulating cell growth, survival, and differentiation), INSR (insulin receptor, a plasma membrane receptor tyrosine kinase), PLA2G4A (a cytosolic phospholipase, releasing arachidonic acid from membrane phospholipids), CREB1 (a cAMP-responsive transcription factor), and LIFR (leukemia inhibitory factor receptor). GO enrichment analysis of up- and downregulated DAPs at 150 days reinforced that *Abca7* deficiency-induced proteomic changes are associated with cellular responses to peptide and immune stimuli (Fig. 3C).

At 200 days, the number of DAPs increased again to 130, with 76 upregulated and 54 downregulated proteins (Fig. 3D). The most strongly upregulated DAPs included GIT2 (a GTPase-activating protein and scaffolding molecule), AZI2 (an adaptor protein that promotes NF-*κ*B and IRF signaling), NFKBIB (NF-*κ*B signaling inhibitor), EIF4E (a key regulator of translation initiation), and NCKAP1L (a regulatory protein controlling actin cytoskeleton dynamics in hematopoietic cells). Conversely, MAPK11 (a serine/threonine kinase in stress-activated MAPK signaling), YES1 (a non-receptor tyrosine kinase), COPS3 (a subunit of the COP9 signalosome complex that regulates protein ubiquitination), S100A10 (S100 family protein), and STXBP1 (a syntaxin-binding protein regulating synaptic vesicle exocytosis) were among the most strongly downregulated proteins. GO enrichment analysis of DAPs at 200 days indicated that *Abca7* deficiency is associated with altered intracellular signaling processes induced by immune and peptide stimuli (Fig. 3D).

Across ages, many of the most strongly altered proteins are receptors, ion transporters, Gprotein subunits, SNARE components, or adaptor and signaling molecules that reside at, or closely associate with the plasma membrane or membrane-related signaling complexes. This observation was reinforced by subcellular compartment enrichment analysis of the detected DAPs, which showed that the plasma membrane was the most significantly enriched subcellular compartment at each age. In addition, endosome, intracellular vesicle, nucleus, and cytoskeleton were enriched at 50 days; nucleus, cytoskeleton, and extracellular region at 100 days; nucleus at 150 days; and cytoskeleton and nucleus at 200 days (Supplementary Material, Fig. S8, first column). This pattern is consistent with the known role of ABCA7 in plasma membrane-associated lipid metabolism and transport, suggesting that *Abca7* deficiency may preferentially impact membrane-associated signaling pathways that ultimately lead to gene expression changes in the nucleus.

To identify potential interactions among DAPs detected in hA7ko animals, we mapped functional PPIs using STRING and visualized the networks in Cytoscape. At 50 days, 113 of 138 DAPs (82%) were incorporated into the PPI network, with 104 proteins forming a major connected component and four small subnetworks comprising 2, 2, 2, and 3 proteins, respectively (Fig. 4A). In the main component, the 104 proteins were connected by 427 STRING interactions, largely supported by coexpression, experimentally determined interactions, and annotated database information. Among the 113 network DAPs, 23 were specific to the B6 vs. hA7ko comparison. These comparison-specific DAPs were mainly associated with biological processes related to immune responses and general cellular signaling, and several of them (e.g., AKT1, MAPK14, PSMD12) acted as local hubs with high numbers of connections. Considering the entire network, AKT1, AKT2, AKT3, SRC, and RAC1 were identified as the top hub proteins. GLay clustering partitioned the network into 10 clusters, 6 of which were interconnected, whereas 4 formed independent peripheral clusters. GO enrichment analysis of these clusters suggested that they represent biologically distinct functional modules, including processes such as regulation of cytoplasmic translation, cellular responses to increased oxygen levels, and prostaglandin biosynthesis. The clustered network, together with the top biological process terms per cluster and the direction and magnitude of change (node size and color), is shown in the Supplementary Material (Fig. S9A).

**Figure 4:**
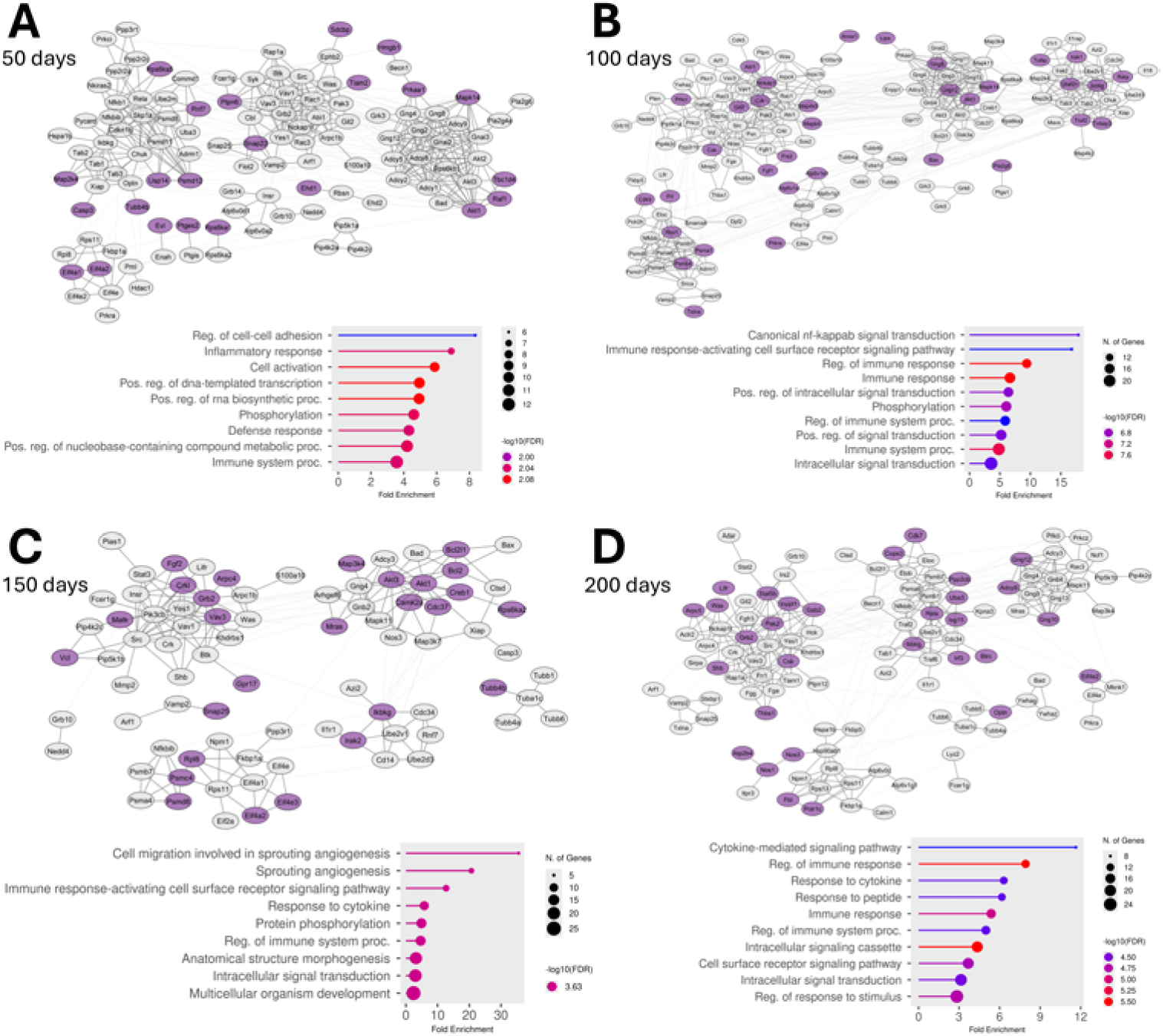
Protein-protein interaction networks of differentially abundant proteins between B6 and hA7ko mice across ages. STRING-based PPI networks are shown for DAPs at 50 (A), 100 (B), 150 (C), and 200 days (D). Nodes represent DAPs and edges represent known or predicted functional interactions. The layout of the networks reflects clustering performed in Cytoscape, with thicker edges indicating within-cluster PPIs and thinner edges indicating between-cluster PPIs. Comparison-specific DAPs (unique to the B6 vs. hA7ko comparison at a given age) are highlighted in the networks (purple nodes). For each age, lollipop plots under the network figures show the top GO biological process terms (up to 10) enriched among the comparison-specific DAPs, indicating that *Abca7* deficiency is associated with age-dependent changes in immune and cytokine responses, intracellular signaling pathways, and membrane-associated processes.

At 100 days, 139 of 158 DAPs (88%) were included in the PPI network, with 131 proteins forming a major connected component linked by 581 interactions, and two small subnetworks containing 6 and 2 proteins, respectively (Fig. 4B). Of the 139 network DAPs, 34 were specific for the B6 vs. hA7ko comparison. These comparison-specific DAPs were predominantly related to inflammatory and other immune system processes. As at 50 days, comparison-specific AKT1 and MAPK14 appeared as hub proteins with a high number of connections. In the whole network, SRC, AKT1, AKT2, AKT3, and NRAS were the top five hub proteins. Clustering of the network revealed 8 protein clusters, 6 of which were interconnected, while 2 formed independent peripheral clusters (Supplementary Material, Fig. S9B). GO enrichment analysis of these clusters indicated that they may reflect *Abca7* deficiency-related changes in biological processes such as SNARE complex assembly, Toll-like receptor signaling and K63-linked protein ubiquitination, negative regulation of fatty acid transport across the plasma membrane, and synaptic vesicle lumen acidification.

At 150 days, PPI analysis showed that 83 of 106 DAPs (78%) participated in functional interactions. The main network component comprised 78 proteins connected by 196 interactions, and a small independent subnetwork contained 5 proteins (Fig. 4C). Within the main network, 27 DAPs were specific for the B6 vs. hA7ko comparison. These comparison-specific DAPs appeared to be involved in angiogenesis, responses to cytokines, and other immune responses mediated by cell surface receptor signaling pathways. Among the comparison-specific DAPs, GRB2 and AKT1 had the highest numbers of connections, whereas in the whole network PIK3CB, SRC, GRB2, AKT1, and AKT3 were the top hub proteins. Clustering revealed 7 protein clusters, 6 of which were interconnected and 1 of which formed an independent peripheral cluster (Supplementary Material, Fig. S9C). GO enrichment analysis indicated that these clusters correspond to distinct biological processes, including regulation of immune cell diapedesis and protein polyubiquitination, SNARE complex assembly, translational regulation, and microtubule cytoskeleton organization.

At 200 days, 112 of 130 DAPs (86%) were incorporated into the PPI network, with 110 proteins and 339 interactions forming a major component and 2 proteins forming a small independent subnetwork (Fig. 4D). In total, 30 DAPs in the network were specific for the B6 vs. hA7ko comparison. These comparison-specific DAPs were involved in processes such as cellular responses to cytokines and peptides and intracellular signaling triggered by cell surface receptor activation. Among the comparison-specific DAPs, GRB2 and GNG10 were local hubs with high connectivity, while in the whole network GRB2, SRC, RAC3, NCKAP1L, and GNG12 were the top hub proteins. The network was partitioned into 8 clusters, 7 of which were interconnected, with the small 2-protein subnetwork forming the remaining independent cluster (Supplementary Material, Fig. S9D). GO enrichment analysis of these clusters showed that each represents a functionally distinct group of interconnected proteins involved in processes such as negative regulation of fatty acid transport across the plasma membrane, positive regulation of vesicle docking, and CD40 signaling.

### 3.4. APP expression in Abca7-deficient brain is associated with altered immune, hormonal, and receptor-mediated signaling across ages

At 50 days, cerebral APP expression in the *Abca7* -deficient background (hA7ko vs. APP-hA7ko) was associated with 156 DAPs, of which 63 were increased and 93 decreased in APP-hA7ko relative to hA7ko (Fig. 5A). The most significantly upregulated proteins included ATP6V0C (a V-type proton ATPase subunit involved in vesicular acidification), ADCY5 (a membrane-bound adenylate cyclase), SIRPA (an immunoglobulin-like cell surface receptor for CD47), ADCY9 (adenylate cyclase type 9), and EPHB2 (a receptor tyrosine kinase involved in axon guidance and synaptic plasticity). Among the most significantly downregulated proteins were TAB1 (a key adaptor protein in JNK and NF-*κ*B signaling), NFKBIB (NF-*κ*B signaling inhibitor), RELA (the p65 subunit of the NF-*κ*B transcription factor complex), RBSN (a Rab4/Rab5 effector protein involved in endocytic membrane fusion and trafficking), and HDAC5 (a histone deacetylase regulating transcription). GO enrichment analysis of DAPs at 50 days indicated changes in cellular responses to chemical, immunological, and oxygen-containing stimuli, cell surface receptor signaling, and intracellular signal transduction (Fig. 5A).

**Figure 5:**
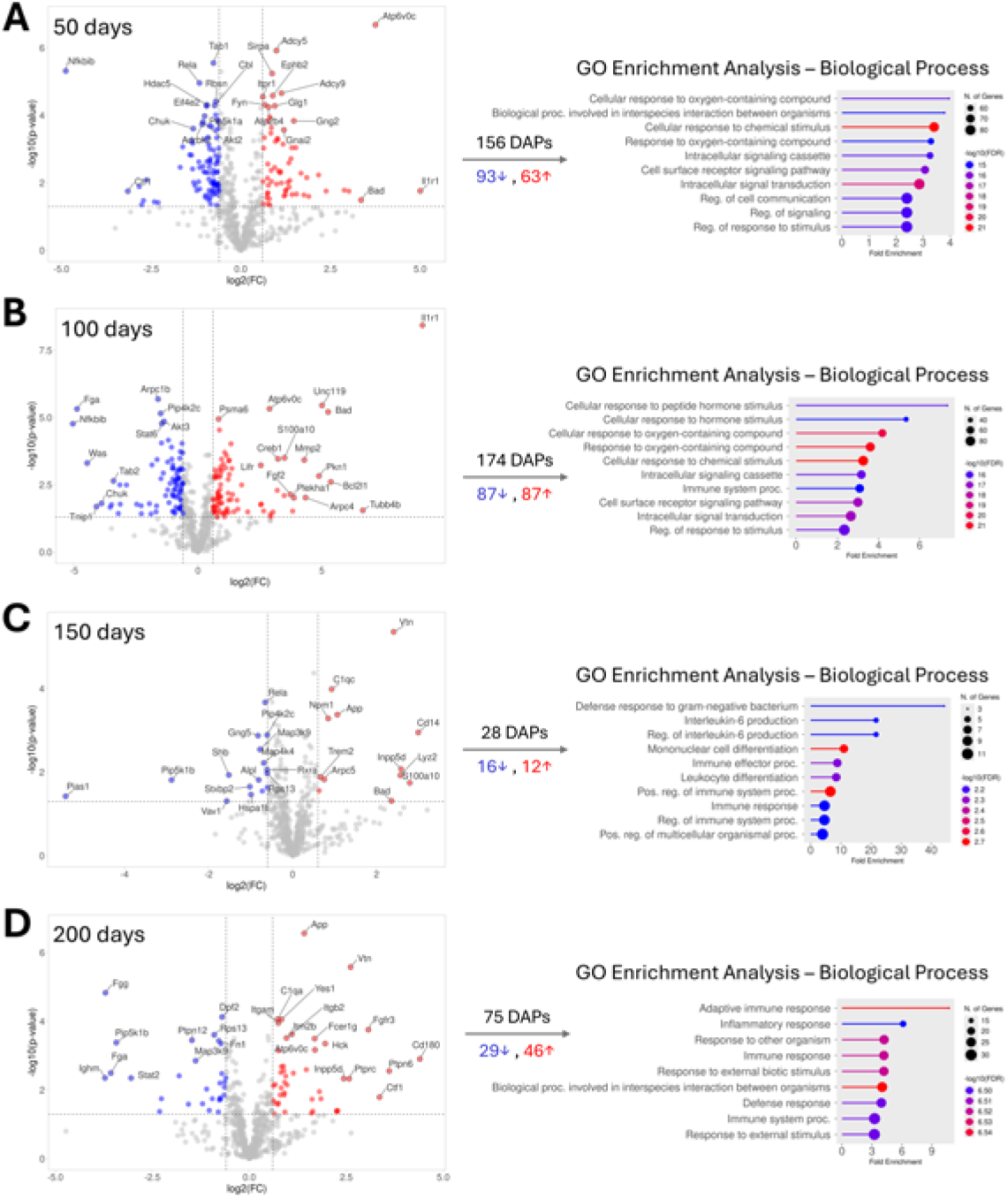
Differentially abundant proteins between hA7ko and APP-hA7ko mice across ages. Volcano plots show DAPs at 50 (A), 100 (B), 150 (C), and 200 days (D), with proteins significantly increased (red dots) or decreased (blue dots) in APP-hA7ko brains relative to hA7ko (DAPs were defined as p-value *≤* 0.05 and |FC| *≥* 1.5, corresponding to -log10(p-value) *≥* 1.30 and |log2(FC)| *≥* 0.585). Top 25 DAPs with the largest and most significant abundance changes are labeled, and the number of DAPs (total, increased, decreased) is shown as well. For each age, lollipop plots on the right summarize the top GO biological process terms (up to 10) enriched among all DAPs (upand downregulated together), highlighting predominant alterations in immune and inflammatory responses, hormone and receptor-mediated signaling, and cellular responses to external and endogenous stimuli.

At 100 days, 174 DAPs were identified, with equal numbers of proteins increased and decreased (87 upregulated and 87 downregulated in APP-hA7ko compared with hA7ko; Fig. 5B). The most significantly upregulated proteins included IL1R1 (IL-1 receptor, a key mediator of pro-inflammatory signaling), UNC119 (a lipid-binding chaperone that regulates SRC-family kinases), ATP6V0C (a V-type proton ATPase subunit), BAD (a pro-apoptotic Bcl-2 family protein), and PSMA6 (a 20S proteasome subunit). The most significantly downregulated DAPs included ARPC1B (a component of the Arp2/3 complex mediating actin polymerization), FGA (fibrinogen alpha chain), PIP4K2C (a phosphatidylinositol-5-phosphate 4-kinase modulating phosphoinositide signaling), AKT3 (a serine/threonine kinase in PI3K/AKT signaling), and STAT6 (a transcription factor involved in cytokine signaling pathways). GO enrichment analysis of DAPs at 100 days pointed to altered cellular responses to hormones and peptide hormones, immune system processes, and cell surface receptormediated intracellular signaling (Fig. 5B).

At 150 days, the number of DAPs dropped markedly to 28, with 12 proteins increased and 16 decreased in APP-hA7ko relative to hA7ko (Fig. 5C). The most significantly upregulated proteins included VTN (vitronectin, an extracellular matrix protein involved in cell adhesion and inflammation), C1QC (core component of the complement C1 complex), APP (amyloid-beta precursor protein), NPM1 (a multifunctional nucleolar protein involved in ribosome biogenesis, protein chaperoning, and cell ploriferation), and CD14 (a coreceptor for bacterial lipopolysaccharide). The most significantly downregulated DAPs included RELA (NF-*κ*B p65 subunit), PIP4K2C (a phosphatidylinositol-5-phosphate 4-kinase), GNG5 (a gamma subunit of heterotrimeric G proteins), MAP3K9 (a serine/threonine kinase involved in the MAP kinase signaling pathway), and MAP4K4 (a serine/threonine kinase implicated in stress response and inflammatory signaling). GO enrichment analysis at 150 days indicated a focused modulation of immune effector processes, including cytokine (IL-6) production and immune cell differentiation (Fig. 5C).

At 200 days, 75 DAPs were detected, with 46 upregulated and 29 downregulated in APP-hA7ko vs. hA7ko (Fig. 5D). The most significantly upregulated proteins included APP (amyloid-beta precursor protein), VTN (vitronectin), C1QA (a complement C1q subunit), YES1 (a non-receptor tyrosine kinase), and ITGAM (an integrin *α*-chain highly expressed in myeloid cells and microglia). Among the most significantly downregulated DAPs were FGG (fibrinogen gamma chain), DPF2 (a histone binding protein associated with transcriptional regulation), RPS13 (a component of the small ribosomal subunit), PTPN12 (a non-receptor tyrosine phosphatase regulating focal adhesion signaling), and FN1 (fibronectin 1, an extracellular matrix glycoprotein). GO enrichment analysis at 200 days mostly highlighted processes related to immunobiological processes, including inflammatory and adaptive immune responses (Fig. 5D).

Subcellular compartment enrichment analysis supported these observations. At 50 days, DAPs were significantly enriched at the plasma membrane, as well as in endosomes, intracellular vesicles, the cytoskeleton, nucleus, and vacuole. At 100 days, the plasma membrane and nucleus were the main enriched compartments, whereas at 150 days enrichment was restricted to the extracellular region and vacuole, and at 200 days to the extracellular region and plasma membrane (Supplementary Material, Fig. S8, second column). Together, these data suggest that APP expression on an *Abca7* -deficient background primarily affects plasma membrane-associated and extracellular proteins involved in immune and receptor-mediated signaling, with age-dependent shifts in nuclear and vesicular components

To explore how these proteins interact, we visualized the STRING-based PPI networks of DAPs across ages using Cytoscape (Fig. 6). At 50 days, 130 of 156 DAPs (83%) were incorporated into the PPI network, with 126 proteins forming a major connected component linked by 539 interactions and two small subnetworks each composed of 2 proteins (Fig. 6A). Thirty-six network DAPs were specific to the hA7ko vs. APP-hA7ko comparison at this age. These comparison-specific proteins were associated with processes such as regulation of immune cell activation, regulation of protein modification, and pathways linked to cell cycle and cell migration. Among comparison-specific DAPs, GNG11, GNG3, and PPP2R1B acted as local hubs, while across the entire network SRC, AKT2, AKT3, GRB2, and FYN were the most highly connected proteins. GLay clustering divided the network into 9 clusters, 7 of which were interconnected and 2 of which formed independent peripheral clusters (Supplementary Material, Fig. S10A). GO enrichment analysis of these clusters suggested functionally distinct groups of proteins related to processes like Toll-like receptor signaling, vesicular and vacuolar transport, cAMP and phosphoinositide metabolism, regulation of calcium-dependent neurotransmitter release, synaptic vesicle lumen acidification, and Golgi-lysosome/endosome trafficking.

**Figure 6:**
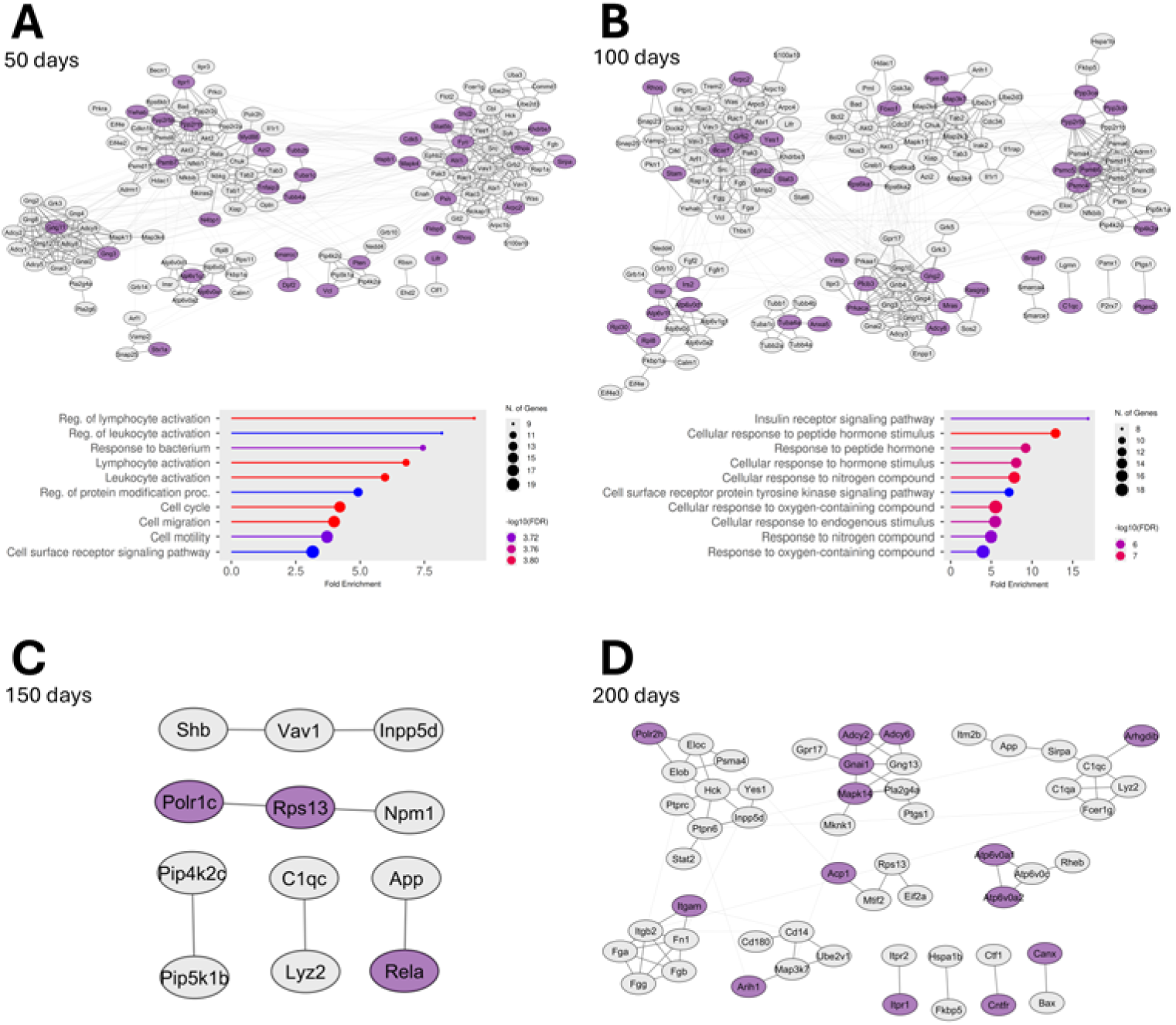
Protein-protein interaction networks of differentially abundant proteins between hA7ko and APP-hA7ko mice across ages. STRING-based PPI networks are shown for DAPs at 50 (A), 100 (B), 150 (C), and 200 days (D). Nodes represent DAPs and edges represent known or predicted functional interactions. The layout of the networks reflects clustering performed in Cytoscape, with thicker edges indicating within-cluster PPIs and thinner edges indicating between-cluster PPIs. Comparison-specific DAPs (unique to the hA7ko vs. APP-hA7ko comparison at a given age) are highlighted in the networks (purple nodes). For 50and 100-days old animals, lollipop plots under the network figures show the top 10 GO biological process terms enriched among the comparison-specific DAPs, suggesting that APP expression on an *Abca7* -deficient background is associated with age-dependent alterations in immune and cytokine signaling, hormone and receptor tyrosine kinase pathways, and vesicleand cytoskeleton-related processes. At 150 and 200 days, the small subsets of comparison-specific DAPs did not result in significant GO term enrichment.

At 100 days, 152 of 174 DAPs (87%) participated in the PPI network, with 146 proteins forming a major component connected by 595 interactions and three small subnetworks composed of 2 proteins each (Fig. 6B). Among these, 37 DAPs were specific to the hA7ko vs. APP-hA7ko comparison at this age. These comparison-specific proteins were enriched for processes related to insulin receptor signaling, cellular responses to peptide hormones and endogenous stimuli, and receptor tyrosine kinase-mediated signaling. GRB2, GNG2, and PPP2R5B could be identified as local hubs among the comparison-specific DAPs, whereas SRC, GRB2, AKT2, AKT3, and RAC1 were the top hubs in the entire network. Clustering identified 10 protein clusters, 7 interconnected and 3 small peripheral clusters (Supplementary Material, Fig. S10B). GO enrichment analysis of the clusters indicated that these subnetworks might be associated with biologically distinct processes, e.g., microtubule cytoskeleton organization, regulation of Toll-like receptor 6 signaling, protein-DNA complex remodeling, prostaglandin biosynthesis, protein localization to lipid droplets, phosphoinositide metabolism, and synaptic vesicle lumen acidification.

At 150 days, only 12 of 28 DAPs (43%) were connected in the STRING network, forming five independent small subnetworks: two consisting of 3-3 proteins and 2-2 interactions, and three consisting of 2 proteins linked by a single interaction each (Fig. 6C). Three of these 12 network DAPs were specific for the hA7ko vs. APP-hA7ko comparison, but this small set did not show significant GO enrichment, and no hub proteins were defined. Clustering correspondingly defined 5 independent clusters (Supplementary Material, Fig. S10C). Only two of them could be associated with a few GO terms, including immune cell processes, and processes related to APP catabolism and regulation of miRNA transcription.

At 200 days, 54 of 75 DAPs (72%) were incorporated into the PPI network, with 42 proteins forming a main component connected by 69 interactions and 5 additional small subnetworks (four formed by 2 proteins each, and one by 4 proteins with 4 interactions; Fig. 6D). Fourteen of the network DAPs were specific to the hA7ko vs. APP-hA7ko comparison at this age, but this small subset of proteins did not show significant GO enrichment. Among comparison-specific DAPs, GNAI1 and MAPK14 could be considered local hubs, whereas across the entire network ITGB2, GNAI1, HCK, PTPN6, and FN1 had the highest number of connections. GLay clustering identified 11 clusters, 6 of which were interconnected and 5 were independent (Supplementary Material, Fig. S10D). GO enrichment analysis of the clusters associated them with processes like target-directed miRNA degradation, cellular responses to cAMP, translation, synapse pruning and cell junction disassembly, and vacuolar and synaptic vesicle acidification.

Overall, APP expression on an *Abca7* -deficient background is associated with age-dependent remodeling of plasma membrane-related, vesicular, and extracellular protein networks in the murine brain, with prominent effects on immune and cytokine signaling, receptorand hormone-driven pathways (e.g., insulin receptor signaling), and vesicle-related processes.

### 3.5. Abca7 deficiency is associated with age-dependent immune and stress pathway changes in APP-expressing mouse brain

At 50 days, *Abca7* deletion in APP-expressing mice (APP vs. APP-hA7ko) was associated with a small set of 14 DAPs, of which 4 were increased and 10 decreased in APP-hA7ko relative to APP (Fig. 7A). The upregulated proteins were PRKCZ (an atypical protein kinase C involved in NF-*κ*B signaling and cell polarity), S100A1 (a calcium-binding protein involved in calcium-dependent signaling processes), MAPK11 (a stress-activated serine/threonine kinase), and HCK (a non-receptor tyrosine kinase expressed in myeloid cells). The most significantly downregulated proteins included FGG and FGB (fibrinogen gamma and beta chains), PFDN2 (a prefoldin subunit involved in protein folding), LGMN (a lysosomal/endosomal protease involved in antigen processing), and HDAC1 (a histone deacetylase regulating chromatin structure and transcription). GO enrichment analysis of DAPs at 50 days indicated altered cellular responses to inflammatory and calcium signals, including IL-1-mediated responses, positive regulation of ERK1/2 signaling, and modulation of apoptotic processes (Fig. 7A).

**Figure 7:**
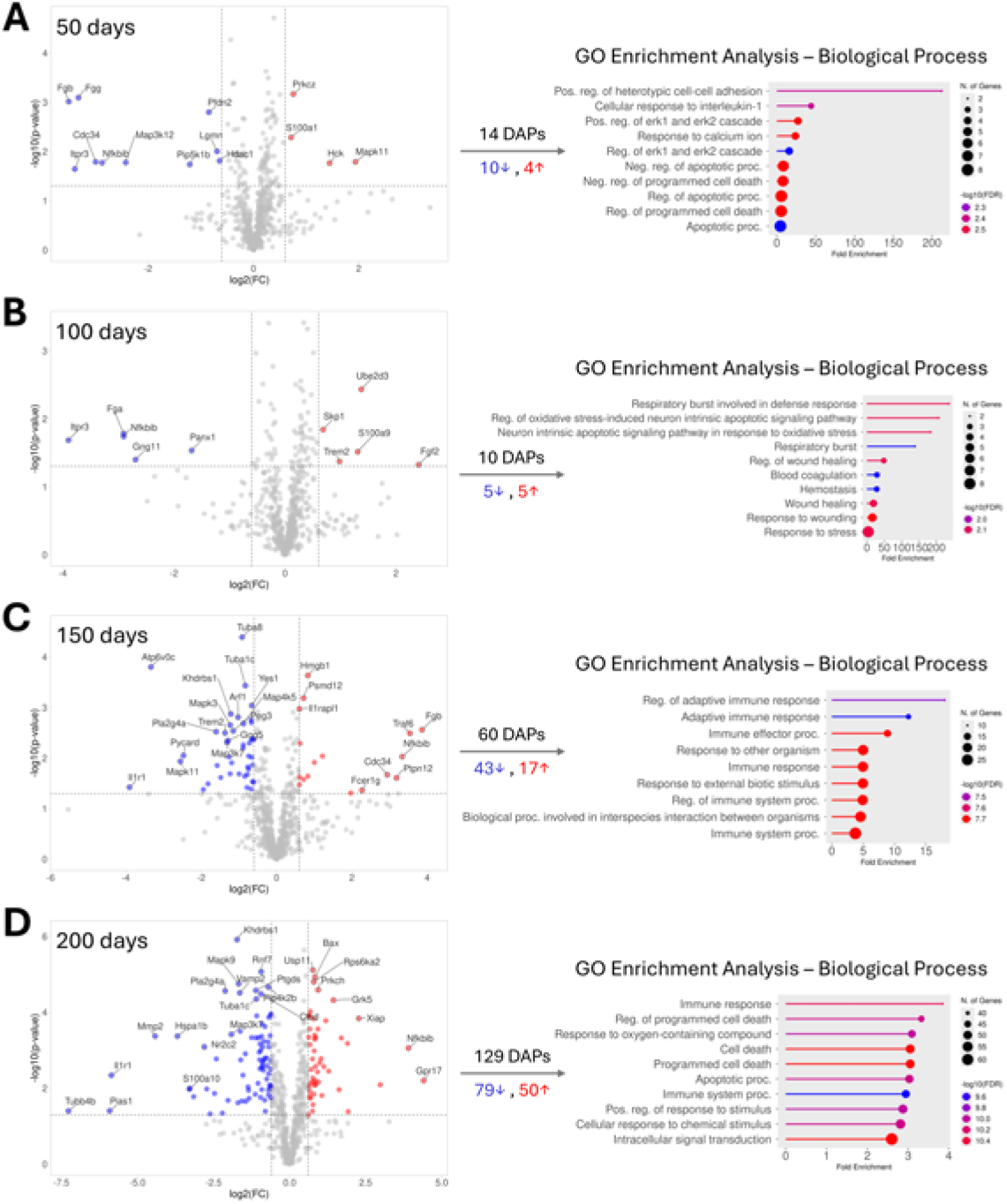
Differentially abundant proteins between APP and APP-hA7ko mice across ages. Volcano plots show DAPs at 50 (A), 100 (B), 150 (C), and 200 days (D), with proteins significantly increased (red dots) or decreased (blue dots) in APP-hA7ko brains compared with APP (DAPs were defined as p-value *≤* 0.05 and |FC| *≥* 1.5, corresponding to -log10(p-value) *≥* 1.30 and |log2(FC)| *≥* 0.585). Top DAPs with the largest and most significant abundance changes are labeled (up to 25 proteins), and the number of DAPs (total, increased, decreased) is shown as well. For each age, lollipop plots on the right summarize the top GO biological process terms (up to 10) enriched among all DAPs (upand downregulated together), suggesting predominant alterations in immune and inflammatory responses, respiratory burst and oxidative stress, and regulation of apoptotic processes.

At 100 days, only 10 DAPs were identified, with 5 proteins showing increased and 5 showing decreased LFQ intensities in APP-hA7ko compared with APP animals (Fig. 7B). The upregulated proteins were UBE2D3 (a ubiquitin-conjugating E2 enzyme), SKP1 (a core component of the SCF ubiquitin ligase complex), S100A9 (a calcium- and zinc-binding protein associated with inflammation), TREM2 (a microglial A*β*42 receptor involved in phagocytosis and inflammation), and FGF2 (fibroblast growth factor 2). Downregulated proteins comprised FGA (fibrinogen alpha chain), NFKBIB (NF-*κ*B signaling inhibitor), ITPR3 (an inositol 1,4,5-trisphosphate receptor regulating intracellular calcium release), PANX1 (an ion channel involved in a variety of physiological functions), and GNG11 (a gamma subunit of heterotrimeric G proteins). GO enrichment analysis at 100 days suggested modulation of innate immune and stress-related processes, including respiratory burst during defense responses, oxidative stress-induced neuronal apoptotic signaling, and general responses to tissue damage and stress (Fig. 7B).

At 150 days, the number of DAPs increased to 60, with 17 proteins upregulated and 43 downregulated in APP-hA7ko relative to APP (Fig. 7C). The most strongly upregulated proteins included HMGB1 (a multifunctional protein that can modulate innate and adaptive immune responses), PSMD12 (a regulatory subunit of the 26S proteasome), IL1RAPL1 (an IL-1 receptor accessory-like protein involved in synapse development), FGB (fibrinogen beta chain), and TRAF6 (an E3 ubiquitin ligase central to inflammatory signaling pathways). Among the most prominently down-regulated DAPs were TUBA8 and TUBA1C (alpha-tubulin isoforms important for microtubule structure), ATP6V0C (a V-type proton ATPase subunit), YES1 (a non-receptor tyrosine kinase), and KHDRBS1 (an RNA-binding protein involved both in immune and insulin signaling processes). GO enrichment analysis at 150 days indicated broad modulation of adaptive and innate immune pathways, including immune effector functions and regulation of immune system processes (Fig. 7C). At 200 days, the number of DAPs further increased to 129, with 50 proteins upregulated and 79 downregulated in APP-hA7ko compared with APP animals (Fig. 7D). The most significantly upregulated proteins included USP11 (a deubiquitinating enzyme that regulates the stability of ubiquitinated proteins), BAX (a pro-apoptotic Bcl-2 family protein), RPS6KA2 (a serine/threonine kinase involved in MAPK signaling), PRKCH (a phospholipid- and diacylglycerol-dependent serine/threonine kinase), and GRK5 (a G protein-coupled receptor kinase). The most significantly downregulated DAPs included KHDRBS1 (an RNA-binding protein), RNF7 (a catalytic component of various ubiquitin ligase complexes), MAPK9 (a stress-activated MAP kinase), PIP4K2B (a phosphatidylinositol-5-phosphate 4-kinase isoform), and PTGDS (prostaglandin D synthase, a lipid metabolism-related protein). GO enrichment analysis of DAPs at 200 days highlighted processes like immune responses, regulation of programmed cell death, and intracellular signal transduction (Fig. 7D).

Subcellular compartment enrichment analysis indicated that the brain proteomic effects of *Abca7* deletion in APP-expressing mice are not restricted to a single compartment. At 50 days, DAPs were enriched mostly in the cytoplasm. At 100 days, enrichment was observed in the extracellular region and plasma membrane, whereas at 150 days the plasma membrane was the predominant enriched compartment. At 200 days, DAPs were enriched in the plasma membrane, nucleus, and cytoplasm (Supplementary Material, Fig. S8, third column). These patterns suggest that *Abca7* deletion in APP-expressing mouse brain affects cytoplasmic and membrane-associated components early on, with increasing involvement of extracellular and nuclear compartments at later ages.

The in silico analysis of PPIs revealed that the extent of network organization also changed with age. At 50 and 100 days, no STRING-based PPIs were detected among the relatively few DAPs, suggesting that early proteomic changes in APP-hA7ko compared with APP animals are dispersed across proteins that do not form an interaction network. At 150 days, however, 45 of the 60 DAPs (75%) could be incorporated into a PPI network (Fig. 8A). The main network component comprised 33 proteins connected by 68 interactions, while 5 additional small subnetworks were identified (three pairs of proteins connected by single interactions and two subnetworks of three proteins and two interactions each). Fifteen of the network DAPs were specific to the APP vs. APP-hA7ko comparison at 150 days. These comparison-specific proteins were associated with processes such as positive regulation of cell-cell adhesion, regulation of immune system processes, and cell surface receptor signaling. MAPK1 and TRAF6 could be identified as local hubs among the comparison-specific DAPs, whereas in the whole network MAPK1, MAPK3, GNB2, GNG5, and GNG3 showed the highest connectivity. GLay clustering identified 9 clusters in the network, 4 of which were interconnected, while 5 formed independent peripheral clusters (Supplementary Material, Fig. S11A). GO enrichment analysis of the clusters revealed that they represent biologically distinct functional modules related to vesicle docking and exocytosis, IL-34-mediated signaling, cellular defense responses, and synaptic vesicle maturation and lumen acidification.

**Figure 8:**
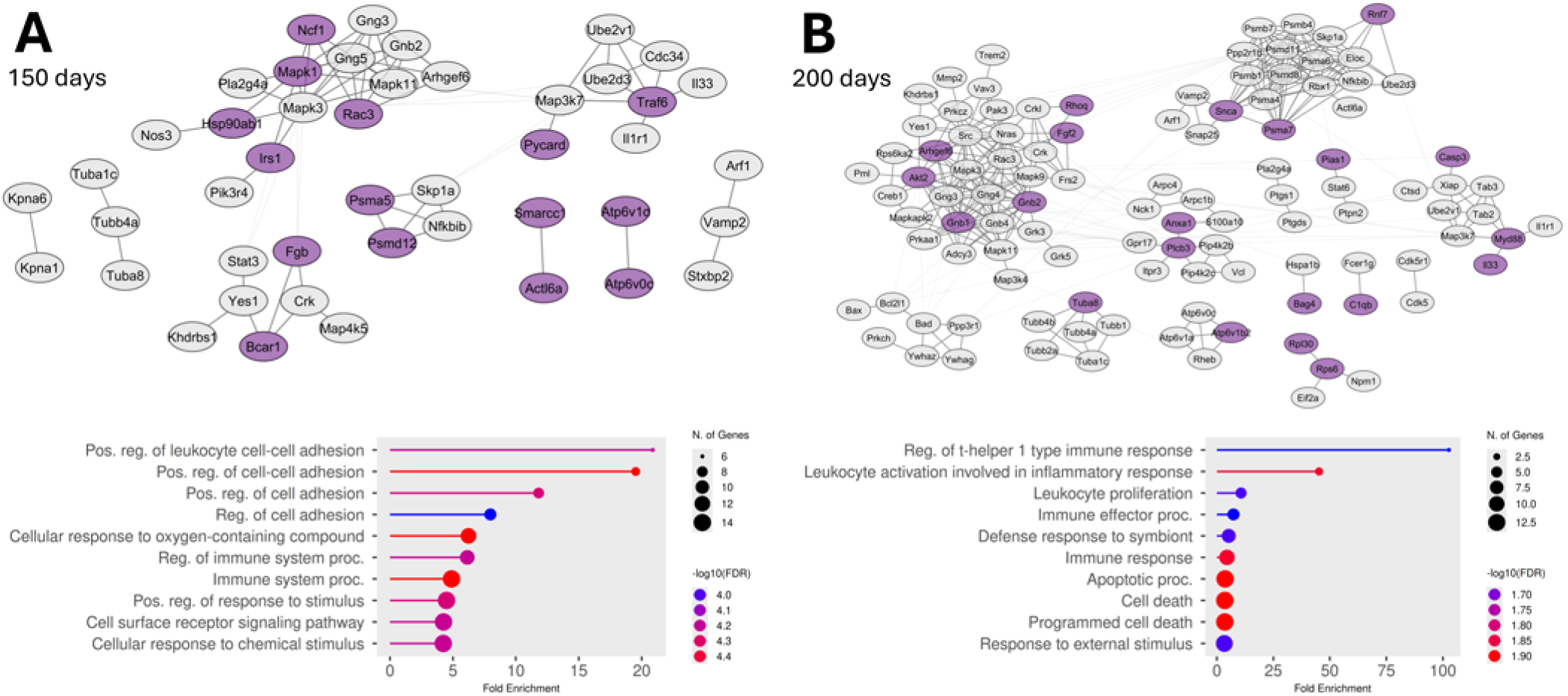
Protein-protein interaction networks of differentially abundant proteins between APP and APP-hA7ko mice across ages. STRING-based PPI networks are shown for DAPs at 150 (A) and 200 days (B). No PPIs were identified among the few DAPs detected at 50 and 100 days. Nodes represent DAPs and edges represent known or predicted functional interactions. The layout of the networks reflects clustering performed in Cytoscape, with thicker edges indicating within-cluster PPIs and thinner edges indicating between-cluster PPIs. Comparison-specific DAPs (unique to the APP vs. APP-hA7ko comparison at a given age) are highlighted in the networks (purple nodes). Lollipop plots under the network figures show the top 10 GO biological process terms enriched among the comparison-specific DAPs, suggesting that *Abca7* deletion in APP-expressing mice is associated with age-dependent changes in immune cell adhesion and activation, regulation of immune system and apoptotic processes, and cellular responses to external stimuli.

At 200 days, 108 of 129 DAPs (84%) were incorporated into the PPI network, with 94 proteins and 310 interactions forming a major network component and 4 additional independent subnetworks (one with 6 proteins and 8 interactions, one with 4 proteins and 3 interactions, and two with 2 proteins and single interactions each; Fig. 8B). Twenty-one DAPs in the network were specific to the APP vs. APP-hA7ko comparison at this age. These comparison-specific proteins were enriched for immune and cell death-related processes, including regulation of inflammatory immune cell activation and proliferation, immune effector functions, and apoptotic and programmed cell death pathways, as well as responses to external stimuli. Among the comparison-specific DAPs, GNB1, GNB2, and PSMA7 appeared as local hubs, while in the entire network MAPK3, SRC, NRAS, GNB1, and GNB4 were the most highly connected proteins. Clustering identified 13 protein clusters, 9 of which were interconnected and 4 formed small independent clusters (Supplementary Material, Fig. S11B). GO enrichment analysis of these clusters suggested that they may be associated with processes such as microtubule cytoskeleton organization, lymphocyte diapedesis and extravasation, SNARE complex assembly and long-term synaptic potentiation, cyclooxygenase and prostaglandin biosynthetic pathways, apoptotic signaling, phosphoinositide metabolism, IL-33 and p38 MAPK signaling, and synaptic vesicle maturation and lumen acidification.

Taken together, these data imply that *Abca7* deletion in APP-expressing mice might lead to age-dependent changes of cytoplasmic, plasma membrane, and nuclear protein networks in the brain, with increasingly pronounced effects on immune and inflammatory signaling, stress and apoptotic pathways, and vesicle- and receptor-related processes at later stages.

### 3.6. Combined Abca7 knockout and APP expression drive age-dependent brain proteomic changes associated with immune and stress pathways

At 50 days, comparison of APP-hA7ko mice with wild-type controls (B6 vs. APP-hA7ko) revealed 18 DAPs, of which 13 were increased and 5 decreased in APP-hA7ko relative to B6 (Fig. 9A). The most significantly upregulated proteins included CALM1 (calmodulin-1, a key calcium sensor in many signaling pathways), APP (amyloid-beta precursor protein), PPP3R1 (the regulatory subunit of calcineurin, a calcium-dependent phosphatase), ATP6V0C (a V-type proton ATPase subunit), and PRKCB (a calcium-activated, phospholipid- and diacylglycerol-dependent serine/threonine kinase). The downregulated proteins were TUBB6 (a beta-tubulin isoform), HSPA1B (an Hsp70 family chaperone), BTK (a non-receptor tyrosine kinase involved in B cell and microglial signaling), FGG (fibrinogen gamma chain), and NOS3 (endothelial nitric oxide synthase). GO enrichment analysis at 50 days indicated that these early proteomic changes predominantly affect cellular calcium handling and signaling, with overrepresentation of terms such as regulation of calcium ion sequestration and transport, regulation of calcium-mediated signaling, negative regulation of transmembrane transport, and vesicle-mediated transport in synapses, together with broader regulation of intracellular signal transduction (Fig. 9A).

**Figure 9:**
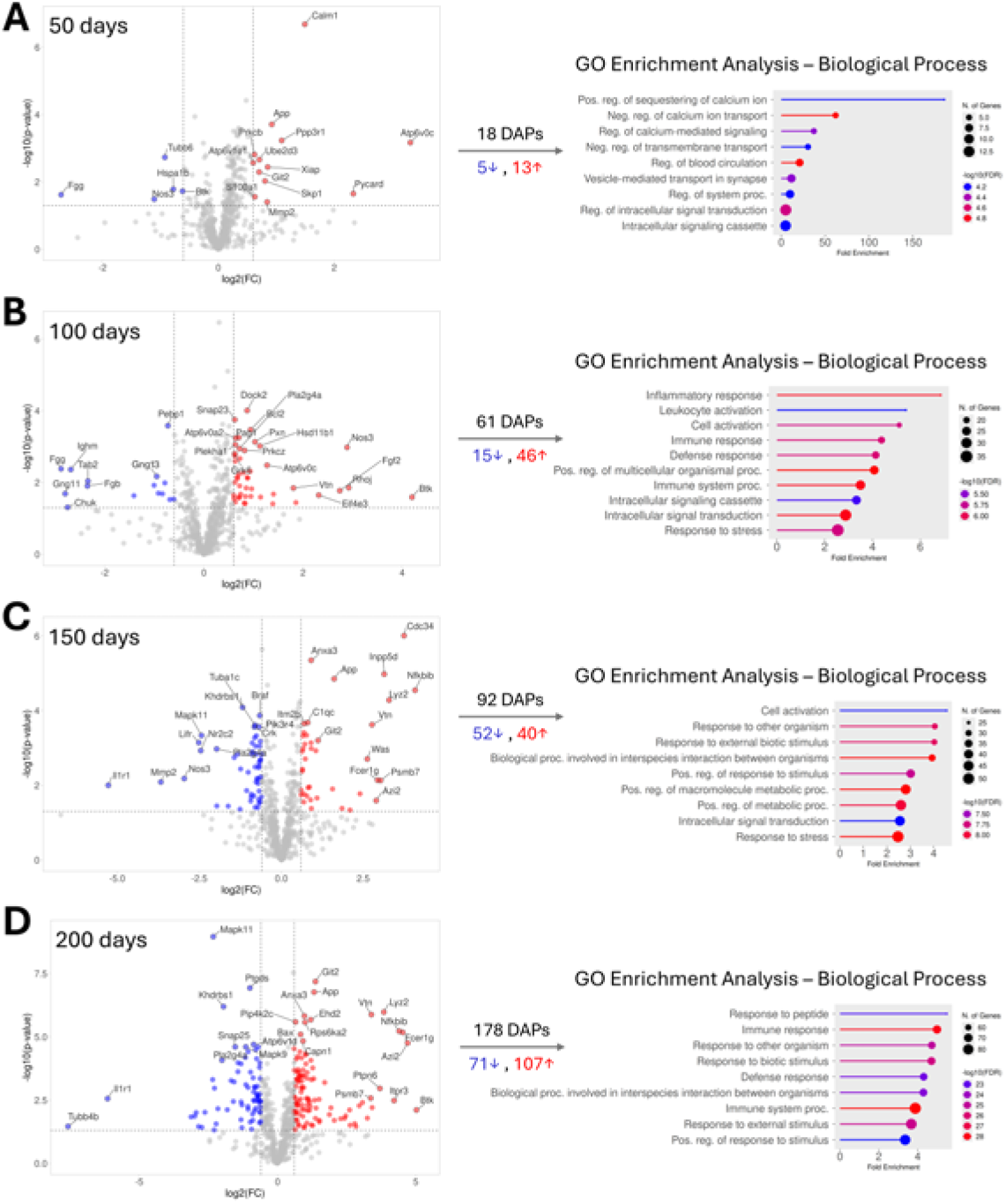
Differentially abundant proteins between B6 and APP-hA7ko mice across ages. Volcano plots show DAPs at 50 (A), 100 (B), 150 (C), and 200 days (D), with proteins significantly increased (red dots) or decreased (blue dots) in APP-hA7ko brains compared with B6 (DAPs were defined as p-value *≤* 0.05 and |FC| *≥* 1.5, corresponding to -log10(p-value) *≥* 1.30 and |log2(FC)| *≥* 0.585). Top DAPs with the largest and most significant abundance changes are labeled (up to 25 proteins), and the number of DAPs (total, increased, decreased) is shown as well. For each age, lollipop plots on the right summarize the top GO biological process terms (up to 10) enriched among all DAPs (upand downregulated together), highlighting predominant alterations in calcium-mediated signaling, inflammatory and defense responses, cell activation and intracellular signal transduction, and cellular responses to stress.

At 100 days, the number of DAPs increased to 61, with 46 proteins upregulated and 15 downregulated in APP-hA7ko compared with B6 (Fig. 9B). Among the most significantly upregulated proteins were DOCK2 (a guanine nucleotide exchange factor involved in lymphocyte migration and activation), SNAP23 (a SNARE protein involved in vesicle fusion and exocytosis), BCL2 (an anti-apoptotic Bcl-2 family protein), PAG1 (a transmembrane adaptor implicated in negative regulation of immune receptor signaling), and ATP6V0A2 (another V-type proton ATPase subunit). Prominent downregulated proteins included PEBP1 (a phosphatidylethanolamine-binding protein), FGG (fibrinogen gamma chain), IGHM (immunoglobulin mu chain), GNG13 (a gamma subunit of heterotrimeric G proteins), and TAB2 (an adaptor protein in JNK and NF-*κ*B signaling). GO enrichment analysis at 100 days indicated emerging immune activation and stress responses, with enrichment of inflammatory response, leukocyte and cell activation, immune and defense response, intracellular signal transduction, and response to stress (Fig. 9B).

At 150 days, 92 DAPs were identified, with 40 proteins showing increased and 52 showing decreased LFQ intensities in APP-hA7ko compared with B6 mice (Fig. 9C). The most strongly upregulated DAPs included CDC34 (a ubiquitin-conjugating E2 enzyme), ANXA3 (a calciumdependent phospholipid-binding protein), INPP5D (an inositol polyphosphate 5-phosphatase that negatively regulates PI3K/AKT signaling in immune cells), APP (amyloid-beta precursor protein), and NFKBIB (NF-*κ*B signaling inhibitor). Prominent downregulated proteins were KHDRBS1 (an RNA-binding protein), BRAF (a serine/threonine kinase in the MAPK pathway), TUBA1C (an alpha-tubulin isoform), PIK3R4 (a regulatory subunit of class III PI3K complexes), and CRK (an adaptor protein involved in cell branching and adhesion). GO enrichment analysis at 150 days suggested broader changes in immune and signaling processes, including cell activation, responses to external biotic stimuli, positive regulation of metabolic processes, intracellular signal transduction, and stress responses (Fig. 9C).

At 200 days, brain proteomic differences were most pronounced: 178 DAPs were detected, with 107 proteins upregulated and 71 downregulated in APP-hA7ko compared with B6 animals (Fig. 9D). The most significantly upregulated proteins were GIT2 (a GTPase-activating protein and scaffolding molecule), APP (amyloid-beta precursor protein), LYZ2 (an enzyme linked to innate immune and microglial activity), VTN (an extracellular matrix protein involved in cell adhesion and inflammation), and ANXA3 (a calcium-dependent phospholipid-binding protein). Among the most strongly downregulated DAPs were MAPK11 (a stress-activated serine/threonine kinase), PTGDS (prostaglandin D synthase), KHDRBS1 (an RNA-binding protein), ATP6V1D (V-type proton ATPase subunit), and SNAP25 (a SNARE protein involved in vesicle docking and membrane fusion). GO enrichment analysis at 200 days highlighted alterations in immune and stimulus-response pathways, including response to peptides, immune response, immune system processes, and responses to external stimuli (Fig. 9D).

Subcellular compartment enrichment analysis indicated that the brain proteomic impact of combined *Abca7* deletion and APP expression is associated with multiple subcellular compartments and becomes more widespread with age (Supplementary Material, Fig. S8, fourth column). At 50 days, DAPs were significantly enriched in the cytoskeleton, mitochondrion, plasma membrane, and nucleus, suggesting early alterations in structural, energetic, and signaling components. At 100 days, enrichment was restricted to the plasma membrane. At 150 days, no compartment showed significant enrichment, indicating a more diffuse effect at the level of the detected DAPs. At 200 days, however, DAPs were enriched in the plasma membrane, cytoplasm, extracellular region, nucleus, and cytoskeleton, consistent with extensive changes in both intra- and extracellular protein networks in APP-hA7ko brains at later stages.

The in silico PPI analysis further illustrated age-dependent changes in DAP network complexity and organization. At 50 days, only 6 of the 18 DAPs (33%) could be identified as PPI network components, forming three small subnetworks, each with 2 proteins and a single STRING interaction between them (Fig. 10A). Three of these network proteins were specific to the B6 vs. APP-hA7ko comparison at this age, but no meaningful GO enrichment or hub proteins were identified, and no proper clusters were detected (Supplementary Material, Fig. S12A). This suggests that early proteomic alterations in APP-hA7ko mice compared with B6 are rather modest and do not yet organize into a complex PPI network.

**Figure 10:**
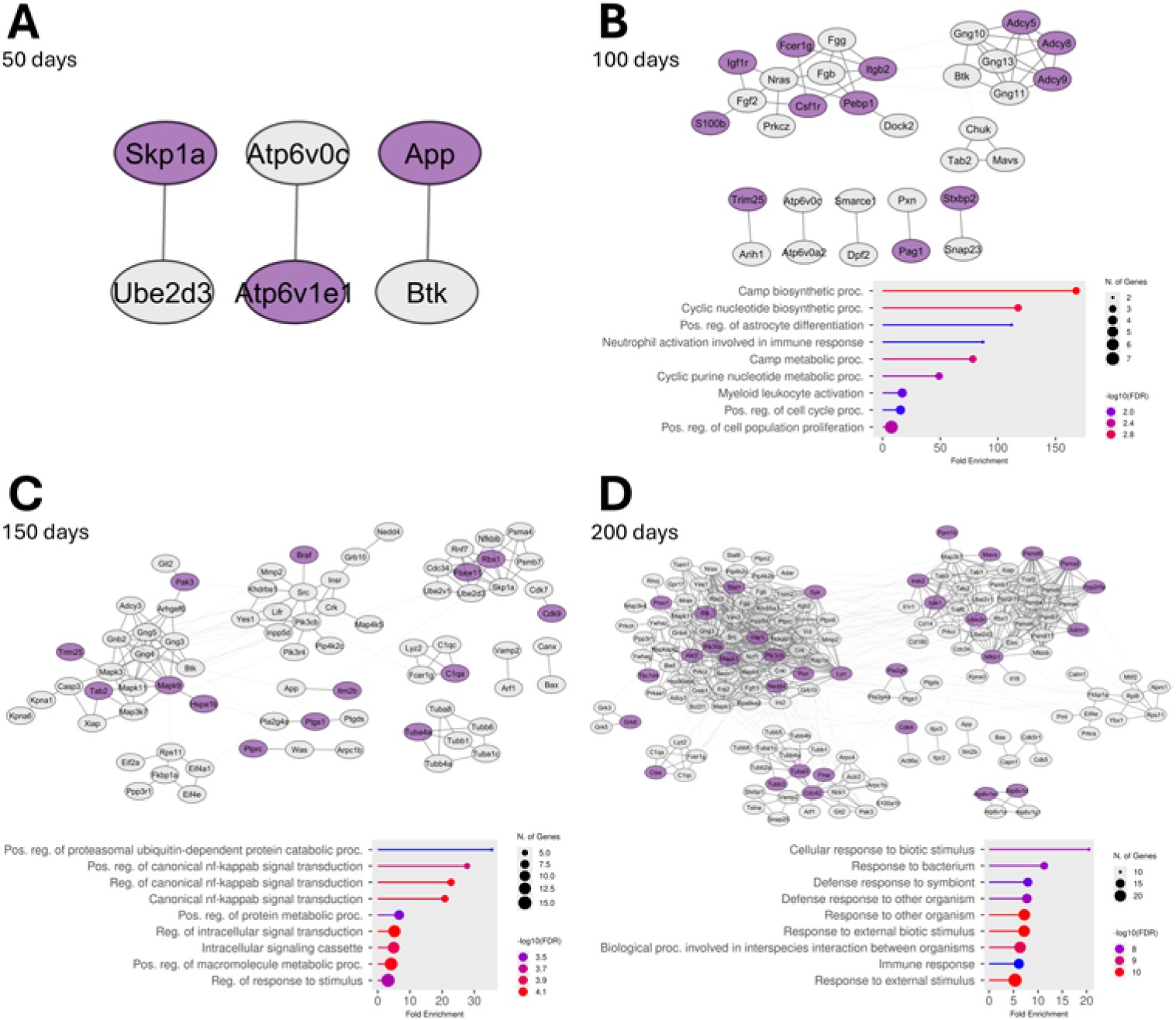
Protein-protein interaction networks of differentially abundant proteins between B6 and APP-hA7ko mice across ages. STRING-based PPI networks are shown for DAPs at 50 (A), 100 (B), 150 (C), and 200 days (D). Nodes represent DAPs and edges represent known or predicted functional interactions. The layout of the networks reflects clustering performed in Cytoscape, with thicker edges indicating within-cluster PPIs and thinner edges indicating between-cluster PPIs. Comparison-specific DAPs (unique to the B6 vs. APP-hA7ko comparison at a given age) are highlighted in the networks (purple nodes). Lollipop plots under the network figures show the top GO biological process terms (up to 10) enriched among the comparison-specific DAPs, suggesting that the combined effect of *Abca7* deletion and APP expression is associated with age-dependent changes in cAMP signaling, proteasomal ubiquitindependent protein degradation, inflammatory signal transduction, and cellular responses to immunological stimuli. At 50 days, the small subsets of comparison-specific DAPs did not result in significant GO term enrichment.

At 100 days, 32 of the 61 DAPs (52%) were incorporated into the PPI network (Fig. 10B). The major network component comprised 22 proteins connected by 43 interactions, while the remaining network DAPs formed 5 additional subnetworks of protein pairs. Twelve network DAPs were specific to the B6 vs. APP-hA7ko comparison. These comparison-specific DAPs were enriched for processes related to cyclic nucleotide signaling and immune activation, including cAMP biosynthetic and metabolic processes, positive regulation of astrocyte differentiation, neutrophil and myeloid leukocyte activation, and positive regulation of cell cycle and cell population proliferation. ADCY5 and ADCY9 had the highest number of local PPIs among comparison-specific DAPs, whereas in the entire network NRAS, GNG11, GNG10, GNG13, and ADCY5 showed the highest degrees. GLay clustering identified 8 clusters, of which 3 larger ones were interconnected, while the 5 small subnetworks of protein pairs clustered independently (Supplementary Material, Fig. S12B). GO enrichment of these clusters pointed to functional modules related to leukocyte degranulation and synaptic vesicle exocytosis, positive regulation of heterotypic cell adhesion, positive regulation of interferon-alpha production, cAMP biosynthetic and metabolic processes, and vacuolar acidification and intracellular pH reduction.

At 150 days, PPI network size and connectivity increased substantially. Seventy-five of the 92 DAPs (82%) were included in the PPI network (Fig. 10C). The major network component included 65 proteins connected by 136 interactions, and three additional subnetworks were detected (one with 6 proteins and 9 interactions, and two with protein pairs connected by single interactions). Fourteen of the network DAPs were specific to the B6 vs. APP-hA7ko comparison at this age. These comparison-specific proteins were associated with regulation of protein degradation and NF*κ*B signaling, including positive regulation of proteasomal ubiquitin-dependent protein catabolism, regulation of canonical NF-*κ*B signal transduction, positive regulation of protein and macromolecule metabolic processes, regulation of intracellular signaling, and regulation of responses to stimuli. MAPK9 and RBX1 were identified as local hubs among the comparison-specific DAPs, whereas in the entire network SRC, PIK3CB, MAPK3, GNG4, and GNB2 were the most highly connected proteins. Clustering divided the network into 12 clusters, 9 of which were interconnected, while 3 formed small peripheral clusters (Supplementary Material, Fig. S12C). GO enrichment analysis of these clusters associated them with processes like positive regulation of cGAS/STING signaling, stress-activated MAPK cascades, progesterone receptor signaling, synapse pruning and cell junction disassembly, transcription pausing by RNA polymerase II, translation and translational initiation, cyclooxygenase and prostaglandin biosynthetic pathways, and mitotic cell cycle and microtubule cytoskeleton organization.

At 200 days, the PPI network of DAPs became highly connected reflecting the high number of altered proteins at this age. A total of 162 of 178 DAPs (91%) could be integrated into the PPI network, with 156 proteins and 757 PPIs forming a major component and two additional small subnetworks (one with 4 proteins and 6 interactions, and one with a protein pair; Fig. 10D). Thirtythree of the network proteins were specific to the B6 vs. APP-hA7ko comparison at 200 days. These comparison-specific DAPs were enriched for immune and host defense processes, including cellular responses to biotic stimuli, bacteria, and other external stimuli. Among the comparison-specific DAPs, PIK3CA, PIK3CB, MAPK1, and CDC42 appeared as hubs, while in the whole network PIK3CA, PIK3CB, SRC, MAPK1, and MAPK3 had the highest degrees. Clustering resulted in 12 clusters, of which 10 were interconnected and 2 formed small peripheral clusters (Supplementary Material, Fig. S12D). GO enrichment of these clusters linked them to biological processes such as positive regulation of JNK activity, IL-33-mediated signaling, regulation of glutamate secretion and neurotransmission, modulation of Toll-like receptor 6 signaling, synaptic vesicle lumen acidification, synapse pruning and cell junction disassembly, cellular responses to cAMP, and phosphatidylcholine catabolism.

Overall, these findings indicate that in APP-hA7ko mice, combined *Abca7* deletion and APP expression lead to increasingly extensive and interconnected alterations in brain protein networks with age. Early changes predominantly affect calcium handling, vesicle trafficking, and plasma membrane-associated signaling, while later stages are characterized by widespread reorganization of cytoplasmic, plasma membrane, nuclear, and extracellular proteins, with prominent involvement of immune and inflammatory signaling, stress and apoptotic pathways, and synapseand receptorrelated processes.

### 3.7. Recurrently altered proteins identify core molecular processes associated with Abca7 deficiency, Aβ pathology, and their interaction

Based on manual curation of reviewed UniProtKB (Swiss-Prot) entries for the top DAPs in each genotype-age comparison, together with GO and PPI-based pathway annotation (see Figs. 3–10), several major molecular systems emerged as recurrently affected by *Abca7* deficiency and its interactions with A*β* pathology: the PI3K/AKT pathway and phosphoinositide signaling, the ubiquitin-proteasome protein degradation system, vesicle trafficking and exocytosis, vesicle lumen acidification, and MAPK signaling. To visualize how age and genotype jointly influence these pathways, we examined the age-dependent LFQ intensity changes of a set of frequently altered key proteins, selected as proteins that were repeatedly identified as DAPs across multiple comparisons and assigned to these key pathways, or that are strongly implicated in insulin signaling, neuroinflammatory processes, and AD based on the literature (Figs. 11 and 12). For each protein, mean LFQ intensities at 50, 100, 150, and 200 days were plotted for all four genotypes, allowing direct comparison of patterns of age-dependent abundance changes.

**Figure 11:**
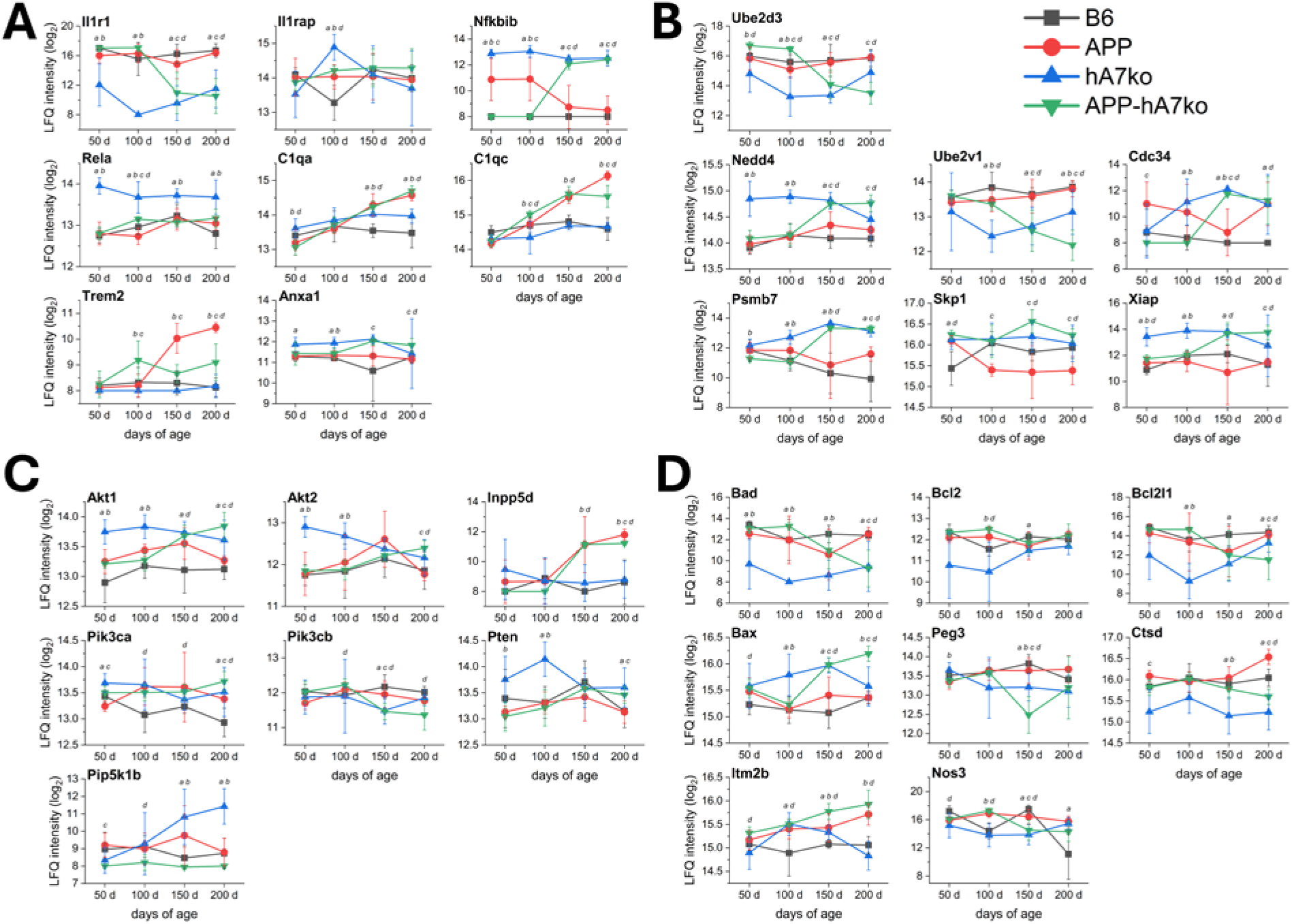
Age-dependent LFQ intensity profiles of immune, ubiquitin-proteasome, and signaling pathway proteins across genotypes. For each protein, mean LFQ intensities (*±* SD) are plotted at 50, 100, 150, and 200 days for B6 (grey), APP (red), hA7ko (blue), and APP-hA7ko (green) mice, with mean values connected by lines to illustrate age-dependent changes. The figure shows representative proteins that were recurrently identified as DAPs across multiple comparisons and are associated with key biological functions and processes identified by manual curation and GO enrichment analysis. (A) Inflammatory and immune-related proteins, including complement components and microglial/innate immune receptors. (B) Ubiquitin-proteasome system components involved in protein ubiquitination and proteasomal degradation. (C) PI3K/AKT pathway proteins and related phosphoinositide signaling regulators. (D) Additional AD-relevant proteins with diverse biological functions. Statistically significant differences (p *≤* 0.05; two-tailed Welch’s t-test) for the comparisons performed are indicated by letters at each age: a, B6 vs. hA7ko; b, hA7ko vs. APP-hA7ko; c, APP vs. APP-hA7ko; d, B6 vs. APP-hA7ko.

**Figure 12:**
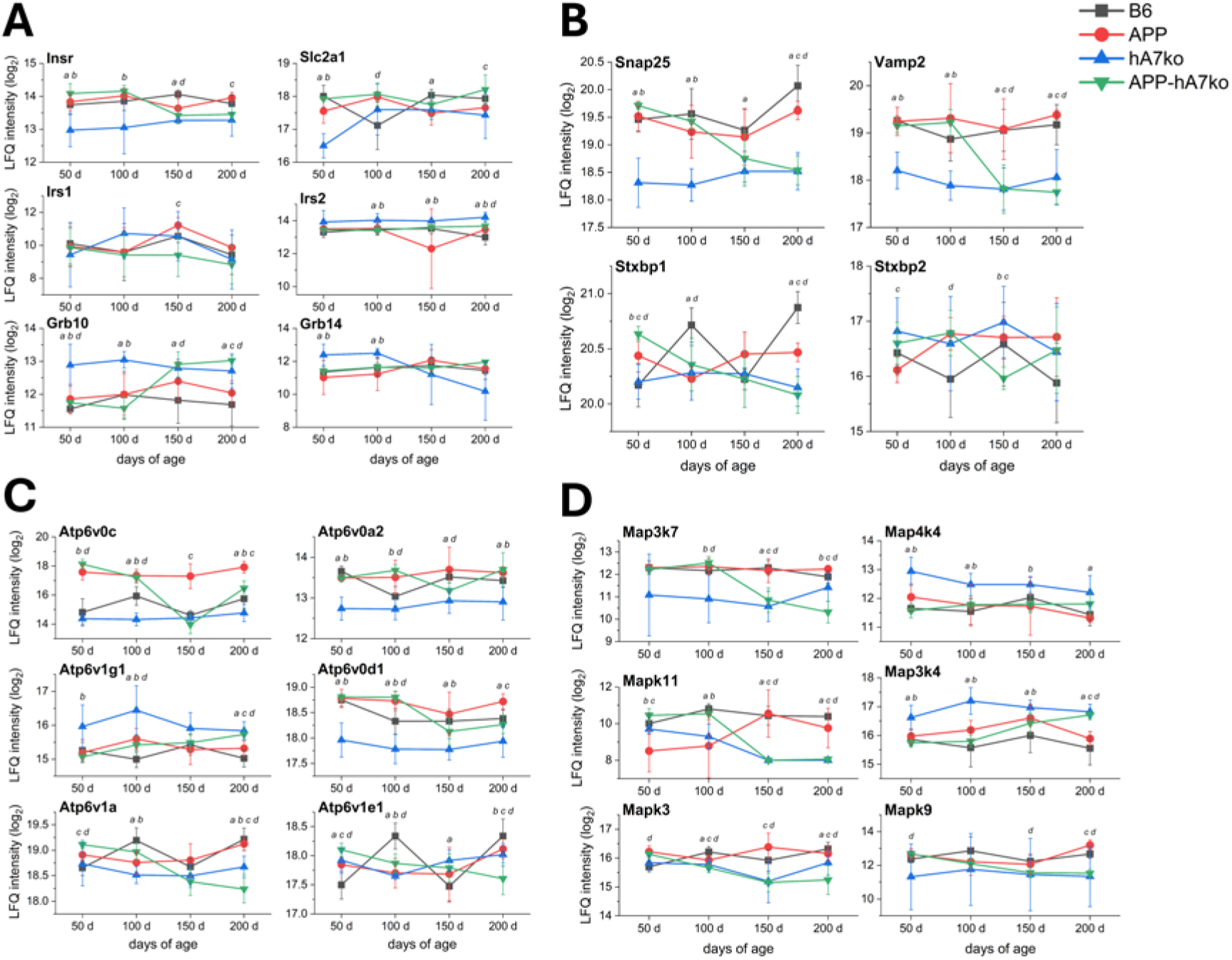
Age-dependent LFQ intensity profiles of insulin signaling, vesicle exocytosis and acidification, and MAPK pathway proteins across genotypes. Mean LFQ intensities (*±* SD) for B6 (grey), APP (red), hA7ko (blue), and APP-hA7ko (green) mice are plotted at 50, 100, 150, and 200 days for each protein, with lines connecting time points to visualize age-dependent changes. The figure shows representative proteins that were recurrently identified as DAPs across multiple comparisons and are associated with key biological functions and processes identified by manual curation and GO enrichment analysis. (A) Core insulin signaling pathway components. (B) Proteins involved in vesicle exocytosis and synaptic vesicle release. (C) Proteins associated with vesicle lumen acidification and V-type proton ATPase function. (D) MAPK pathway proteins related to stress and inflammatory signaling. Statistically significant differences (p *≤* 0.05; two-tailed Welch’s t-test) for the comparisons performed are indicated by letters at each age: a, B6 vs. hA7ko; b, hA7ko vs. APP-hA7ko; c, APP vs. APP-hA7ko; d, B6 vs. APP-hA7ko.

Several characteristic temporal patterns were observed. For some proteins, *Abca7* deficiency was associated with an early and persistent shift: hA7ko mice already differed from the other genotypes at young ages and remained distinct at older ages as well (e.g., RELA, CTSD, ATP6V0A2). For other proteins, hA7ko mice separated already at early ages and APP-hA7ko mice later converged toward the same profile, while B6 and APP mice tended to cluster together (e.g., IL1R1, NFKBIB, NEDD4, XIAP, SNAP25, VAMP2, GRB10, MAPK11). This pattern illustrates an agedependent proteomic separation of *Abca7* -deficient vs. *Abca7* -expressing brains. Another set of proteins showed little or no genotype separation at 50-100 days, followed by a more evident divergence at 150-200 days. This was the case, for example, for complement components and microglial receptors (C1QA, C1QC, TREM2), phosphoinositide regulators (INPP5D, PIP5K1B), and ubiquitin-proteasome-related proteins (PSMB7, UBE2V1), reflecting the late emergence of brain proteomic differences among the investigated genotypes. In contrast, a small number of proteins (e.g., AKT2, BCL2) displayed converging trajectories, with initial genotype differences that attenuated by 200 days, indicating early effects that become partially normalized over time. Finally, some proteins (e.g., PIK3CA, PIK3CB, PTEN, NOS3, STXBP1, STXBP2, ATP6V1E1) showed more complex or fluctuating changes that did not fit clear patterns, likely reflecting secondary responses to upstream molecular alterations. Overall, the recurrent identification of these proteins across multiple comparisons, together with their manual functional annotation and GO enrichment analysis (see sections 3.3-3.6), provides representative molecular markers of the major biological processes underlying the brain proteomic response to *Abca7* hypofunction, A*β* pathology, and their interaction. These include immune and complement signaling, insulin and PI3K/AKT signaling, the ubiquitin-proteasome system, vesicle exocytosis and acidification machinery, and MAPK signaling (Figs. 11 and 12).

To place these findings in the context of human AD, we compared our DAP lists from all 16 genotype-age comparisons with the NeuroPro database, a curated resource integrating more than 8000 proteins reported as altered in the human brain due to AD across 55 proteomic studies. For each comparison and age, we quantified the fraction of DAPs not included in NeuroPro, i.e., proteins that have not been reported in those human AD datasets and thus represent novel candidates in this context. Averaged over the four ages, the proportion of such novel DAPs was 15.5% for B6 vs. hA7ko (range 13.0–18.5%), 16.5% for hA7ko vs. APP-hA7ko (12.1–28.6%), 26.1% for APP vs. APP-hA7ko (15.5–35.7%), and 12.1% for B6 vs. APP-hA7ko (5.6–16.3%). These novel DAPs are highlighted in the Supplementary Material (Table S3). The highest proportion of novel proteins was thus observed for APP vs. APP-hA7ko, indicating that *Abca7* loss on an amyloidogenic background leads to a brain proteomic response that is only partly captured by existing human AD datasets. At the same time, the overlapping fraction of DAPs that are present in NeuroPro (approximately 75–85%) demonstrates substantial consistency between our mouse results and human AD brain proteomics, highlighting the translational value of both the mouse models and the applied analytical workflow. Together, these overlaps and novel findings suggest that *Abca7* hypofunction and A*β* pathology converge on core immune, insulin, vesicle, ubiquitin-proteasome, and MAPK pathways that are shared with human AD, while also highlighting additional candidate proteins and modules that may refine our understanding of how *Abca7* contributes to AD-related brain pathology.

## 4. Discussion

### 4.1. Major findings and general interpretation of the brain proteomic results

In the present study, we used a systematic comparative analysis of mouse brain proteomic data to investigate the protein-level consequences of *Abca7* deficiency in the presence and absence of A*β* pathology across four ages, with a specific focus on insulin signaling- and neuroinflammation-related processes. By combining quantitative proteomics with functional annotation, GO enrichment, PPI network analysis, subcellular localization analysis, and comparison with human AD brain proteomic data, we obtained an integrated view of the molecular changes associated with *Abca7* deficiency and A*β* pathology. Although the number and identity of DAPs varied substantially among genotype comparisons and ages, several consistent molecular processes emerged. The principal changes involved inflammatory pathways, canonical insulin receptor and PI3K/AKT signaling, phosphoinositide and MAPK signaling, the ubiquitin-proteasome system, and vesicle exocytosis and acidification. Proteins associated with the plasma membrane were preferentially enriched, and approximately 80% of the identified DAPs overlapped with proteins previously reported in human AD brain proteomic studies. Together, these findings indicate that *Abca7* deficiency and its interaction with A*β* pathology are associated with age-dependent remodeling of interconnected brain protein networks rather than isolated alterations in individual proteins.

Several general features of comparative proteomics should be considered when interpreting these findings. The identified DAPs are unlikely to represent only the direct molecular consequences of the genetic perturbations, but instead reflect a proteomic phenotype established by primary effects, downstream pathway changes, and homeostatic or adaptive responses. The mammalian brain maintains tissue function through coordinated regulation of gene expression, protein synthesis and degradation, metabolism, and intercellular signaling (Gräff et al., 2011; Ding et al., 2007; Camandola and Mattson, 2017; Shin et al., 2026). Consequently, the number and magnitude of proteomic differences should not be interpreted solely as measures of pathological severity. They represent the net outcome of *Abca7* deficiency, with or without concomitant A*β* pathology, and compensatory mechanisms that establish a new molecular state. This systems-level perspective is particularly relevant to chronic genetic models, in which constitutive modifications may permit extensive proteomic adaptation before tissue collection, including at the earliest age examined here.

### 4.2. Comparative analysis: effects of Abca7 deficiency, aging, and Aβ pathology

The four pairwise comparisons provided complementary perspectives on the overall APP-hA7ko phenotype. The B6 versus hA7ko comparison reflected the effects of *Abca7* deficiency without A*β* pathology and revealed numerous DAPs already at young ages, including proteins associated with oxidative stress, protein phosphorylation, and immune responses. GO enrichment of PPI network clusters also indicated altered transmembrane fatty acid transport, consistent with established ABCA7 functions in lipid transport and phagocytosis (Aikawa et al., 2018). The hA7ko versus APP-hA7ko comparison isolated the additional influence of APP expression and A*β* pathology in the absence of functional ABCA7 protein, identifying changes related to oxidative stress, immune and defense responses, and insulin receptor signaling. These findings are consistent with the established associations of A*β* pathology with oxidative stress and membrane damage, innate immune activation, and altered cerebral glucose metabolism (Cheignon et al., 2018; Halle et al., 2008; Dewanjee et al., 2022). The APP versus APP-hA7ko comparison revealed the impact of *Abca7* deficiency on an amyloidogenic background and contained the highest proportion of proteins not represented in NeuroPro. Its DAPs were predominantly associated with apoptosis and cellular immune responses, suggesting that *Abca7* deficiency may enhance processes implicated in AD-related neurodegeneration (Kumari et al., 2023). Finally, B6 versus APP-hA7ko captured the combined effects of *Abca7* deficiency and APP expression, revealing extensive changes in cellular stress responses, inflammatory activation, and intracellular signal transduction. Despite their distinct biological contexts, the four comparisons repeatedly converged on a limited group of interconnected pathways, demonstrating that the comparative design provides information that could not be obtained from any single comparison alone.

### 4.3. Age-dependent remodeling of insulin signaling- and neuroinflammation-related networks

Because aging is the principal established risk factor for AD (Guerreiro and Bras, 2015; Munoz and Feldman, 2000), brain samples from mice at four ages were examined to characterize temporal changes. DAP numbers followed distinct age-dependent patterns across comparisons. No monotonic trend was observed for B6 versus hA7ko or hA7ko versus APP-hA7ko, whereas DAP numbers increased progressively with age in APP versus APP-hA7ko and, most prominently, B6 versus APP-hA7ko. This pattern may reflect the increasing contribution of A*β* pathology superimposed on agerelated molecular changes. The investigated ages span the progressive development of amyloidosis in APPPS1-21 mice, in which the first cerebral A*β* deposits appear at approximately 6–8 weeks and subsequently increase with age (Radde et al., 2006). Consistently, PCA showed progressively clearer genotype separation, particularly between *Abca7* -deficient and *Abca7* -expressing groups at 150 and 200 days. The affected biological processes also varied across ages, indicating that the proteomic effects of *Abca7* deficiency are dynamic rather than static. Longitudinal profiles of representative proteins further showed that *Abca7* deficiency was frequently associated with early divergence of protein abundance trajectories, whereas A*β* pathology progressively enhanced or modified these differences. These temporal patterns suggest that molecular disturbances associated with *Abca7* deficiency may precede more pronounced metabolic, inflammatory, and neurodegenerative changes and subsequently interact with A*β* pathology to reshape downstream cellular responses.

### 4.4. Mechanistic interpretation of the proteomic changes

The study was based on the hypothesis that ABCA7 may contribute to the intersection between neuroinflammation and impaired brain insulin signaling. ABCA7 has established roles in lipid metabolism and immune function, and emerging evidence links its dysfunction to altered cellular energy metabolism (Dib et al., 2021; Kawatani et al., 2024; Nowyhed et al., 2017). Based on the present proteomic findings, altered lipid homeostasis represents a plausible upstream process connecting ABCA7 deficiency to these molecular systems. ABCA7 hypofunction may disturb cellular lipid transport and membrane lipid composition, thereby modifying the lipid environment and lipid–protein interactions of membrane-associated signaling proteins. Membrane lipid composition can influence the conformation, interactions, and activity of integral membrane proteins, providing a mechanistic basis for this hypothesis (Bernhardt et al., 2025; Yang et al., 2024). Consistent with this interpretation, plasma membrane- and endosome-associated proteins were enriched among the DAPs, including the insulin receptor, interleukin receptors, ion channels, and several phosphoinositide-metabolizing enzymes. Altered membrane lipid–protein interactions could therefore affect signal transduction and propagate through intracellular networks that regulate metabolism, immune responses, vesicle trafficking, and protein homeostasis. This sequence should be regarded as a data-driven hypothesis rather than a mechanism directly demonstrated by abundance-based proteomics.

Independent studies support several components of this model. ABCA7 deficiency has been shown to alter neuronal phosphatidylcholine and sphingomyelin profiles in association with mitochondrial dysfunction, oxidative stress, and synaptic abnormalities (Kawatani et al., 2024). Analyses of human *ABCA7* loss-of-function variants similarly identified altered phosphatidylcholine metabolism and perturbations of mitochondrial, proteostatic, and synaptic pathways in neurons (von Maydell et al., 2025). Neuronal *Abca7* deficiency has also been reported to alter brain and synaptosomal lipid profiles and to exacerbate mitochondrial dysfunction, A*β* pathology, and neuronal injury in an amyloidogenic mouse model (Wang et al., 2025). Complementary evidence supports an immune component: ABCA7 modulates neuroinflammation through the NLRP3 inflammasome in AD mice (Santos-Garcia et al., 2025), the AD-associated ABCA7 p.A696S protein variant modifies the microglial response to amyloid pathology (Ma et al., 2025), and recent single-cell and experimental analyses have linked *Abca7* -related microglial responses to the interaction between type 2 diabetes and AD (Cheng et al., 2026). The present changes in PI3K/AKT, phosphoinositide and MAPK signaling, vesicle biology, and the ubiquitin-proteasome system therefore support a model in which ABCA7 dysfunction affects multiple interacting molecular systems rather than a single linear pathway.

### 4.5. Limitations

Several limitations should be considered. First, whole-brain homogenates provide broad proteomic coverage but prevent assignment of the observed changes to specific brain regions or cell types. Given the distinct contributions of neuronal and glial populations to AD pathogenesis, region-, cell type-, and spatially resolved approaches will be required to localize the identified alterations (Mathys et al., 2024). Second, protein abundance does not necessarily reflect protein activity, which is additionally regulated by post-translational modifications, protein interactions, and subcellular localization (Alganem et al., 2022; Zhong et al., 2023). The affected pathways should therefore be interpreted as changes in pathway-associated protein abundance rather than direct evidence of altered signaling activity or insulin resistance. Third, the analysis focused on predefined insulin signaling and neuroinflammation protein sets, increasing sensitivity for the study hypothesis while excluding unrelated networks. Fourth, constant-value imputation may influence estimates for low-abundance proteins, and DAP screening used nominal p-value and fold-change thresholds without adjustment for multiple testing; individual DAPs should therefore be interpreted together with recurrent comparison patterns, pathway enrichment, and network-level evidence. Fifth, the sample sizes were not designed for adequately powered sex-stratified analyses. Although no clear sex-related clustering was observed, biological responses in APP-transgenic mice can depend on sex and disease stage (Bascunana et al., 2023). Sixth, APP-transgenic models reproduce major features of cerebral A*β* pathology but do not capture the full complexity of human AD (Radde et al., 2006; Kitazawa et al., 2012). Amyloid deposition and glial responses may also be influenced by genomic background and endogenous mouse *App* expression (Frohlich et al., 2013; Steffen et al., 2017). Finally, the present study is descriptive and hypothesis-generating; the proposed links among ABCA7 deficiency, lipid homeostasis, insulin signaling, neuroinflammation, and A*β* pathology require direct experimental validation.

### 4.6. Conclusions and future perspectives

In conclusion, this study provides a systematic, age-resolved proteomic characterization of molecular changes associated with *Abca7* deficiency in the presence and absence of A*β* pathology, with particular emphasis on insulin signaling and neuroinflammation. The findings indicate dynamic remodeling of interconnected protein networks and support a model in which altered lipid homeostasis contributes to broader changes in membrane-associated signaling, cellular metabolism, immune responses, vesicle biology, and proteostasis. The substantial overlap between the identified DAPs and proteins reported in human AD brain proteomic studies supports the translational relevance of these alterations, while the comparison-specific proteins provide candidates for further investigation. Nevertheless, the proposed relationships should not be interpreted as demonstrated causal mechanisms.

Future studies should combine targeted functional experiments with phosphoproteomics, lipidomics, and cell typeand brain region-resolved approaches. Measurements of insulin responsiveness, endolysosomal function, glial activation, proteostasis, synaptic activity, and mitochondrial metabolism will be needed to establish the sequence and functional consequences of the identified changes. Genetic or pharmacological restoration of ABCA7 function would provide a direct test of causality, while adequately powered cohorts should assess sex-dependent effects. Pharmacological modulation and structure-guided ligand discovery may also help evaluate ABCA transporters as diagnostic or therapeutic targets (Namasivayam et al., 2021; Pahnke et al., 2021). Such studies may clarify how ABCA7 hypofunction contributes to AD development and progression and identify molecular processes with therapeutic relevance.

Given the central role of ABCA7 in lipid transport and membrane homeostasis, defining the metabolic and lipidomic consequences of ABCA7 deficiency or dysfunction should be a major priority for future research. Mass spectrometry imaging (MSI) is particularly well suited to this objective because it maps lipids and metabolites directly in intact tissue while preserving anatomical and pathological context. In transgenic AD mouse models, multimodal MSI has resolved distinct ganglioside, phosphoinositol, ceramide, lysophospholipid, and other lipid distributions associated with individual A*β* plaques, plaque maturation, and structural plaque polymorphism (Michno et al., 2022; Ge et al., 2023). Plaque-resolved MSI of postmortem familial AD brain has likewise identified extensive local enrichment and depletion of sphingolipid and phospholipid species, including lipid patterns that differed from those observed in mouse models (Michno et al., 2024). Applying such spatial neurolipidomic approaches to *Abca7* -deficient brains could determine whether the proteomic changes reported here coincide with region- or plaque-associated lipid microenvironments. Integration with high-resolution visualization, explainable machine learning, quantitative proteomics, conventional lipidomics, histopathology, and functional experiments may therefore clarify how ABCA7-dependent lipid disturbances propagate across brain regions and contribute to metabolic dysfunction, neuroinflammation, and A*β* pathology (Gildenblat and Pahnke, 2026; Gildenblat et al., 2026).

## Acknowledgements

The mass spectrometry-based measurements of mouse brains were performed at the Proteomics Core Facility at the University of Oslo/Oslo University Hospital, which is supported by the Core Facilities Program of the South-Eastern Norway Regional Health Authority (HSØ) and NAPI (www.napi.uio.no, NFR, Norway; 295910).

## Data and materials availability

The proteomics dataset is available through the PRIDE Proteomics Identifications Database under accession PXD053250 (Santos-Garcia et al., 2025). No new biological materials were generated in this study.

## Funding

D.M. received funding from Nasjonalforeningen for folkehelse (ProjectID: 52219). J.P. received funding from Nasjonalforeningen for folkehelse (Demensforskningsprisen 2025, Norway), Norges forskningsråd (NFR, Norway; 327571 (PETABC), 295910 (NAPI)), Helse Sør-Øst (Norway, 2022046), and the EIC Pathfinder Open Challenges program (European Commission; OPTIPATH 7D, 101185769). The funders had no role in study design; data collection, analysis, or interpretation; preparation of the manuscript; or the decision to submit the work for publication.

## CRediT authorship contribution statement

Conceptualization (J.P.), Methodology (D.M., J.P.), Validation (D.M., J.P.), Formal analysis (D.M.), Investigation (D.M., J.P.), Resources (J.P.), Data curation (D.M., J.P.), Writing–original draft (D.M., J.P.), Writing–review and editing (D.M., J.P.), Supervision (J.P.), Project administration (D.M., J.P.), Funding acquisition (D.M., J.P.).

## Declaration of competing interest

The authors declare no competing interests.

## Declaration of generative AI and AI-assisted technologies in the manuscript preparation process

During the preparation of this work, the authors used PrismAI, an AI-assisted writing and editing environment powered by OpenAI language-model technology, to organize source materials and improve manuscript structure, language, readability, and LaTeX formatting. After using this tool, the authors reviewed, edited, and verified the content as needed and take full responsibility for the content of the published article. No generative AI tool was used to create or alter the research data or figures.

## Supplemetary Materials

- Supplementary Material 1: Figures S1–S12 and Table S5.
- Supplementary Material 2: Excel file with Tables S1–S4.

---

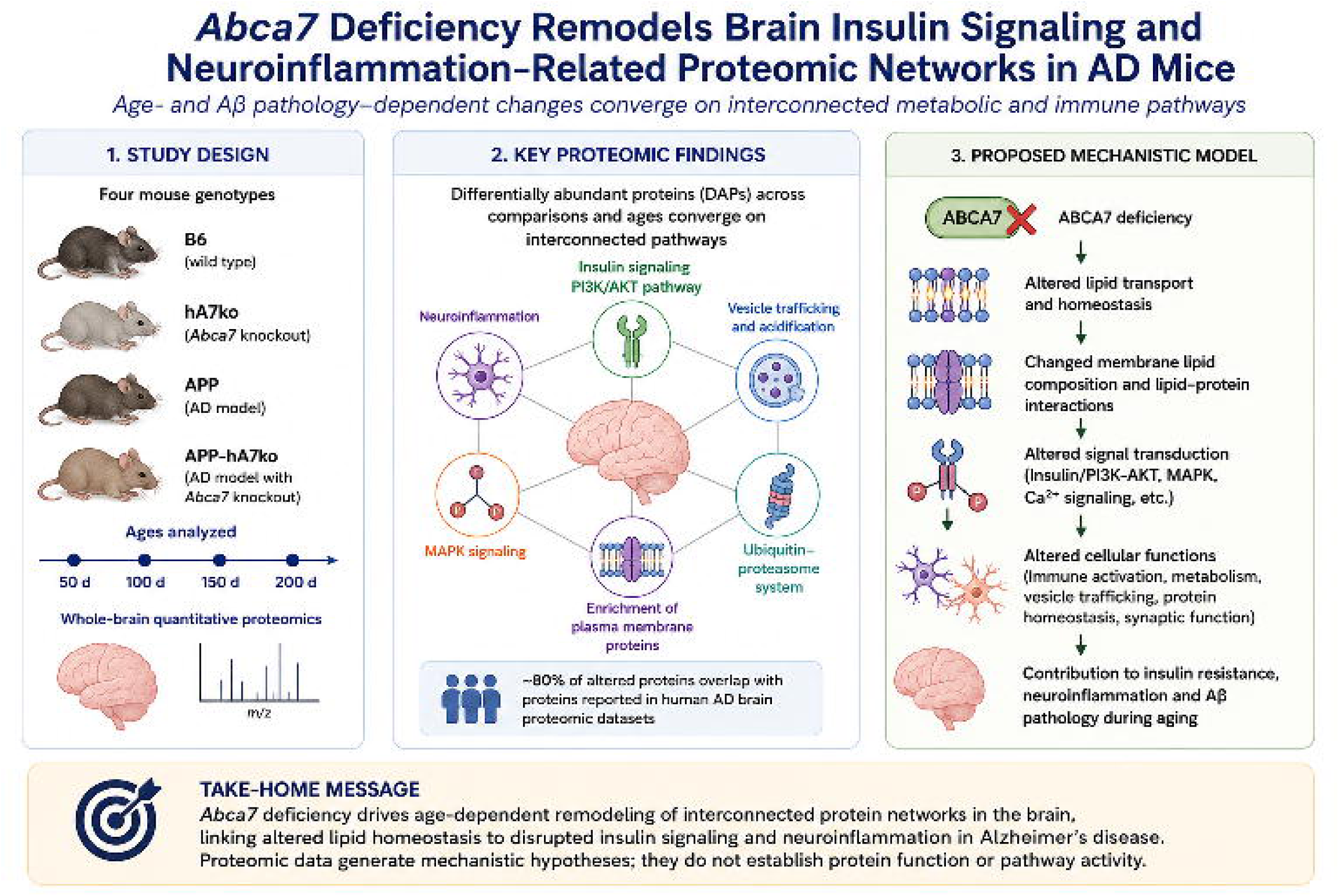

## Highlights

- *Abca7* deficiency remodels brain proteomes in an age- and context-dependent manner.
- Early changes precede extensive amyloid-associated proteome remodeling.
- Insulin/PI3K-AKT and immune networks converge with vesicle and MAPK pathways.
- Altered proteins are enriched at membranes, consistent with ABCA7 lipid transport.
- About 80% of altered mouse proteins overlap with human AD proteomic datasets.

## Notes

### Competing Interest Statement

The authors have declared no competing interest.

