## Supplementary material for "Age-dependent brain proteome remodeling links *Abca7* deficiency to insulin signaling and neuroinflammation in Alzheimer’s disease mice": Suppl Material 1

---

**This PDF includes:**

- Supplementary Figures S1–S12: proteomics quality-control analyses, PCA scree plots, subcellular compartment enrichment analysis, and clustered protein–protein interaction networks.
- Supplementary Table S5: an alphabetical list of the gene symbols used in the manuscript and their corresponding full names.

*, www.pahnkelab.eu

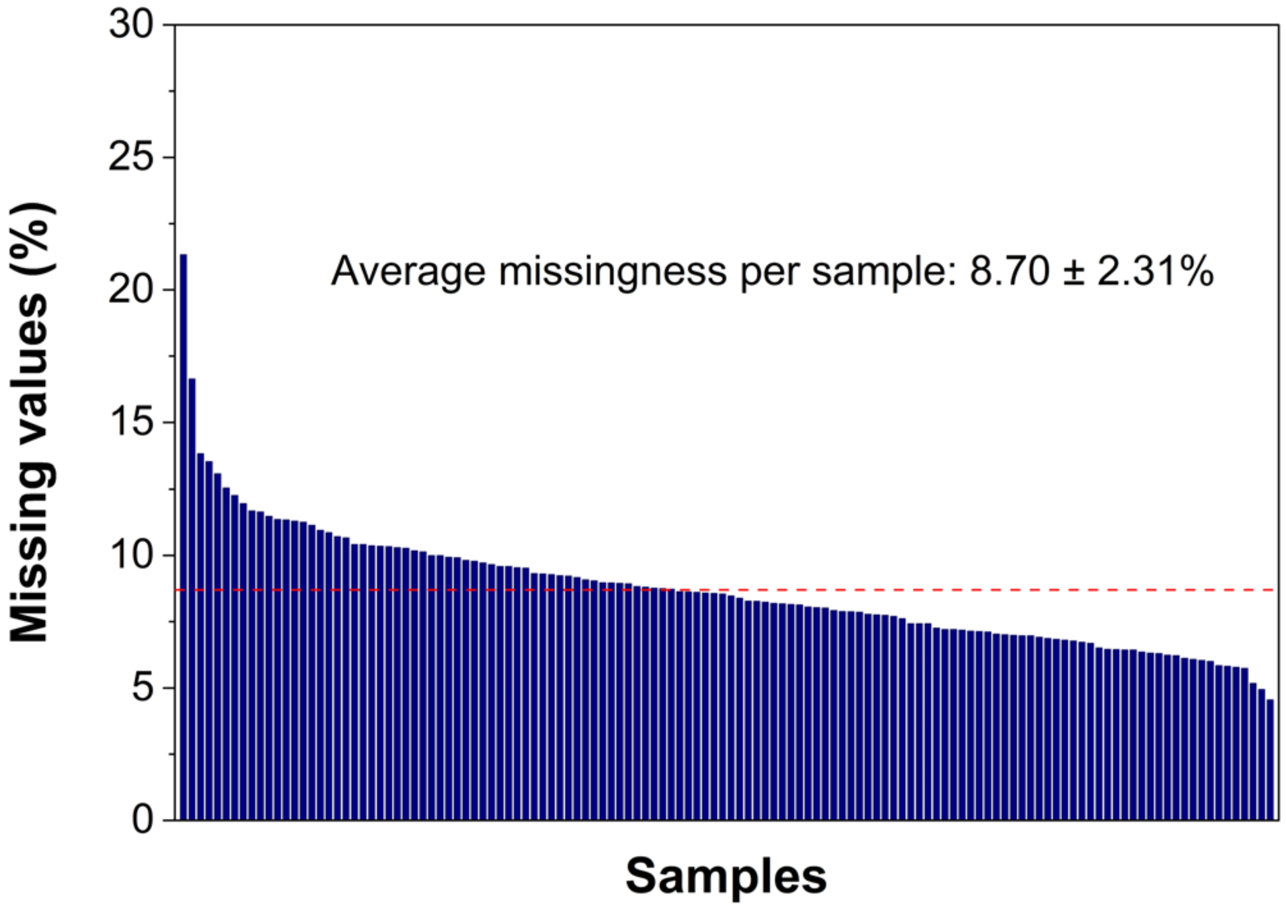


Figure S1: Missingness per sample in the quantitative proteomics dataset. For each brain sample (represented by a column), the proportion of missing LFQ intensity values across all quantified proteins is shown, illustrating generally low levels of missingness (ranging from 4.54% to 21.32%) and the absence of samples with excessive missing values. The average per-sample missingness equals the global missingness across the entire dataset (8.70%, shown by the red dashed line), which contains 128 whole-brain samples and 7636 identified proteins.

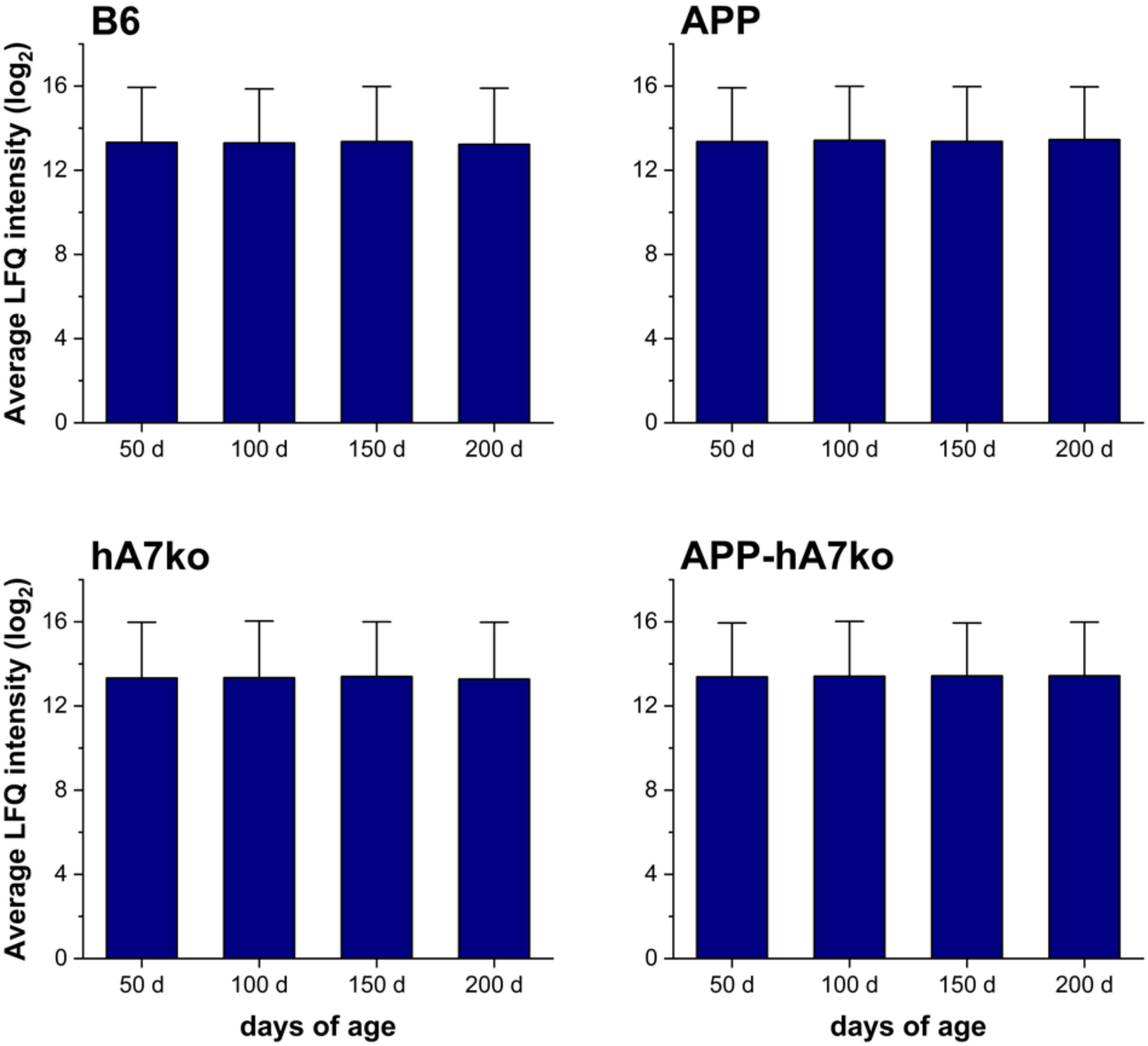


Figure S2: Distribution of average LFQ intensities across all proteins and experimental groups. For each genotype-age group, the mean LFQ intensity across all 7636 quantified proteins is shown, indicating very similar overall signal intensities between groups and the absence of major global intensity shifts that could indicate systematic technical bias. Average protein LFQ intensities in the investigated groups were as follows: B6 50 days: 13.32 ± 2.62; B6 100 days: 13.30 ± 2.57; B6 150 days: 13.36 ± 2.62; B6 200 days: 13.23 ± 2.67; APP 50 days: 13.36 ± 2.56; APP 100 days: 13.42 ± 2.57; APP 150 days: 13.37 ± 2.61; APP 200 days: 13.46 ± 2.51; hA7ko 50 days: 13.33 ± 2.65; hA7ko 100 days: 13.35 ± 2.69; hA7ko 150 days: 13.40 ± 2.60; hA7ko 200 days: 13.28 ± 2.70; APP-hA7ko 50 days: 13.39 ± 2.56; APP-hA7ko 100 days: 13.41 ± 2.61; APP-hA7ko 150 days: 13.43 ± 2.52; APP-hA7ko 200 days: 13.43 ± 2.55 (data provided as mean ± SD).

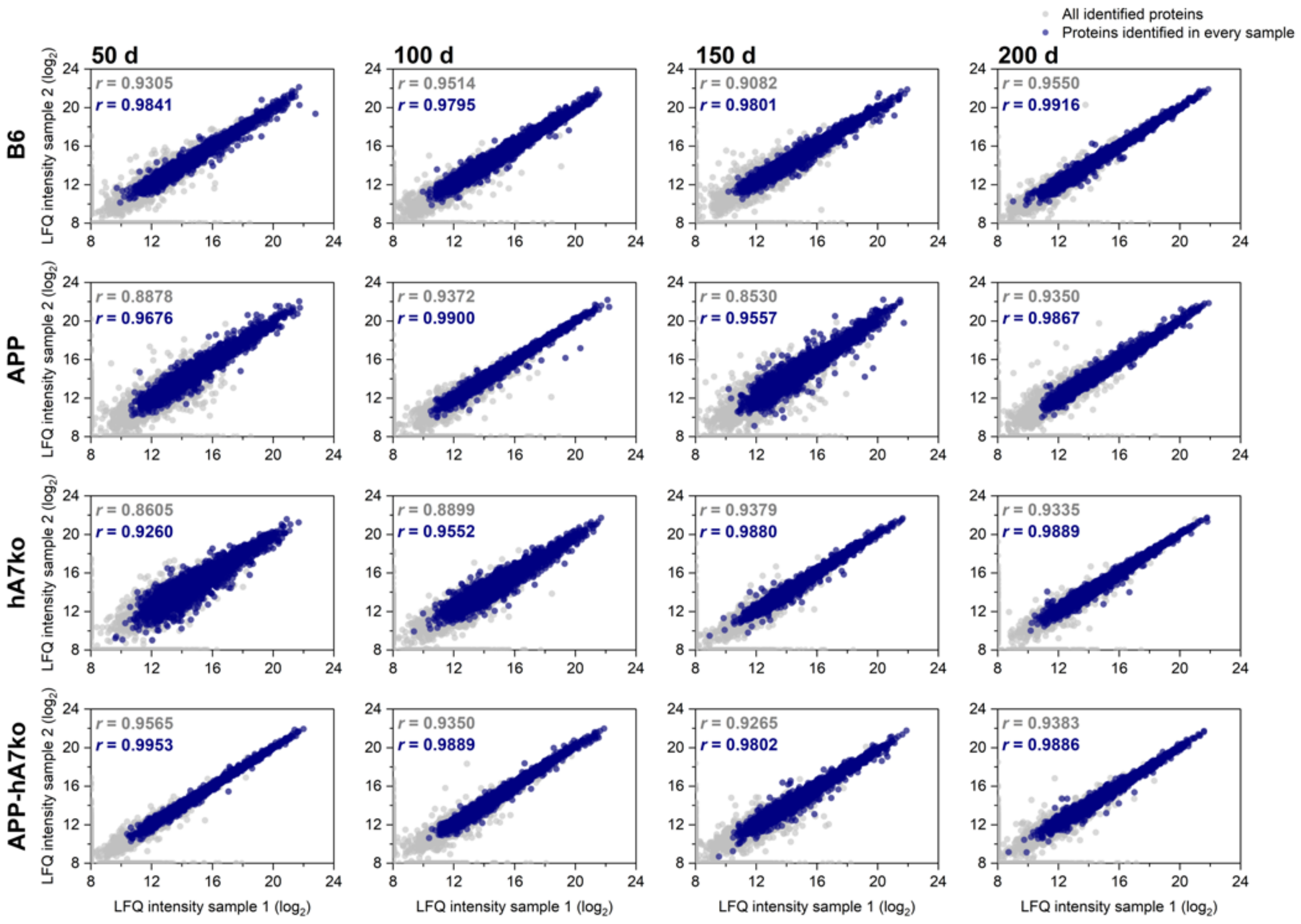


Figure S3: Pairwise correlation plots of LFQ intensities between biological replicates. For each pair of randomly chosen replicate samples within the same genotype-age group, Pearson correlation coefficients (r) of LFQ intensities are shown for all identified proteins (7636 proteins, gray dots) and for proteins quantified in every sample (4774 proteins with 100% data completeness, blue dots). The consistently high correlation values indicate good reproducibility of the quantitative proteomics measurements across replicates. As can be seen, the proteins showing 100% completeness form a more stable part of the cerebral proteome, with higher abundances and lower spread, while the proteins not identified in every sample generally have lower abundances with higher spread.

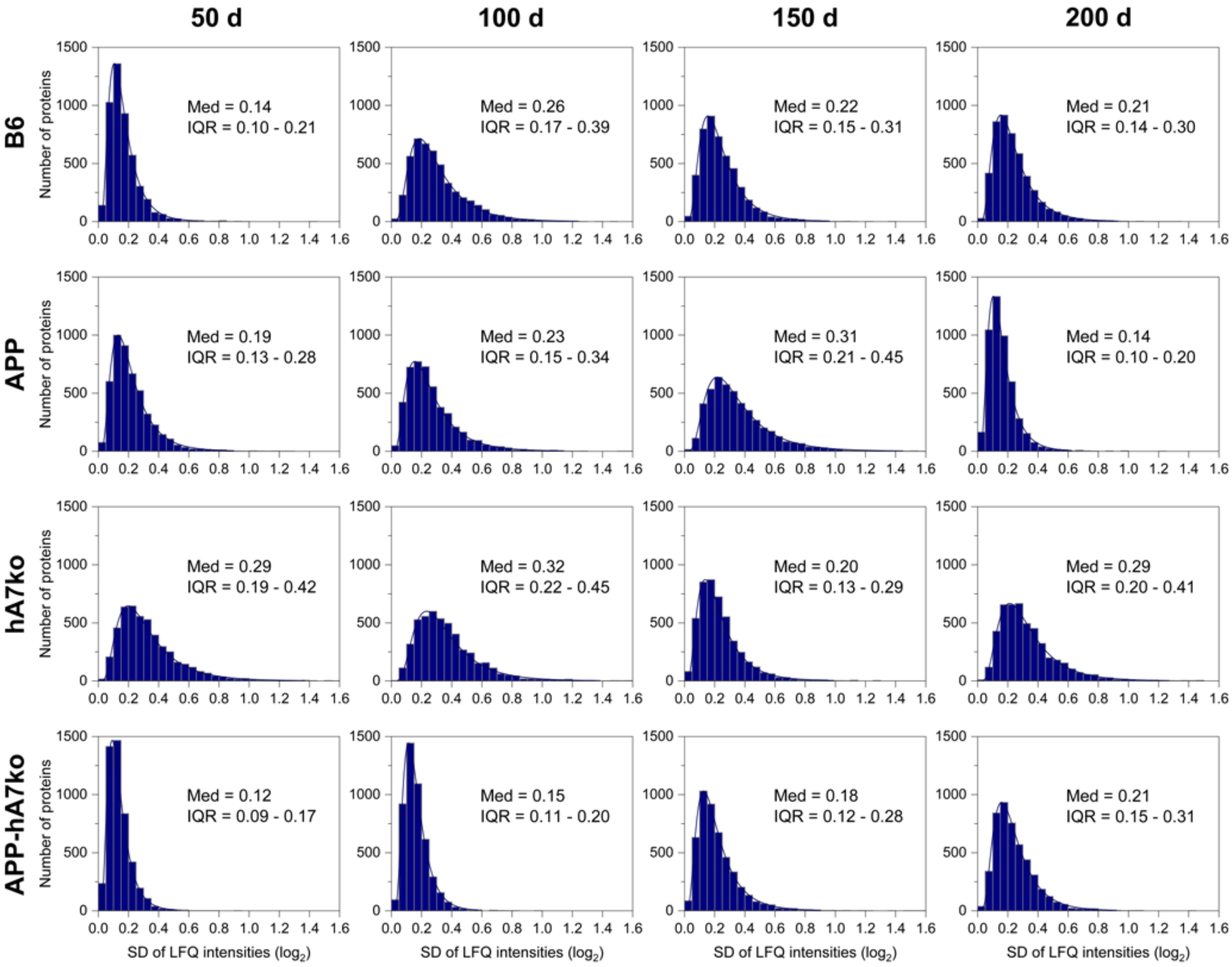


Figure S4: Protein-wise distribution of LFQ intensity variability (SD) across all experimental groups. For each genotype-age group (n = 5–6 mice in each group), the within-group SD was calculated for the 4774 proteins identified in every sample (i.e., showing 100% data completeness). Histograms show the typically right-skewed distribution of SD values of LFQ intensities for individual proteins, illustrating the overall spread of protein-wise variability and supporting the absence of widespread extreme variance that could compromise downstream differential abundance analyses. Log-normal curves are fitted to the histograms, and the median (Med) and interquartile range (IQR) values are shown for each genotype-age group.

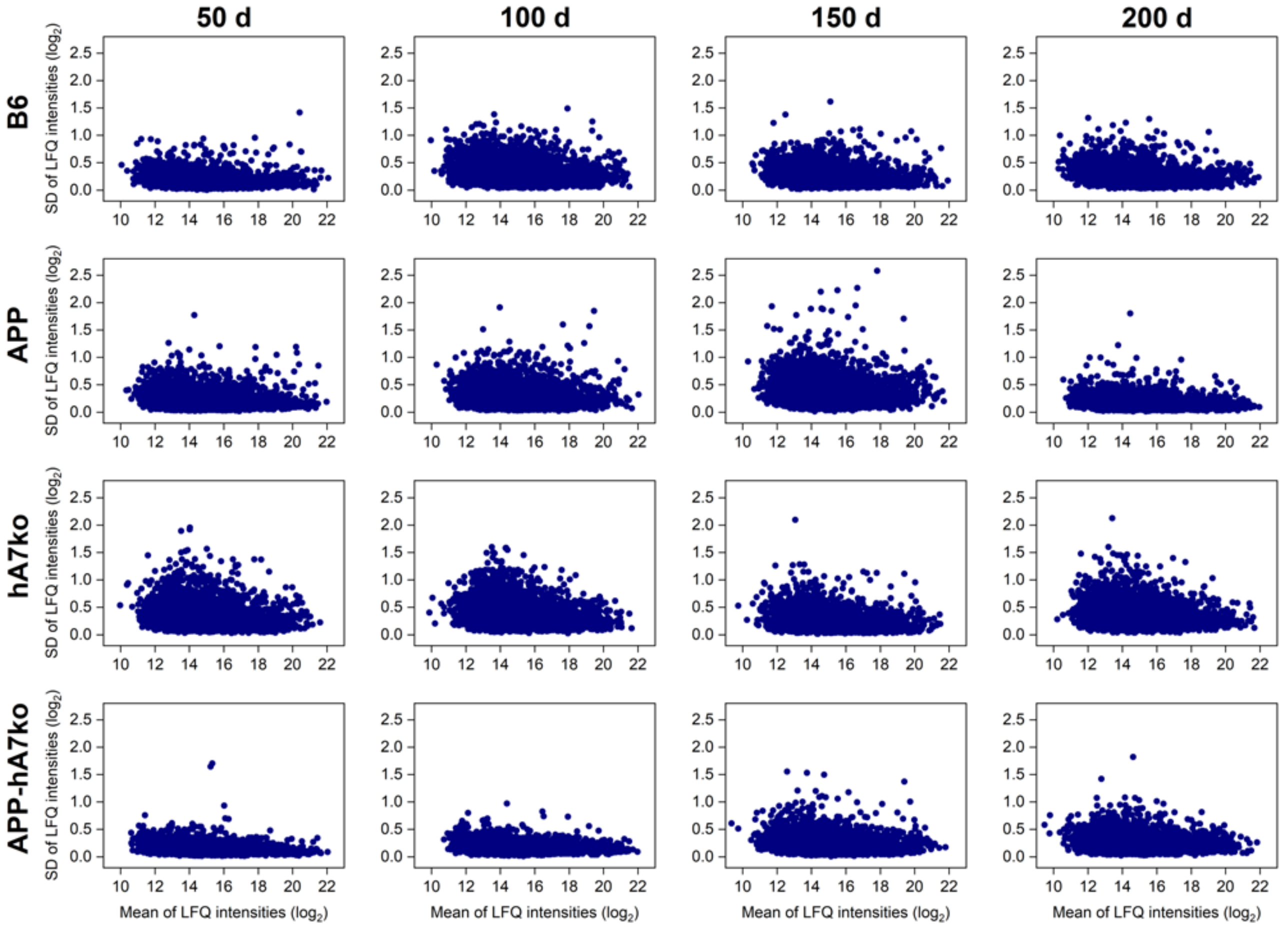


Figure S5: Mean-SD trend plots of protein LFQ intensities across all experimental groups. For each protein with 100% data completeness (4774 proteins), the SD of LFQ intensities is plotted against the mean LFQ intensity within the same genotype-age group (n = 5–6 mice in each group), showing the relationship between signal intensity and variability. In some cases, proteins with lower mean intensities tend to show higher variance; however, no major intensity-dependent changes of SD can be observed (i.e., the quantitative proteomic data show homoscedasticity).

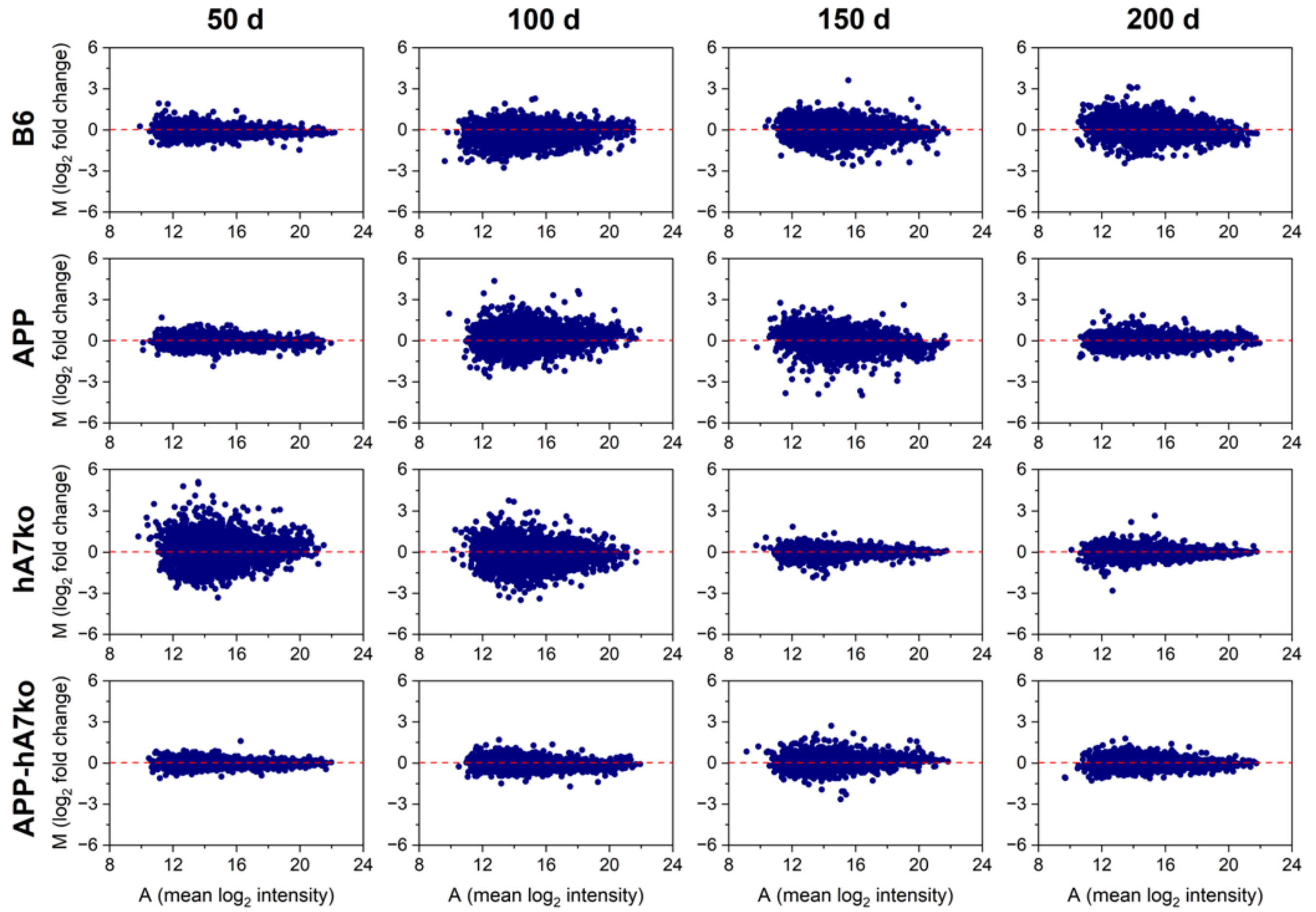


Figure S6: MA plots of protein LFQ intensities for randomly selected replicate pairs. For randomly chosen pairs of biological replicates within the same genotype-age group, MA plots display the difference between the two samples (M = LFQ intensity 1 − LFQ intensity 2) versus their average LFQ intensity (A = (LFQ intensity 1 + LFQ intensity 2) / 2) for all proteins with 100% completeness (4774 proteins). The data points are symmetrically distributed around zero (red dashed line) with no systematic intensity-dependent bias (i.e., showing homoscedasticity), indicating good technical consistency between biological replicates.

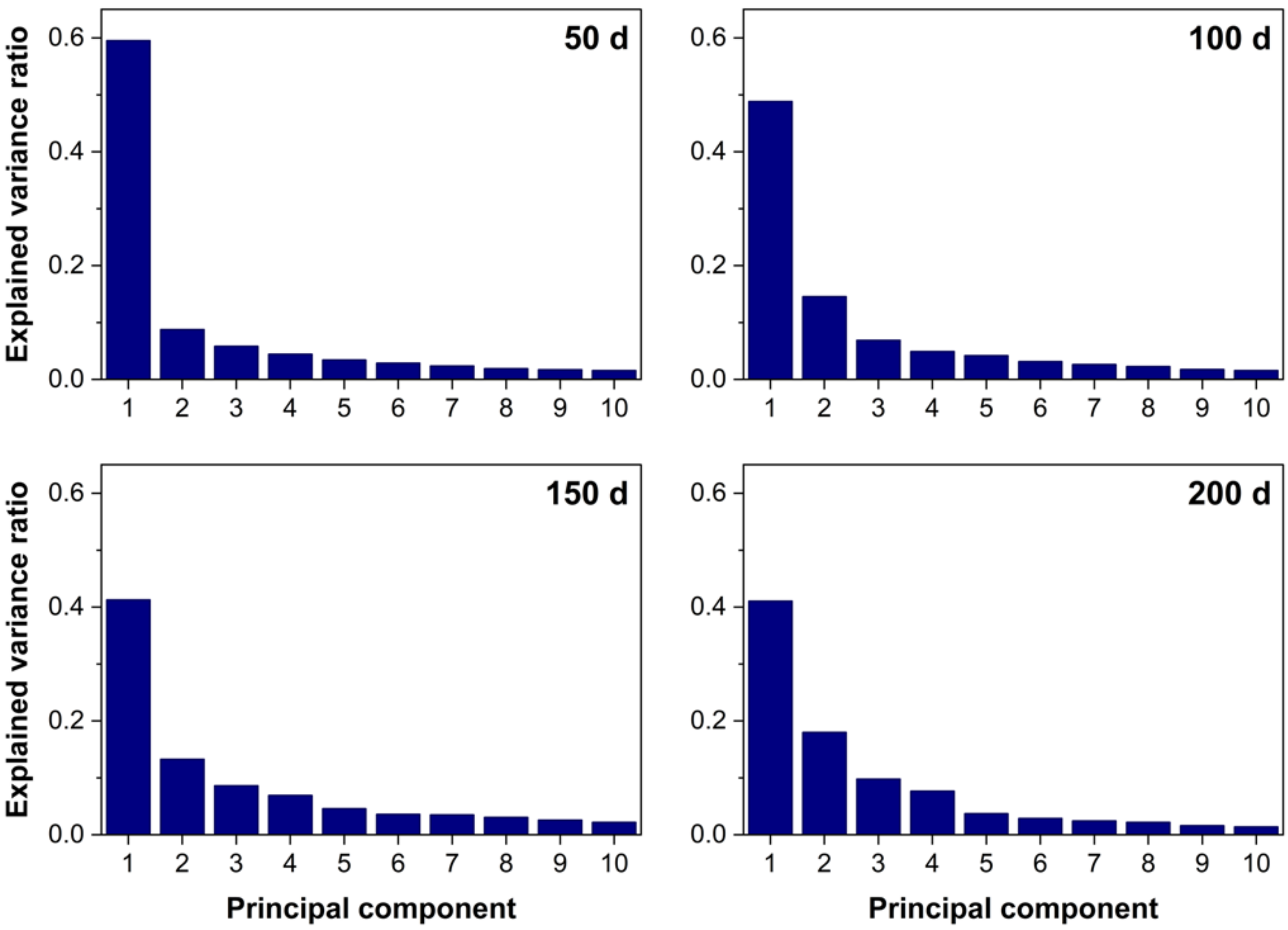


Figure S7: Scree plots of PCA performed on DAPs by genotype at each age. For each age (50, 100, 150, and 200 days), scree plots show the proportion of total variance explained by successive principal components based on LFQ intensity values of differentially abundant proteins. In all cases, the first few components account for a substantial fraction of the variance, supporting the use of the first two principal components to visualize genotype-related separation in the main PCA plots.

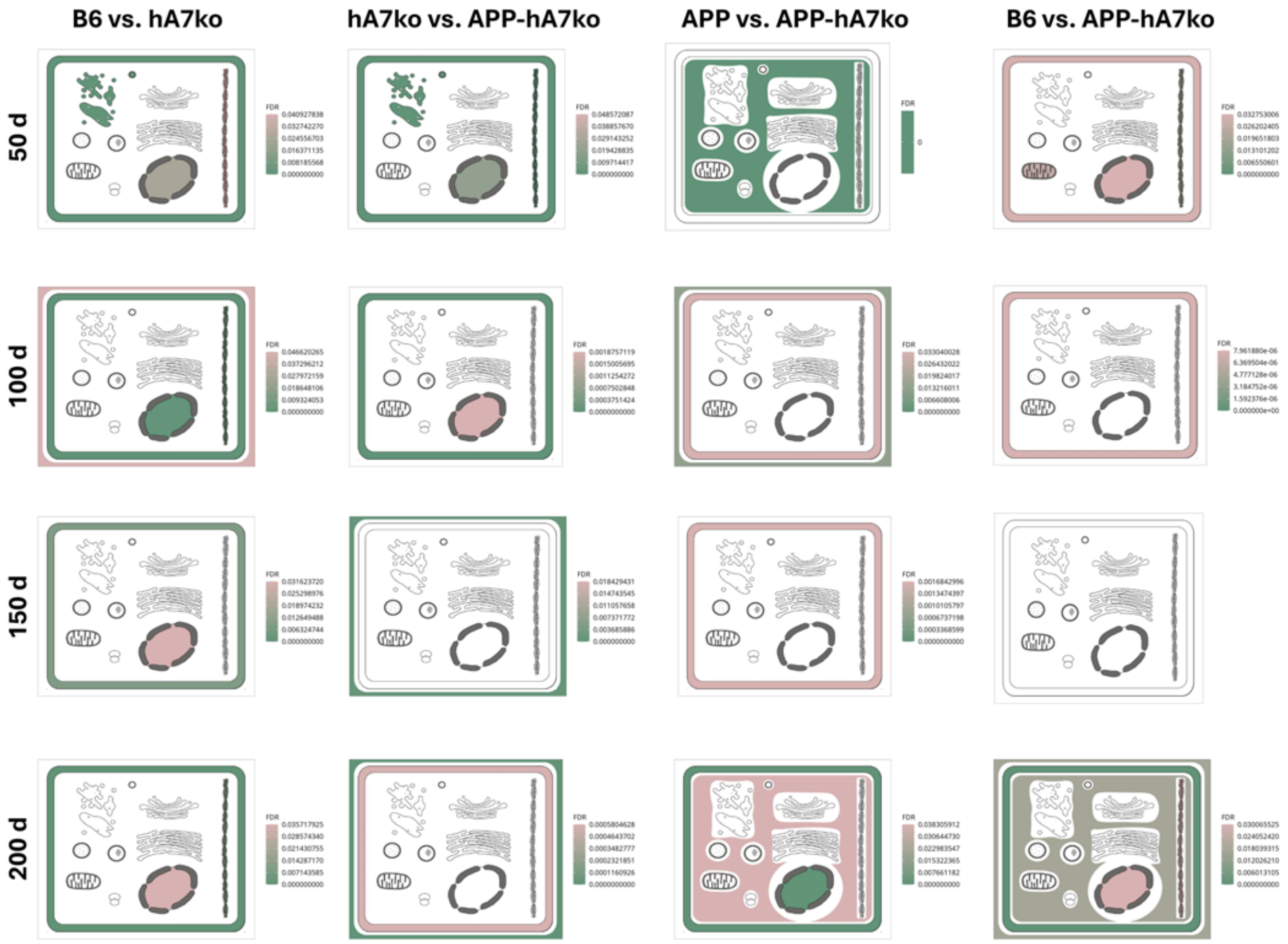


Figure S8: Subcellular compartment enrichment analysis of DAPs across genotype comparisons and ages. Subcellular localization enrichment analysis was performed for DAPs identified in each genotype comparison at 50, 100, 150, and 200 days of age using SubcellulaRVis. SubcellulaRVis calculates protein localization enrichment based on the cellular component aspect of gene ontology (GO) and includes the following subcellular compartments: plasma membrane, endosome, intracellular vesicle, nucleus, cytoskeleton, cytoplasm, vacuole, Golgi apparatus, mitochondrion, endoplasmic reticulum, extracellular region, ribosome, lysosome, and peroxisome. Significantly enriched subcellular compartments are colored according to the false discovery rate (FDR), with green indicating higher statistical significance (lower FDR) and red indicating lower statistical significance (higher FDR). White indicates that no significant enrichment was detected for the corresponding subcellular compartment. The analysis demonstrates comparison- and age-dependent differences in the subcellular distribution of altered proteins, with membrane-associated compartments showing recurrent enrichment across multiple comparisons.

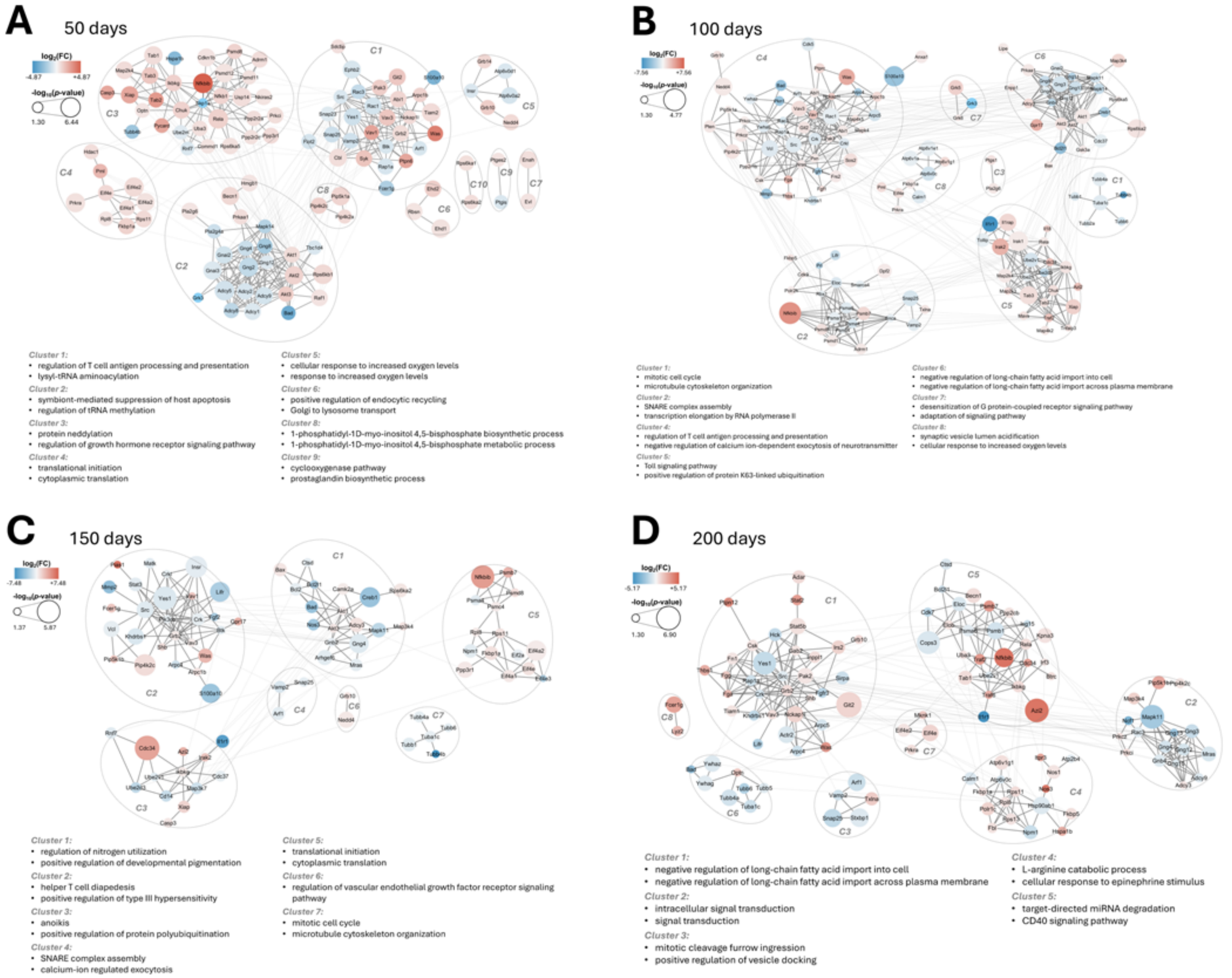


Figure S9: Clustered protein–protein interaction networks of DAPs in B6 vs. hA7ko. Clustered protein–protein interaction networks of DAPs identified in the B6 vs. hA7ko comparison at 50 (A), 100 (B), 150 (C), and 200 (D) days of age. Protein-protein interaction networks were constructed using STRING and visualized in Cytoscape. Node size represents statistical significance ($-\log_{10}(p)$), node fill color indicates the magnitude and direction of protein abundance changes ($\log_2(\text{FC})$), and edges represent predicted functional protein–protein interactions (thicker edges indicate within-cluster interactions, whereas thinner edges indicate between-cluster interactions). Clusters (indicated by ellipses) were identified using the GLay community clustering algorithm and annotated with the top enriched gene ontology (GO) biological process terms. Note that not all clusters showed significant GO term enrichment.

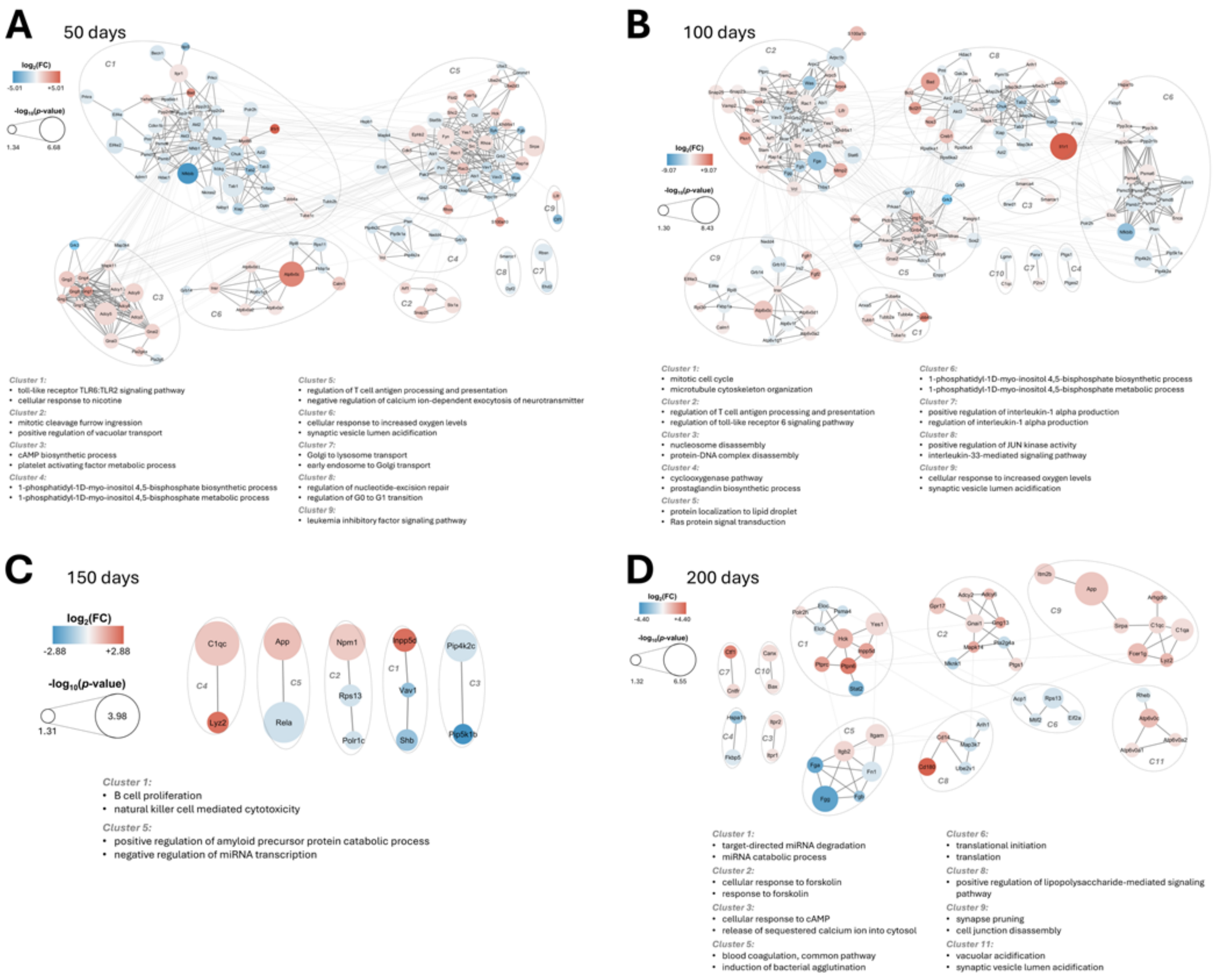


Figure S10: Clustered protein–protein interaction networks of DAPs in hA7ko vs. APP-hA7ko. Clustered protein–protein interaction networks of DAPs identified in the hA7ko vs. APP-hA7ko comparison at 50 (A), 100 (B), 150 (C), and 200 (D) days of age. Protein-protein interaction networks were constructed using STRING and visualized in Cytoscape. Node size represents statistical significance ($-\log_{10}(p)$), node fill color indicates the magnitude and direction of protein abundance changes ($\log_2$(FC)), and edges represent predicted functional protein–protein interactions (thicker edges indicate within-cluster interactions, whereas thinner edges indicate between-cluster interactions). Clusters (indicated by ellipses) were identified using the GLay community clustering algorithm and annotated with the top enriched gene ontology (GO) biological process terms. Note that not all clusters showed significant GO term enrichment.

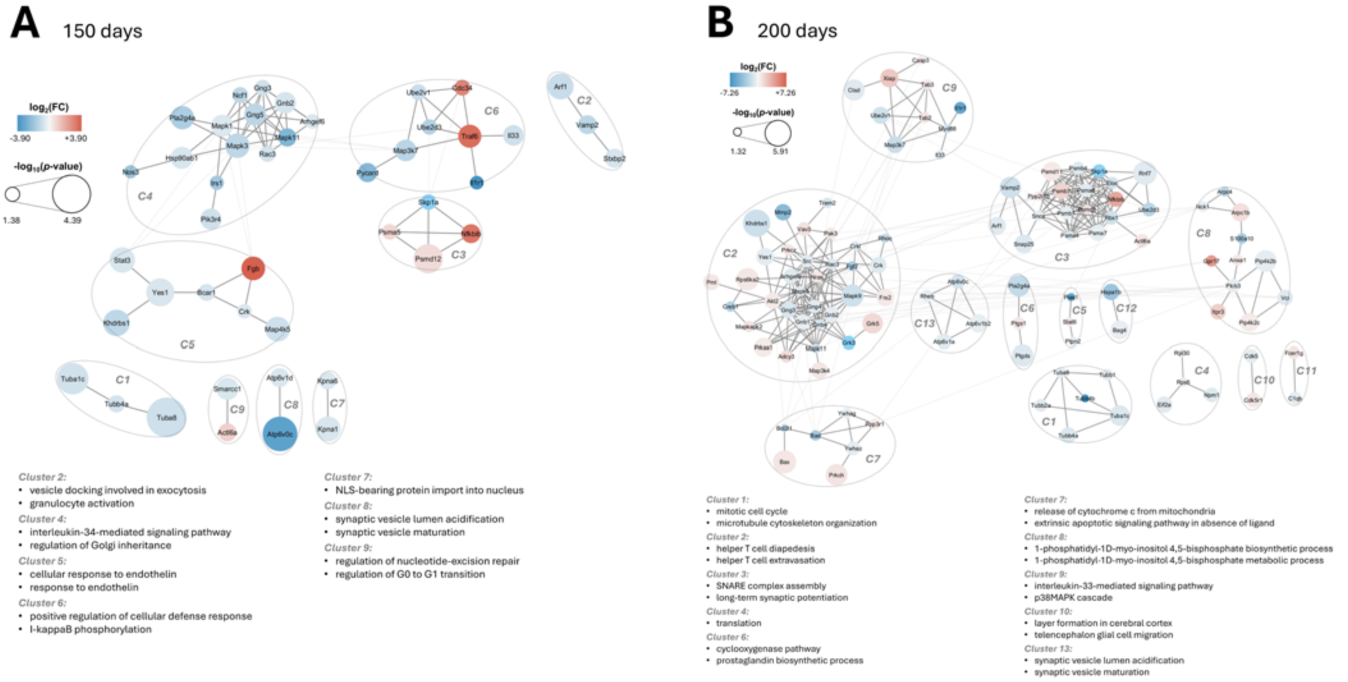


Figure S11: Clustered protein–protein interaction networks of DAPs in APP vs. APP-hA7ko. Clustered protein–protein interaction networks of DAPs identified in the APP vs. APP-hA7ko comparison at 150 (A) and 200 (B) days of age. No PPIs were identified among the DAPs detected at 50 and 100 days. Protein-protein interaction networks were constructed using STRING and visualized in Cytoscape. Node size represents statistical significance ($-\log_{10}(p)$), node fill color indicates the magnitude and direction of protein abundance changes ($\log_2(\mathrm{FC})$), and edges represent predicted functional protein–protein interactions (thicker edges indicate within-cluster interactions, whereas thinner edges indicate between-cluster interactions). Clusters (indicated by ellipses) were identified using the GLay community clustering algorithm and annotated with the top enriched gene ontology (GO) biological process terms. Note that not all clusters showed significant GO term enrichment.

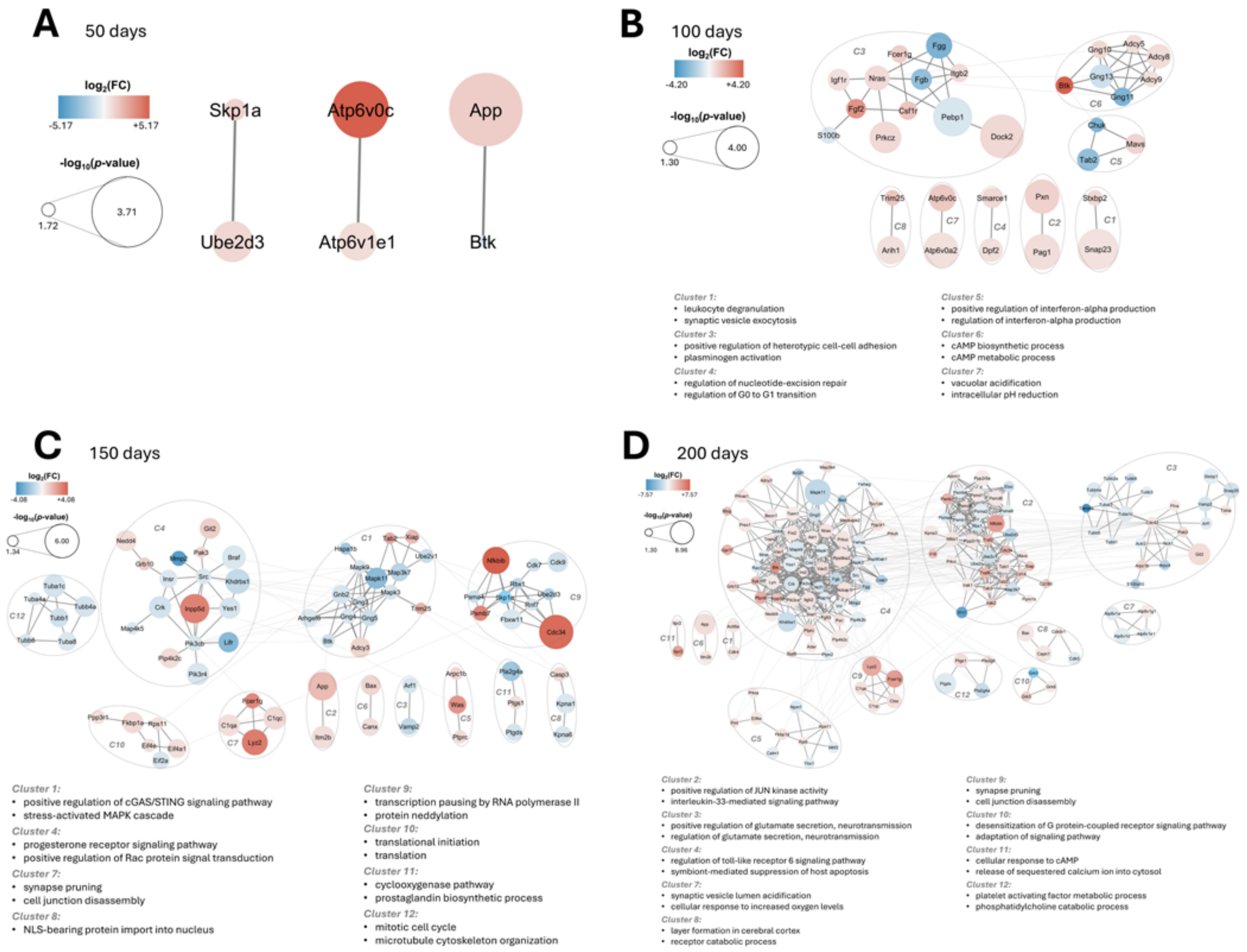


Figure S12: Clustered protein–protein interaction networks of DAPs in B6 vs. APP-hA7ko. Clustered protein–protein interaction networks of DAPs identified in the B6 vs. APP-hA7ko comparison at 50 (A), 100 (B), 150 (C), and 200 (D) days of age. Protein-protein interaction networks were constructed using STRING and visualized in Cytoscape. Node size represents statistical significance ($-\log_{10}(p)$), node fill color indicates the magnitude and direction of protein abundance changes ($\log_2(\mathrm{FC})$), and edges represent predicted functional protein–protein interactions (thicker edges indicate within-cluster interactions, whereas thinner edges indicate between-cluster interactions). Clusters (indicated by ellipses) were identified using the GLay community clustering algorithm and annotated with the top enriched gene ontology (GO) biological process terms. Note that no significant GO term enrichment was observed at 50 days or for some individual clusters because of the low number of DAPs.

**Table S5.** Gene symbols mentioned in the manuscript and their corresponding full names. Symbols are listed alphabetically.

| **Gene symbol** | **Full name** |
|---|---|
| *Abca7* | ATP-binding cassette, subfamily A, member 7 |
| *Adcy5* | adenylate cyclase 5 |
| *Adcy9* | adenylate cyclase 9 |
| *Akt1* | AKT serine/threonine kinase 1 |
| *Akt2* | AKT serine/threonine kinase 2 |
| *Akt3* | AKT serine/threonine kinase 3 |
| *Anxa3* | annexin A3 |
| *Apoe* | apolipoprotein E |
| *App* | amyloid beta precursor protein |
| *Arpc1b* | actin-related protein 2/3 complex, subunit 1B |
| *Atp2b4* | ATPase, calcium transporting, plasma membrane 4 |
| *Atp6v0a2* | ATPase, H+ transporting, lysosomal V0 subunit A2 |
| *Atp6v0c* | ATPase, H+ transporting, lysosomal V0 subunit C |
| *Atp6v1d* | ATPase, H+ transporting, lysosomal V1 subunit D |
| *Atp6v1e1* | ATPase, H+ transporting, lysosomal V1 subunit E1 |
| *Azi2* | 5-azacytidine-induced protein 2 |
| *Bad* | BCL2-associated agonist of cell death |
| *Bax* | BCL2-associated X protein |
| *Bcl2* | B cell leukemia/lymphoma 2 |
| *Braf* | B-Raf proto-oncogene, serine/threonine kinase |
| *Btk* | Bruton agammaglobulinemia tyrosine kinase |
| *C1qa* | complement component 1, q subcomponent, alpha polypeptide |
| *C1qc* | complement component 1, q subcomponent, C chain |
| *Calm1* | calmodulin 1 |
| *Cd14* | CD14 antigen |
| *Cdc34* | cell division cycle 34 |
| *Cdc42* | cell division cycle 42 |
| *Cops3* | COP9 signalosome subunit 3 |
| *Creb1* | cAMP-responsive element-binding protein 1 |
| *Crk* | CRK proto-oncogene, adaptor protein |

**Table S5 (continued).**

| Gene symbol | Full name |
|---|---|
| *Ctsd* | cathepsin D |
| *Dock2* | dedicator of cytokinesis 2 |
| *Dpf2* | double PHD fingers 2 |
| *Eif4a2* | eukaryotic translation initiation factor 4A2 |
| *Eif4e* | eukaryotic translation initiation factor 4E |
| *Eloc* | elongin C |
| *Ephb2* | Eph receptor B2 |
| *Fcer1g* | Fc receptor, IgE, high affinity I, gamma polypeptide |
| *Fga* | fibrinogen alpha chain |
| *Fgb* | fibrinogen beta chain |
| *Fgf2* | fibroblast growth factor 2 |
| *Fgg* | fibrinogen gamma chain |
| *Fn1* | fibronectin 1 |
| *Fyn* | Fyn proto-oncogene, Src family tyrosine kinase |
| *Git2* | G protein-coupled receptor kinase-interacting ArfGAP 2 |
| *Glg1* | Golgi glycoprotein 1 |
| *Gnai1* | guanine nucleotide-binding protein, alpha inhibiting 1 |
| *Gnai3* | guanine nucleotide-binding protein, alpha inhibiting 3 |
| *Gnb1* | guanine nucleotide-binding protein, beta 1 |
| *Gnb2* | guanine nucleotide-binding protein, beta 2 |
| *Gnb4* | guanine nucleotide-binding protein, beta 4 |
| *Gng2* | guanine nucleotide-binding protein, gamma 2 |
| *Gng3* | guanine nucleotide-binding protein, gamma 3 |
| *Gng4* | guanine nucleotide-binding protein, gamma 4 |
| *Gng5* | guanine nucleotide-binding protein, gamma 5 |
| *Gng10* | guanine nucleotide-binding protein, gamma 10 |
| *Gng11* | guanine nucleotide-binding protein, gamma 11 |
| *Gng12* | guanine nucleotide-binding protein, gamma 12 |
| *Gng13* | guanine nucleotide-binding protein, gamma 13 |
| *Grb2* | growth factor receptor-bound protein 2 |

**Table S5 (continued).**

| Gene symbol | Full name |
|---|---|
| *Grb10* | growth factor receptor-bound protein 10 |
| *Grk5* | G protein-coupled receptor kinase 5 |
| *Gyg1* | glycogenin 1 |
| *Hck* | hematopoietic cell kinase |
| *Hdac1* | histone deacetylase 1 |
| *Hdac5* | histone deacetylase 5 |
| *Hmgb1* | high mobility group box 1 |
| *Hspa1b* | heat shock protein 1B |
| *Ighm* | immunoglobulin heavy constant mu |
| *Il1r1* | interleukin 1 receptor, type I |
| *Il1rapl1* | interleukin 1 receptor accessory protein-like 1 |
| *Inpp5d* | inositol polyphosphate-5-phosphatase D |
| *Insr* | insulin receptor |
| *Irak1* | interleukin-1 receptor-associated kinase 1 |
| *Itgam* | integrin alpha M |
| *Itgb2* | integrin beta 2 |
| *Itpr3* | inositol 1,4,5-trisphosphate receptor 3 |
| *Khdrbs1* | KH domain-containing, RNA-binding, signal transduction-associated 1 |
| *Lgmn* | legumain |
| *Lifr* | leukemia inhibitory factor receptor |
| *Lyz2* | lysozyme 2 |
| *Map3k9* | mitogen-activated protein kinase kinase kinase 9 |
| *Map4k4* | mitogen-activated protein kinase kinase kinase kinase 4 |
| *Mapk1* | mitogen-activated protein kinase 1 |
| *Mapk3* | mitogen-activated protein kinase 3 |
| *Mapk9* | mitogen-activated protein kinase 9 |
| *Mapk11* | mitogen-activated protein kinase 11 |
| *Mapk14* | mitogen-activated protein kinase 14 |
| *Nckap1l* | NCK-associated protein 1-like |
| *Nedd4* | neural precursor cell expressed, developmentally downregulated 4 |

**Table S5 (continued).**

| Gene symbol | Full name |
|---|---|
| *Nfkbib* | nuclear factor of kappa light polypeptide gene enhancer in B cells inhibitor, beta |
| *Nlrp3* | NLR family pyrin domain containing 3 |
| *Nos3* | nitric oxide synthase 3, endothelial cell |
| *Npm1* | nucleophosmin 1 |
| *Nras* | neuroblastoma RAS viral oncogene homolog |
| *Pag1* | phosphoprotein associated with glycosphingolipid microdomains 1 |
| *Panx1* | pannexin 1 |
| *Papola* | poly(A) polymerase alpha |
| *Pebp1* | phosphatidylethanolamine-binding protein 1 |
| *Peg3* | paternally expressed 3 |
| *Pfdn2* | prefoldin subunit 2 |
| *Pik3ca* | phosphatidylinositol-4,5-bisphosphate 3-kinase catalytic subunit alpha |
| *Pik3cb* | phosphatidylinositol-4,5-bisphosphate 3-kinase catalytic subunit beta |
| *Pik3r4* | phosphoinositide-3-kinase regulatory subunit 4 |
| *Pip4k2b* | phosphatidylinositol-5-phosphate 4-kinase type 2 beta |
| *Pip4k2c* | phosphatidylinositol-5-phosphate 4-kinase type 2 gamma |
| *Pip5k1b* | phosphatidylinositol-4-phosphate 5-kinase type 1 beta |
| *Pla2g4a* | phospholipase A2, group IVA |
| *Ppp1r13l* | protein phosphatase 1 regulatory subunit 13-like |
| *Ppp2r1b* | protein phosphatase 2 scaffold subunit A beta |
| *Ppp2r5b* | protein phosphatase 2 regulatory subunit B' beta |
| *Ppp3r1* | protein phosphatase 3 regulatory subunit B, alpha |
| *Prkcb* | protein kinase C beta |
| *Prkch* | protein kinase C eta |
| *Prkcz* | protein kinase C zeta |
| *Psen1* | presenilin 1 |
| *Psma6* | proteasome subunit alpha type 6 |
| *Psma7* | proteasome subunit alpha type 7 |
| *Psmb7* | proteasome subunit beta type 7 |
| *Psmd12* | proteasome 26S subunit, non-ATPase 12 |

**Table S5 (continued).**

| Gene symbol | Full name |
|---|---|
| *Pten* | phosphatase and tensin homolog |
| *Ptgds* | prostaglandin D2 synthase |
| *Ptpn6* | protein tyrosine phosphatase, non-receptor type 6 |
| *Ptpn12* | protein tyrosine phosphatase, non-receptor type 12 |
| *Rac1* | Rac family small GTPase 1 |
| *Rac3* | Rac family small GTPase 3 |
| *Rbsn* | rabenosyn, RAB effector |
| *Rbx1* | ring-box 1 |
| *Rela* | RELA proto-oncogene, NF-kappa B subunit |
| *Rnf7* | ring finger protein 7 |
| *Rps6ka2* | ribosomal protein S6 kinase A2 |
| *Rps6kb1* | ribosomal protein S6 kinase B1 |
| *Rps13* | ribosomal protein S13 |
| *S100a1* | S100 calcium-binding protein A1 |
| *S100a9* | S100 calcium-binding protein A9 |
| *S100a10* | S100 calcium-binding protein A10 |
| *Sirpa* | signal-regulatory protein alpha |
| *Skp1* | S-phase kinase-associated protein 1 |
| *Smarcc2* | SWI/SNF-related, matrix-associated, actin-dependent regulator of chromatin subfamily C member 2 |
| *Snap23* | synaptosomal-associated protein 23 |
| *Snap25* | synaptosomal-associated protein 25 |
| *Src* | SRC proto-oncogene, non-receptor tyrosine kinase |
| *Stat6* | signal transducer and activator of transcription 6 |
| *Stxbp1* | syntaxin-binding protein 1 |
| *Stxbp2* | syntaxin-binding protein 2 |
| *Tab1* | TGF-beta-activated kinase 1/MAP3K7-binding protein 1 |
| *Tab2* | TGF-beta-activated kinase 1/MAP3K7-binding protein 2 |
| *Tab3* | TGF-beta-activated kinase 1/MAP3K7-binding protein 3 |
| *Thy1* | thymus cell antigen 1, theta |
| *Traf6* | TNF receptor-associated factor 6 |

**Table S5 (continued).**

| Gene symbol | Full name |
|---|---|
| *Trem2* | triggering receptor expressed on myeloid cells 2 |
| *Tuba1c* | tubulin alpha 1C |
| *Tuba8* | tubulin alpha 8 |
| *Tubb6* | tubulin beta 6 |
| *Ube2d3* | ubiquitin-conjugating enzyme E2 D3 |
| *Ube2v1* | ubiquitin-conjugating enzyme E2 variant 1 |
| *Unc119* | UNC-119 lipid-binding chaperone |
| *Usp11* | ubiquitin-specific peptidase 11 |
| *Vamp2* | vesicle-associated membrane protein 2 |
| *Vcl* | vinculin |
| *Vtn* | vitronectin |
| *Xiap* | X-linked inhibitor of apoptosis |
| *Yes1* | YES proto-oncogene 1, Src family tyrosine kinase |
